# An efficient Bayesian meta-analysis approach for studying cross-phenotype genetic associations with application to Kaiser GERA cohort

**DOI:** 10.1101/101543

**Authors:** Arunabha Majumdar, Tanushree Haldar, Sourabh Bhattacharya, John S. Witte

**Affiliations:** Department of Epidemiology and Biostatistics, University of California San Francisco; Institute for Human Genetics, University of California San Francisco; Interdisciplinary Statistical Research Unit, Indian Statistical Institute, Kolkata

**Author notes:** Address for correspondence:* John S. Witte, Department of Epidemiology and Biostatistics, University of California, San Francisco, 1450 3^rd^ St, Box 3110, San Francisco, CA 94158, *e-mail:.

**Keywords:** Pleiotropy, summary statistics, MCMC, Bayes factor, selection, non-null traits

## Abstract

Simultaneous analysis of genetic associations with multiple phenotypes may reveal shared genetic susceptibility across traits (pleiotropy). For a locus exhibiting overall pleiotropy, it is important to identify which specific traits underlie this association. We propose a Bayesian meta-analysis approach (termed CPBayes) that uses summary-level data across multiple phenotypes to simultaneously measure the evidence of aggregate-level pleiotropic association and estimate an optimal subset of traits associated with the risk locus. This method uses a unified Bayesian statistical framework based on a spike and slab prior. CPBayes performs a fully Bayesian analysis by employing the Markov chain Monte Carlo (MCMC) technique Gibbs sampling. It takes into account heterogeneity in the size and direction of the genetic effects across traits. It can be applied to both cohort data and separate studies of multiple traits having overlapping or non-overlapping subjects. Simulations show that CPBayes produces a substantially better accuracy in the selection of associated traits underlying a pleiotropic signal than the subset-based meta-analysis ASSET. We used CPBayes to undertake a genome-wide pleiotropic association study of 22 traits in the large Kaiser GERA cohort and detected nine independent pleiotropic loci associated with at least two phenotypes. This includes a locus at chromosomal region 1q24.2 which exhibits an association simultaneously with the risk of five different diseases: Dermatophytosis, Hemorrhoids, Iron Deficiency, Osteoporosis, and Peripheral Vascular Disease. The GERA cohort analysis suggests that CPBayes is more powerful than ASSET with respect to detecting independent pleiotropic variants. We provide an R-package ‘CPBayes’ implementing the proposed method.

**Author Summary:** Genome-wide association studies (GWASs) have highlighted shared genetic susceptibility to various human diseases (pleiotropy). We propose a Bayesian meta-analysis method CPBayes that simultaneously evaluates the evidence of aggregate-level pleiotropic association and selects an optimal subset of associated traits underlying a pleiotropic signal. CPBayes analyzes pleiotropy using summary-level data across a wide range of studies for two or more phenotypes - separate GWASs with or without shared subjects, cohort study for multiple traits. It performs a fully Bayesian analysis and offers various flexibilities in the inference. In addition to parameters of primary interest (e.g., the measures of overall pleiotropic association, the optimal subset of associated traits), it provides additional interesting insights into a pleiotropic signal (e.g., the trait-specific posterior probability of association, the credible interval of unknown true genetic effects). Using computer simulations and a real data application to the large Kaiser GERA cohort, we demonstrate that CPBayes offers substantially better accuracy while selecting the non-null traits compared to a well known subset-based meta analysis ASSET. In the GERA cohort analysis, CPBayes detected a larger number of independent pleiotropic variants than ASSET. We provide a user-friendly R-package ‘CPBayes’ for general use.

## Introduction

Genome-wide association studies (GWASs) have detected loci associated with multiple different traits and diseases (i.e., pleiotropy) [Sivakumaran et al., 2011]. For example, pleiotropy has been observed for different types of cancers [Sakoda et al., 2013], immune-mediated diseases [Parkes et al., 2013], and psychiatric disorders [Parkes et al., 2013]. As a specific example, Ellinghaus et al. [2016] demonstrated shared genetic susceptibility to five chronic inflammatory diseases: ankylosing spondylitis, Crohn’s disease, psoriasis, primary sclerosing cholangitis, and ulcerative colitis. Pickrell et al. [2016] systematically compared genetic architecture of 42 phenotypes and reported substantial pleiotropy. Analyzing pleiotropy provides a better understanding of shared pathways and biological mechanisms common to multiple different diseases/phenotypes. From the perspective of clinical genetics, the discovery of a locus simultaneously associated with multiple diseases can support the use of a common therapeutic intervention.

When evaluating a group of phenotypes, one may only expect a subset of them to exhibit pleiotropy. For example, the Global Lipids Genetics Consortium [2013] discovered novel pleiotropic loci associated with different subsets of blood lipid traits. In particular, variants in the genes *RSPO3*, *FTO*, *VEGFA*, *PEPD* were associated with HDL and triglycerides, but not with total cholesterol or LDL. Hence, in addition to evaluating the evidence of overall pleiotropic association, it is crucial to determine which traits are associated with the risk locus to better interpret the pleiotropic signal. Another important consideration is the availability of individual level data from multiple GWASs of different phenotypes. When accessing individual level data is difficult, one can use more readily available genome-wide summary statistics for various phenotypes. In recent years, several methods have been developed to analyze pleiotropy using summary-level data (e.g., Bhattacharjee et al. [2012]; Andreassen et al. [2013a,b]; Chung et al. [2014]; Giambartolomei et al. [2014]; Liley and Wallace [2015]).

Similarly as a GWAS for a single phenotype, a pleiotropic association study can be classified into two main categories – a single nucleotide polymorphism (SNP) is associated with multiple traits, or a genomic region is associated with multiple traits. For example, Bhattacharjee et al. [2012] proposed the subset-based meta-analysis approach ASSET focussing on SNP-level pleiotropy, and Giambartolomei et al. [2014] introduced a Bayesian approach to explore whether two association signals in the same genomic region obtained from two different GWASs share a single causal variant or multiple causal variants.

Andreassen et al. [2013a] proposed a conditional FDR (false discovery rate) approach and detected novel loci associated with Schizophrenia by leveraging information on genetic pleiotropy between Schizophrenia and cardiovascular risk factors. Andreassen et al. [2013b] applied the same approach to study shared genetic architecture underlying two closely related psychiatric disorders Schizophrenia and Bipolar disorder. Liley and Wallace [2015] modified the conditional FDR method to allow for shared controls between two GWASs. Chung et al. [2014] proposed a statistical method GPA to prioritize GWAS signals by incorporating pleiotropy and annotation information. They also demonstrated that GPA performs better than the conditional FDR approach with respect to accurately prioritizing risk SNPs. However, these methods were mainly developed to analyze a pair of traits at a time. On the other hand, ASSET is more directly suited for evaluating pleiotropy simultaneously across two or more number of traits. It provides a p-value evaluating the evidence of aggregate-level pleiotropic association and an optimal subset of associated/non-null traits. It can adjust for correlation between summary statistics. Recent studies (e.g., Ellinghaus et al. [2016]; Carty et al. [2014]; Wang et al. [2014]; Gu et al. [2013]; Kar et al. [2016]) have used ASSET as a primary tool for pleiotropy analysis.

In this article, we focus on the SNP-level pleiotropy and propose a Bayesian meta-analysis approach (termed CPBayes [Cross-Phenotype Bayes]) that simultaneously provides a measure of the evidence of aggregate-level pleiotropic association and an optimal subset of associated traits underlying a pleiotropic signal. The evidence of aggregate-level pleiotropic association is measured by a Bayes factor (BF) and a posterior probability of null association (PPNA). CPBayes explicitly takes into account correlation between summary statistics. For multiple case-control studies of different diseases, the summary statistics across traits become correlated mainly due to sample overlap between studies (e.g., controls). Similarly, for a cohort study, summary statistics across traits are correlated due to assessing multiple phenotypes for the same group of individuals. CPBayes considers heterogeneity both in the size and direction of genetic effects across phenotypes. It also estimates the posterior probability of each phenotype being associated with the risk locus that quantifies the relative contribution of the traits underlying a pleiotropic signal. CPBayes is explicitly designed to analyze two or more number of phenotypes simultaneously.

The Bayesian framework of CPBayes is based on a spike and slab prior, which is commonly used due to its appropriateness and simplicity in solving two-class classification problems [Mitchell and Beauchamp, 1988; George and McCulloch, 1993; Malsiner-Walli and Wagner, 2011; Ishwaran and Rao, 2005; Griffin et al., 2010]. The application of the spike and slab prior in genetic association studies is gradually increasing [Zhou et al., 2013; Wen and Stephens, 2014; Vilhjálmsson et al., 2015]. With a spike and slab prior, the spike element represents a null effect, and the slab component represents a non-null effect. The spike part can either be a positive mass at zero (Dirac spike [Mitchell and Beauchamp, 1988]) or be a normal distribution with mean zero and a small variance (continuous spike [George and McCulloch, 1993]). We design the Gibbs samplers for these two type of prior spikes for both uncorrelated and correlated summary statistics across traits. The continuous spike and slab prior can alternatively be viewed as a special case of the scale mixture of two normal distributions [Efron, 2007, 2012]. Such scale mixtures have been employed to estimate the effect size distributions and replication probabilities in GWAS [Thompson et al., 2015; Holland et al., 2016]. We demonstrate by simulations that the continuous spike offers better accuracy in the selection of associated traits than the Dirac spike (also observed by George and McCulloch [1993] in a general context). The former is also computationally much faster than the latter due to simpler analytic expressions of the full conditional posterior distributions of the model parameters. Hence, we adopted the continuous spike for constructing CPBayes.

First, we contrast CPBayes with GPA for a pair of traits using simulations. Next we explore the performance of CPBayes in various simulation scenarios and compare its efficiency in selecting the non-null traits compared to ASSET and the standard Benjamini-Hochberg (BH) FDR controlling procedure with the level of FDR as 0.01 (denoted as BH_0.01_) [Benjamini and Hochberg, 1995]. The choice of the FDR level is guided by Majumdar et al. [2016] who demonstrated that the simple BH procedure provides better selection accuracy than various different approaches while selecting non-null traits underlying a pleiotropic signal. CPBayes resembles ASSET in that both methods simultaneously draw inference on the evidence of aggregate-level pleiotropic association and on the optimal subset of non-null traits. But, the key advantage of CPBayes is that it selects the non-null traits with substantially higher specificity (proportion of null traits discarded from the optimal subset) than ASSET while maintaining a good level of sensitivity (proportion of associated traits included in the subset).

We also compare CPBayes and ASSET in the analysis of 22 phenotypes in the large Kaiser “Resource for Genetic Epidemiology Research on Adult Health and Aging” (GERA) cohort [dbGaP Study Accession: phs000674.v1.p1]. CPBayes identified nine independent pleiotropic loci associated with at least two phenotypes including a locus at chromosomal region 1q24.2 that exhibits an association with five different diseases: Dermatophytosis, Hemorrhoids, Iron Deficiency, Osteoporosis, and Peripheral Vascular Disease. ASSET identified larger number of independent pleiotropic loci associated with more than one trait, but selected many phenotypes with very weak genetic effects. The GERA cohort analysis also suggests that CP-Bayes is more powerful than ASSET with respect to detecting independent pleiotropic variants. We provide an R-package ‘CPBayes’ implementing the proposed method for a general use by other investigators.

## Material and methods

Let *Y*_1_,…, *Y_K_* denote *K* phenotypes, *G* denote genotype at a single nucleotide polymorphism (SNP), and *W* denote a set of covariates. Suppose, a generalized linear model (GLM) is separately fit for each phenotype as: 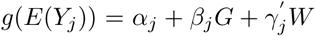, *j* = 1,…, *K*. Let 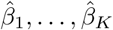 denote the estimates (e.g., maximum likelihood estimates) of ***β***_1_,…, *β_K_* with the corresponding standard errors *s*_1_,…, *s_K_*. Let 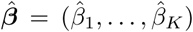, ***β*** = (***β***_1_,…, *β_K_*), and ***s*** = (*s*_1_,…, *s_K_*). Now suppose that we only have the summary statistics (e.g., 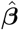 and ***s***). For a large sample size, we can assume that 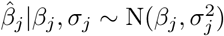. Since *s_j_* is a consistent estimator of *σ_j_*, it is commonly used in place of *σ_j_*. Hence, we assume that 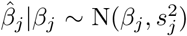. If 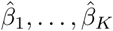 are uncorrelated, 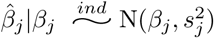; *j* = 1,…, *K*. If 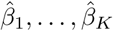 are correlated with a covariance matrix *S*, we assume that 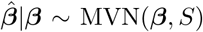. Since the maximum likelihood estimator asymptotically follows a normal distribution and *S* is a consistent estimator of the variance-covariance matrix of 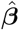, these assumptions should hold well for large sample sizes in contemporary GWASs.

### Continuos spike

The continuous spike and slab prior in our context [George and McCulloch, 1993; Malsiner-Walli and Wagner, 2011] is described as: for *j* = 1,…, *K*,

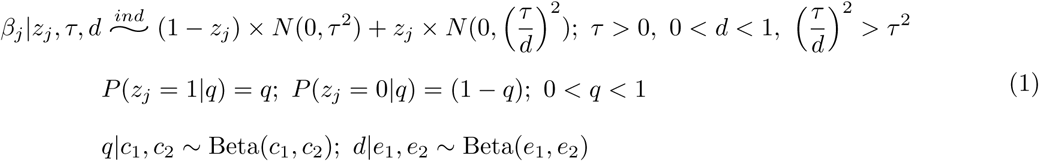

The latent variable *z_j_* denotes the association status of *Y_j_*. When *z_j_* = 0, *β_j_* ∼ *N*(0, *τ*^2^), and when *z_j_* = 1, 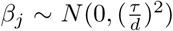, where 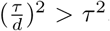. The usefulness of such a formulation is that *τ* can be set small enough so that, if *z_j_* = 0, *|β_j_*| would probably be very small to safely be considered as zero (*Y_j_* is not associated with the SNP), and *d* can be chosen sufficiently small 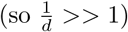 such that, if *z_j_* = 1, *β_j_* can be considered as non-zero (*Y_j_* is associated with the SNP). The proportion of traits having a non-null genetic effect is denoted by *q*. For simplicity and reduction in computational cost, we consider *τ* as fixed. We also choose *c*_1_ = *c*_2_ = *e*_1_ = *e*_2_ = 1 which correspond to the uniform(0, 1) distribution. The parameter *d* is updated in a given range so that the slab variance 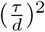 [variance of non-null effects across traits] varies in a pre-fixed range. We describe the continuous spike and slab prior in the context of modeling pleiotropy with diagrams in Figure S1 and S2.

### Dirac spike

The Dirac spike and slab prior in current context [Mitchell and Beauchamp, 1988; Malsiner-Walli and Wagner, 2011] is given by: for *j* = 1,…, *K*,

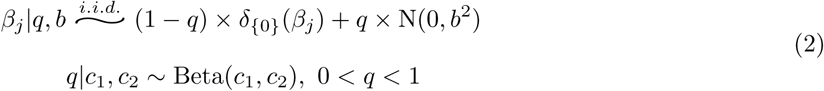

Here, *δ*_{0}_(*β_j_*) = 1 if *β_j_* = 0, and *δ*_{0}_(*β_j_*) = 0 if *β_j_* ≠ 0. So under no association, *β_j_* = 0. The proportion of associated traits is given by *q*. The Dirac spike can be obtained from the continuous spike by first setting *τ* = 0 and 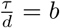 in Equation 1, and then integrating out the latent variables *Z* from the model.

### Statistical inference on pleiotropy by employing MCMC

To perform a fully Bayesian analysis, we implement MCMC by the Gibbs sampling algorithm to generate posterior samples of the model parameters based on which we draw the statistical inference for pleiotropy. We derive the Gibbs samplers for both uncorrelated and correlated summary statistics. Here we describe the inference procedure for the continuous spike. The Gibbs sampling algorithm for the continuous spike (Algorithm 1) is outlined in the Appendix, and the algorithm for the Dirac spike (Algorithm S1) is stated in the supplementary materials. The mathematical derivation of the full conditional posterior distributions underlying the Gibbs samplers are also given in the supplementary materials.

Let {***β***^(*i*)^, *Z*^(*i*)^, *q*^(*i*)^, *d*^(*i*)^; *i* = 1,…, *N*} denote *N* posterior samples of (***β***, *Z*, *q*, *d*) obtained by MCMC after a certain burn-in period. First, we want to test the global null hypothesis of no association (*H*_0_) against the global alternative hypothesis of association with at least one trait (*H*_1_). Since, for the continuous spike, the latent association status distinguishes between an association being null or non-null, we set *H*_0_: *z_1_* =… = *z_K_* = 0 (*Z* = 0) versus *H*_1_: at least one of *z*_1_,…,*z_K_* = 1 (*Z* ≠ 0).

### Bayes factor (BF)

Let *D* denote the summary statistics data at a SNP across traits. The Bayes Factor for testing *H*_1_ against *H*_0_ is given by:

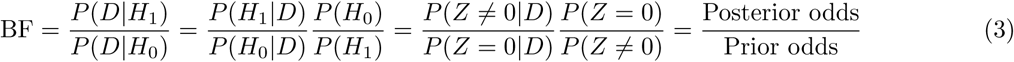

The posterior odds of *H*_1_ vs. 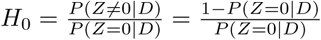, and the prior odds of *H*_1_ vs. 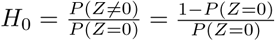 Of note, 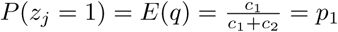. Let *p*_0_ = 1−*p*_1_. Since, *z_j_s* are independently distributed in the prior, 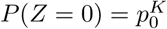 and 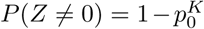. So the prior odds = 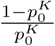. The analytic calculation of *P*(*Z* = 0|*D*) is intractable. Hence, we estimate this conditional probability based on the posterior sample of the model parameters. Note that 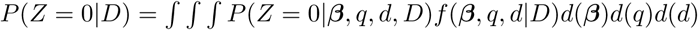. Thus,

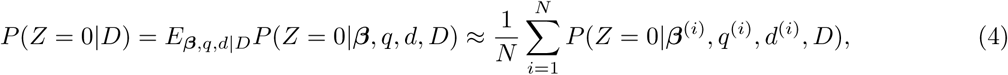
where (***β***^(*i*)^, *q*^(*i*)^, *d*^(*i*)^) denotes the *i^th^* posterior sample of (***β***, *q, d*) obtained by the MCMC. We note that the full conditional posterior distributions of *z*_1_,…, *z_K_* are independent (see step 5 in Algorithm 1 and the derivation of full conditional distributions of *z*_1_,…, *z_K_* in the supplementary materials). Hence, 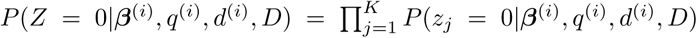. This independence property of the full conditional posterior distributions of *z*_1_,…, *z_K_* is crucial for the explicit estimation of the Bayes factor.

### Posterior probability of null association (PPNA)

We consider another measure for evaluating the aggregate-level pleiotropic association, termed the posterior probability of null association (PPNA). This is based on the posterior probability of association (PPA) introduced in Stephens and Balding [2009]. The posterior odds (PO) of *H*_1_ versus *H*_0_ is 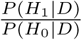. The PPA is given by:

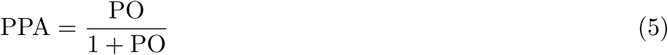

We define PPNA =1 − PPA, which can be viewed as a Bayesian analog of the p-value [Stephens and Balding, 2009]. If the data supports *H*_1_, the PPNA should be close to zero, and if the data supports *H*_0_, it should be close to one (similar to a p-value). The posterior odds is computed in the same way as described above for the Bayes factor. Unlike the Bayes factor, PPNA is not adjusted for the prior odds.

### Selection of optimal subset of associated traits

For *i* = 1,…, *N*, let 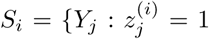 denote the subset of associated traits detected in the *i^th^* MCMC posterior sample. That subset of traits which is observed with the maximum frequency in the posterior sample is estimated as the optimal subset of associated traits. It is the maximum a posteriori (MAP) estimate of the optimal subset.

Let PPA*_j_* denote the marginal trait-specific posterior probability of association which is estimated as 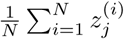 for the phenotype *Y_j_*. PPA*_j_* provides a better insight into a pleiotropic signal. It quantifies the relative contribution of the traits underlying a pleiotropic signal. Even if a trait is not selected in the MAP estimate of the optimal subset of non-null traits, the estimated PPA_*j*_ for the trait may not be negligible, e.g., 25%. An interpretation of such a phenomenon is that even though the estimated genetic effect on a phenotype was not substantial enough to make into the optimal subset, the possibility of the genetic variant having a pleiotropic effect on the trait along with those in the optimal subset seems promising. One can also estimate the joint posterior probability of a particular subset of traits being associated based on the posterior sample of *Z*, e.g., for two traits, it may be of interest to estimate the joint posterior probability that the first trait is associated but the second trait is not [P(*z*_1_ = 1, *z*_2_ = 0|data)]. This probability can be estimated as the proportion of posterior samples in which *z*_1_ = 1 and *z*_2_ = 0.

The direction of association between each non-null trait and a genetic variant can be estimated based on the posterior sample of ***β***. The posterior probability that *Y_j_* is positively associated is estimated as the proportion of positive among *β*_*j*_ the posterior sample of *β*_*j*_. *Y_j_* is classified as being positively associated if this estimated proportion is greater than half. The posterior mean, median, and the 95% credible interval (Bayesian analog of the frequentist confidence interval) of the true genetic effect on each phenotype can be computed based on the posterior sample of ***β***. Of note, we have used a burn-in period of 10000 and MCMC sample size of 10000 while implementing CPBayes in our simulation study and the real data application.

### Specifying the variance of the spike and slab distributions

After extensive experimentation with simulated data, we set the variance of the spike distribution *τ*^2^ (variance of null or very weak genetic effects across traits) to a fixed value 10^−4^. If *β* (log(odds ratio)) follows *N*(0, 10^−4^), then *P*(0.98 < *e^β^* < 1.02) = 0.954. It implies that under the spike (no association), the odds ratio for association between a variant and single trait will vary between 0.98 and 1.02 with a prior probability of 95.4%. In the MCMC, we updated the slab variance 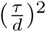 in the range (0.8 − 1.2) with the median value equal to 1. If *β* ∼ *N*(0, 1), then *P*(*e*^*β*^ < 0.95 or *e*^*β*^ > 1.05) = 0.96, which implies that under the slab (association) with variance one, the odds ratio is smaller than 0.95 (a negative association) or larger than 1.05 (a positive association) with a prior probability of 96%. We also explored other choices for these parameters by simulations, such as, *τ*^2^ = 10^−3^, 10^−2^, and 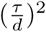 in a range (0.5 − 1.0), (0.7 − 1.1), etc. The values used here 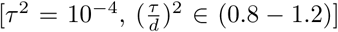 gave an overall high level of specificity while maintaining an overall good level of sensitivity across a wide range of scenarios. The choice of the spike variance and the ratio of the slab and spike variances 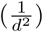 directly impact the selection accuracy [George and McCulloch, 1993]. A smaller choice of the slab variance will increase sensitivity of CPBayes, but at the expense of decreased specificity.

### Estimating the correlation between summary statistics

The summary statistics across traits can be correlated due to overlap or close genetic relatedness among subjects across different studies. For case-control studies, Zaykin and Kozbur [2010] and Lin and Sullivan [2009] derived a simple formula of correlation among 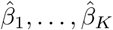. For *k*, *l* ∈ {1,…, *K*} and *k* ≠ l,

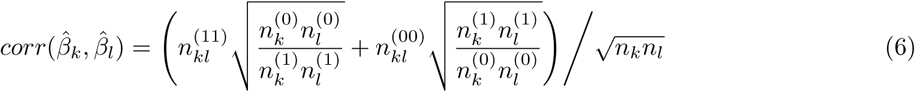

Here 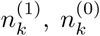 and *n_k_* (or 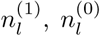, and *n_l_*) denote the number of cases, controls, and total sample size for the study of *Y_k_* (or *Y_l_*); 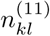 and 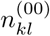 denote the number of cases and controls shared between the studies of *Y_k_* and *Y_l_*. Let 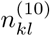 be the number of overlapping subjects that are cases for *Y_k_* but controls for *Y_l_;* similarly, let 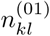 be the number of shared subjects that are controls for *Y_k_* but cases for *Y*_*l*_. Here, the above formula can be generalized to:

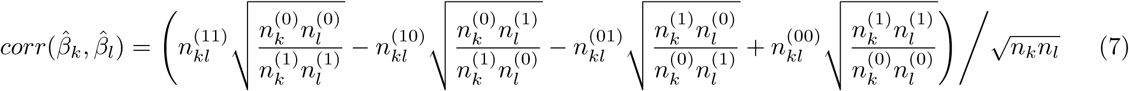

This formula is accurate when none of the phenotypes *Y*_1_,…, *Y_K_* is associated with the SNP or environmental covariates. An alternative strategy [Zhu et al., 2015; Pickrell et al., 2016] is based on using genome-wide summary statistics data to estimate the correlation structure, which is useful when the environmental covariates are associated with the phenotypes or the number of cases or controls shared across studies are not available.

### A combined strategy for correlated summary statistics

For strongly correlated summary statistics, when a majority of the traits are associated with the risk locus (non-sparse scenario), the Gibbs sampler can sometimes be trapped in a local mode rather than the global mode of the posterior distribution due to possible multi-modality of the posterior distribution of model parameters. We observed this pattern in our simulation study. It may result in an incorrect selection of associated traits, reducing the robustness of CPBayes. We noticed that, in such a scenario, if the summary statistics are assumed to be uncorrelated, the MCMC does not get trapped in a local mode and moves around the global mode. But ignoring the correlation can give a lower Bayes factor and sensitivity of the selected traits. Hence, for correlated summary statistics we combine the correlated and the uncorrelated versions of CPBayes as follows. First, we execute CPBayes considering the correlation among 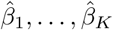 Let A denote the selected subset of non-null traits that contains *K*_1_ traits. Let *B* denote the subset of *K*_1_ traits that have the smallest univariate association p-values. If *A* and *B* match, we accept the results; otherwise, we implement CPBayes assuming that 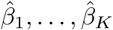 are uncorrelated and take the results obtained. Note that, if *A* is empty, we accept the results obtained by the correlated version of CPBayes.

## Simulation study

We consider multiple case-control studies with or without shared controls [Bhattacharjee et al., 2012]. We also consider a cohort study where the data on multiple disease states are available for a group of individuals [Galesloot et al., 2014; Majumdar et al., 2016]. First, we specify the simulation model and generate the phenotype and genotype data. After computing the summary statistics based on the simulated data, we assume that only the summary-level data are available. For case-control studies with overlapping subjects or a cohort study, we estimate the correlation structure of summary statistics based on the formula given in the Equation 7.

For non-overlapping case-control studies, we consider a separate group of 7000 cases and 10000 controls in each study. For overlapping case-control studies, we consider a distinct set of 7000 cases in each study, and a common set of 10000 controls shared across all the studies. For each disease, we assume an overall disease prevalence of 10% in the whole population. While simulating the genotype data for multiple case-control studies, we assume the standard logistic model of disease probability conditioning on the genotype: logit(*P*(case|*G*)) = *α* + *βG*, where *G* is the genotype at the risk SNP coded as the minor allele count. We simulate the genotypes in cases and controls using the conditional probability of three genotypes conditioning on the case/control status of an individual. In the cohort study, we consider 15000 individuals. First, we generate the continuous traits from a multivariate normal distribution based on the simulation model described in Majumdar et al. [2016] which was adopted from Galesloot et al. [2014]. We choose the trait-specific heritability due to the quantitative trait locus (QTL) at random from (0.2% − 0.5%). Finally, we dichotomize each continuous phenotype into a binary trait subject to an overall disease prevalence of 10%.

We emphasize that the simulation models used here are general in nature and independent of the modeling assumptions underlying CPBayes which are also very general and only require that the sample sizes of the participating GWASs should be sufficiently large to satisfy the standard asymptotic properties. These simulation models were also used in Bhattacharjee et al. [2012], Galesloot et al. [2014], Majumdar et al. [2016], etc.

We contrast CPBayes with GPA for two non-overlapping case-control studies since GPA does not account for shared controls. As the summary statistics are uncorrelated here, we implement the uncorrelated version of CPBayes. We consider 1000 SNPs of which *r*% (*r* = 1, 2) are risk SNPs and (100 − *r*)% are null SNPs (not associated with both traits). SNPs are considered to be in linkage equilibrium (a modeling assumption in GPA). We simulate the minor allele frequency (MAF) at all the SNPs randomly from a Uniform(0.05, 0.5) distribution. For a risk SNP, we consider that one or both of the traits are associated. The odds ratio (OR) for a non-null trait is randomly simulated from Uniform(1.05, 1.25). We assign a larger OR to a risk SNP having a lower MAF and a smaller OR to a risk SNP having a larger MAF. In a given simulation scenario, we implement both the methods and estimate the joint posterior probability of four possible configurations of association with two traits: none of the traits (*p*_00_), 1^st^ trait but not the 2^nd^ trait (*p*_10_), 2^nd^ trait but not the 1^st^ trait (*p*_01_), both the traits (*p*_11_) are associated. Note that, this joint posterior probability is a key component of GPA to evaluate pleiotropy at a particular SNP. We present the estimates of the joint posterior probabilities obtanied by CPBayes and GPA for the risk SNPs and first 10 null SNPs in each simulation scenario in Figure S3 – S12 (see the supplementary material). Interestingly, CPBayes and GPA broadly agree in terms of producing similar estimates of the joint posterior probabilities. We have provided a more detailed summary of this comparison study in the supplementary material.

Next we move to the main simulation study comparing CPBayes and ASSET for two or more case-control studies having overlapping/non-overlapping subjects or multiple case-control phenotypes in a cohort study. Let *K*_1_ denote the number of associated phenotypes among total *K* phenotypes. Suppose, 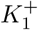 traits are positively associated and 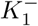 traits are negatively associated 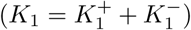. We consider two different choices of MAF at a marker SNP (denoted by *m*): 0.3 and 0.1. Of note, unlike GPA, CPBayes and ASSET analyze each SNP individually. For multiple case-control studies, we consider *K* = 5 (*K*_1_ = 0, 1, 2, 3, 4), *K* =10 (*K*_1_ = 0, 2, 4, 6, 8), and *K* =15 (*K*_1_ = 0, 3, 6, 9). We simulate the odds ratio for a non-null trait at random from Uniform(1.05, 1.25) in each replication under a simulation scenario. We consider the genetic effects across non-null traits to be all positive, and also both positive and negative (obtained by multiplying a positive log odds ratio with −1). For the cohort studies, we considered *K* = 5 (*K*_1_ = 0, 1, 2, 3), *K* =10 (*K*_1_ = 0, 2, 4, 6), and *K* = 15 (*K*_1_ = 0, 3, 6, 9) to save computing time and space.

Note that the Bayes factor and PPNA are not comparable to the p-value. While evaluating the aggregate-level pleiotropic association, we provide various summary measures of log_10_(Bayes factor) (abbreviated as log_10_BF), -log_10_(PPNA) (denoted as -log_10_PPNA), and -log_10_(ASSET p-value) (denoted by -log_10_ASTpv) obtained across 500 replications. Under the global null hypothesis of no association with any trait (*K*_1_ = 0), we describe the summary measures in Table 1 (for non-overlapping case-control studies), 2 (for overlapping case-control studies), and S2 (for a cohort study) provided in the supplementary material. The summary measures in these three tables show that log_10_BF and -log_10_PPNA are very well-controlled under the global null hypothesis. The maximum of log_10_BF in the three tables is observed to be negative (BF < 1). Hence, all the quantiles of log_10_BF also appear to be negative. For example, in Table 1, for *K* = 5 and *m* = 0.3, 75%, 95%, 99% quantiles and the maximum of log_10_BF are -2.68, -2.35, -1.94, and -1.46, respectively. We also observe that 95%, 99% quantiles and the maximum of -log_10_PPNA are smaller than those for -log_10_ASTpv in all the three tables. For example, in Table 2, for *K* = 10 and *m* = 0.1, 95% and 99% quantiles, and the maximum of -log_10_PPNA are 0.07, 0.15, and 1.01, respectively; whereas the same for -log_10_ASTpv are 0.98, 1.91, and 3.42, respectively.

**Table 1:** Simulation study results. Summary of measures of the evidence of the overall pleiotropic association under the global null hypothesis of no association when multiple case-control studies are considered without any overlapping subjects. We considered a separate group of 7000 cases and 10000 controls in each study.

| K | m |  | mean | sd | min | quantiles |  |  |  |  |  |  |
| --- | --- | --- | --- | --- | --- | --- | --- | --- | --- | --- | --- | --- |
|  |  |  |  |  |  | 5% | 25% | 50% | 75% | 95% | 99% | max |
| 5 | 0.3 | $\log_{10}\text{BF}$ | -2.76 | 0.22 | -3.02 | -2.99 | -2.92 | -2.82 | -2.68 | -2.35 | -1.94 | -1.46 |
| | | $-\log_{10}\text{PPNA}$ | 0.03 | 0.02 | 0.01 | 0.01 | 0.02 | 0.02 | 0.03 | 0.06 | 0.13 | 0.32 |
| | | $-\log_{10}\text{ASTpv}$ | 0.48 | 0.47 | 0.00 | 0.01 | 0.13 | 0.33 | 0.68 | 1.45 | 1.97 | 2.88 |
| | 0.1 | $\log_{10}\text{BF}$ | -2.57 | 0.28 | -2.86 | -2.81 | -2.74 | -2.65 | -2.49 | -2.12 | -1.39 | -0.26 |
| | | $-\log_{10}\text{PPNA}$ | 0.05 | 0.08 | 0.02 | 0.02 | 0.02 | 0.03 | 0.04 | 0.09 | 0.35 | 1.25 |
| | | $-\log_{10}\text{ASTpv}$ | 0.43 | 0.44 | 0.00 | 0.005 | 0.11 | 0.30 | 0.60 | 1.24 | 2.14 | 2.63 |
| 10 | 0.3 | $\log_{10}\text{BF}$ | -4.29 | 0.22 | -4.59 | -4.51 | -4.42 | -4.35 | -4.22 | -3.92 | -3.54 | -2.41 |
| | | $-\log_{10}\text{PPNA}$ | 0.03 | 0.04 | 0.01 | 0.01 | 0.02 | 0.02 | 0.03 | 0.05 | 0.11 | 0.70 |
| | | $-\log_{10}\text{ASTpv}$ | 0.32 | 0.42 | 0.00 | 0.00 | 0.04 | 0.17 | 0.44 | 1.14 | 1.82 | 2.99 |
| | 0.1 | $\log_{10}\text{BF}$ | -4.09 | 0.23 | -4.42 | -4.34 | -4.24 | -4.15 | -4.01 | -3.65 | -3.18 | -2.55 |
| | | $-\log_{10}\text{PPNA}$ | 0.04 | 0.04 | 0.02 | 0.02 | 0.02 | 0.03 | 0.04 | 0.09 | 0.22 | 0.59 |
| | | $-\log_{10}\text{ASTpv}$ | 0.30 | 0.38 | 0.00 | 0.00 | 0.04 | 0.17 | 0.44 | 1.05 | 1.66 | 2.53 |
| 15 | 0.3 | $\log_{10}\text{BF}$ | -5.76 | 0.20 | -6.04 | -5.96 | -5.88 | -5.81 | -5.70 | -5.36 | -4.99 | -4.20 |
| | | $-\log_{10}\text{PPNA}$ | 0.03 | 0.03 | 0.01 | 0.02 | 0.02 | 0.02 | 0.03 | 0.06 | 0.12 | 0.48 |
| | | $-\log_{10}\text{ASTpv}$ | 0.24 | 0.38 | 0.00 | 0.00 | 0.001 | 0.08 | 0.34 | 0.89 | 1.42 | 4.37 |
| | 0.1 | $\log_{10}\text{BF}$ | -5.55 | 0.23 | -5.85 | -5.78 | -5.69 | -5.61 | -5.49 | -5.13 | -4.61 | -4.16 |
| | | $-\log_{10}\text{PPNA}$ | 0.05 | 0.04 | 0.02 | 0.02 | 0.03 | 0.03 | 0.04 | 0.09 | 0.25 | 0.51 |
| | | $-\log_{10}\text{ASTpv}$ | 0.24 | 0.41 | 0.00 | 0.00 | 0.00 | 0.08 | 0.32 | 1.05 | 1.78 | 4.40 |

Legend: $K$ - total number of phenotypes, $m$ - allele frequency at the marker SNP; $K = 5, 10, 15$ , and $m = 0.3, 0.1$ . The following abbreviations denote $-\log_{10}\text{BF}$ : $\log_{10}(\text{Bayes factor})$ , $-\log_{10}\text{PPNA}$ : $-\log_{10}(\text{posterior probability of null association})$ , $-\log_{10}\text{ASTpv}$ : $-\log_{10}(\text{ASSET p-value})$ . For multiple studies with no overlapping subjects, the uncorrelated version of CPBayes is implemented. Different summary measures obtained across 500 replications are provided: mean, standard deviation (sd), minimum (min), 5% quantile, 25% quantile, 50% quantile, 75% quantile, 95% quantile, 99% quantile, and maximum (max).

**Table 2:**
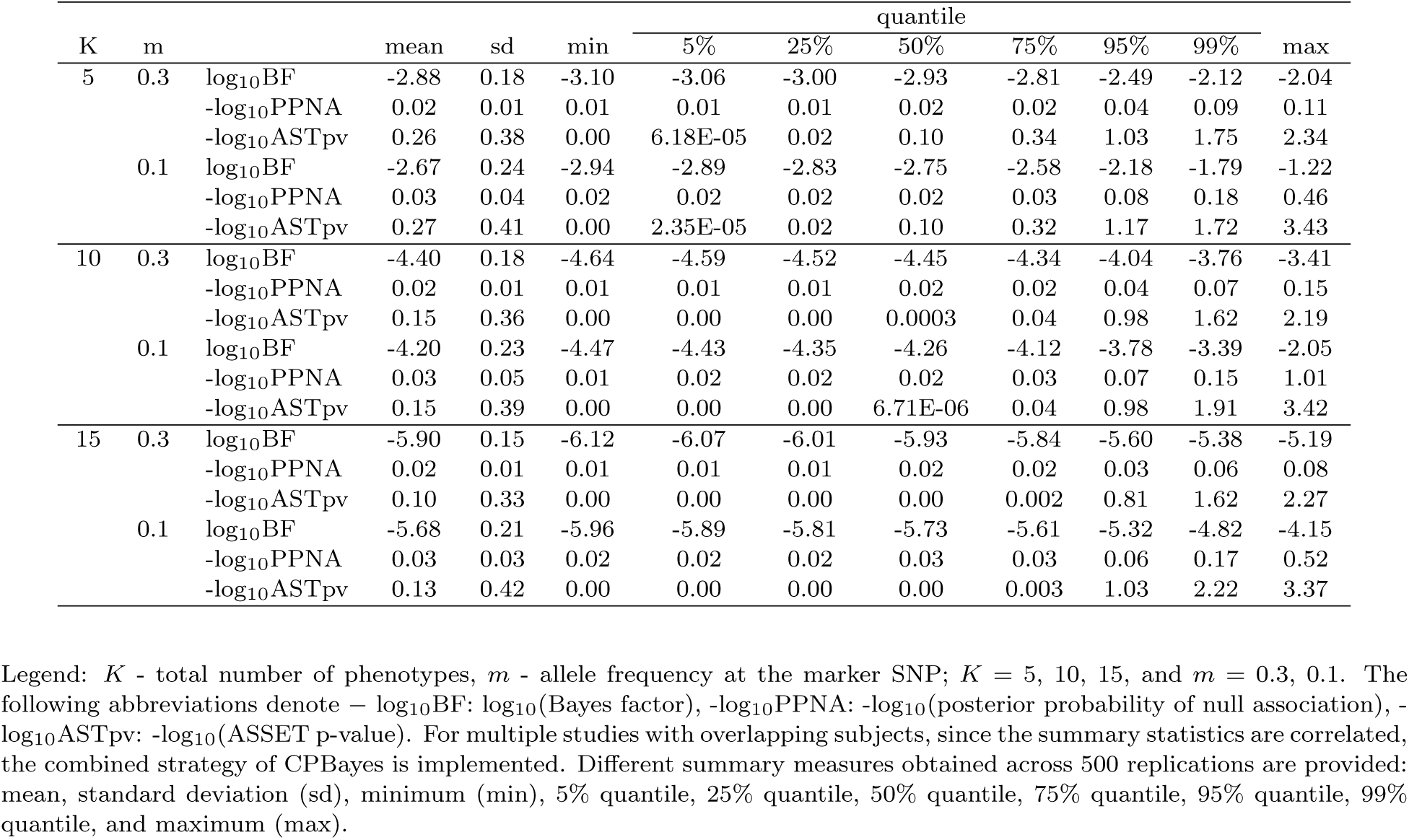
Summary of measures for the evidence of the overall pleiotropic association under the global null hypothesis of no association when multiple case-control studies with overlapping subjects are considered. We considered a distinct set of 7000 cases in each study and a common set of 10000 controls shared across all the studies.

When at least one of the phenotypes is associated (*K*_1_ ≥ 1), we present the summary measures in Table S3, S4, S5, S6, S7 (for non-overlapping case-control studies); S8, S9, S10, S11, S12 (for overlapping case-control studies); and S13, S14, S15 (for cohort study) provided in the supplementary material. As expected, given a choice of *K*, log_10_BF and -log_10_PPNA increase as *K*_1_ increases. For example, in Table S3 (for 5 non-overlapping case-control studies), for *K*_1_ = 1 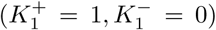 and *m* = 0.3, the mean and median of log_10_BF are 11.05 and 5.12; whereas for *K*_1_ = 2 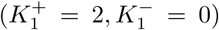 and *m* = 0.3, the mean and median of log10BF are 31.70 and 27.99. Similarly, in Table S10 (for 10 overlapping case-control studies), for *K*_1_ = 2 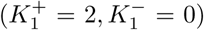 and *m* = 0.1, the mean and median of log_10_BF are 9.70 and 3.30; whereas for *K*_1_ =4 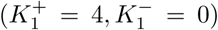 and *m* = 0.1, the mean and median of log_10_BF are 25.97 and 14.82. For overlapping case-control studies and cohort study, for the same choice of *K*_1_, when the non-null effects are both positive and negative, log_10_BF and -log_10_PPNA tend to increase in comparison with when all the non-null effects are positive. For example, in Table S10 (for 10 overlapping case-control studies), for 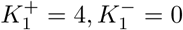 (*K*_1_ = 4) and *m* = 0.3, the mean and median of log_10_BF are 57.66 and 53.68; whereas for 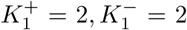 (*K*_1_ = 4) and *m* = 0.3, the mean and median of log_10_BF are 77.81 and 72.52. Similarly, in Table S14 (for binary cohort with 10 phenotypes), 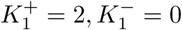 (*K*_1_ = 2) and *m* = 0.3, the mean and median of log_10_BF are 15.43 and 6.94; whereas for 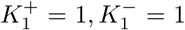 (*K*_1_ = 2) and *m* = 0.3, the mean and median of log_10_BF are 25.82 and 19.64. However, for non-overlapping case-control studies, we do not observe such a trend with respect to the direction of the non-null effects. We also observe that -log_10_ASTpv behaves similarly to log_10_BF and -log_10_PPNA.

In Figure 1, we have plotted log_10_BF and -log_10_PPNA for 100 replications in the same set-up of five non-overlapping and overlapping case-control studies considered above. We chose MAF = 0.3. As expected, when none of the phenotypes is associated, log_10_BF cluster below zero and -log_10_PPNA cluster close to zero (Figure 1). And, the measures tend to increase as the number of associated traits increases (Figure 1).

**Figure 1:**
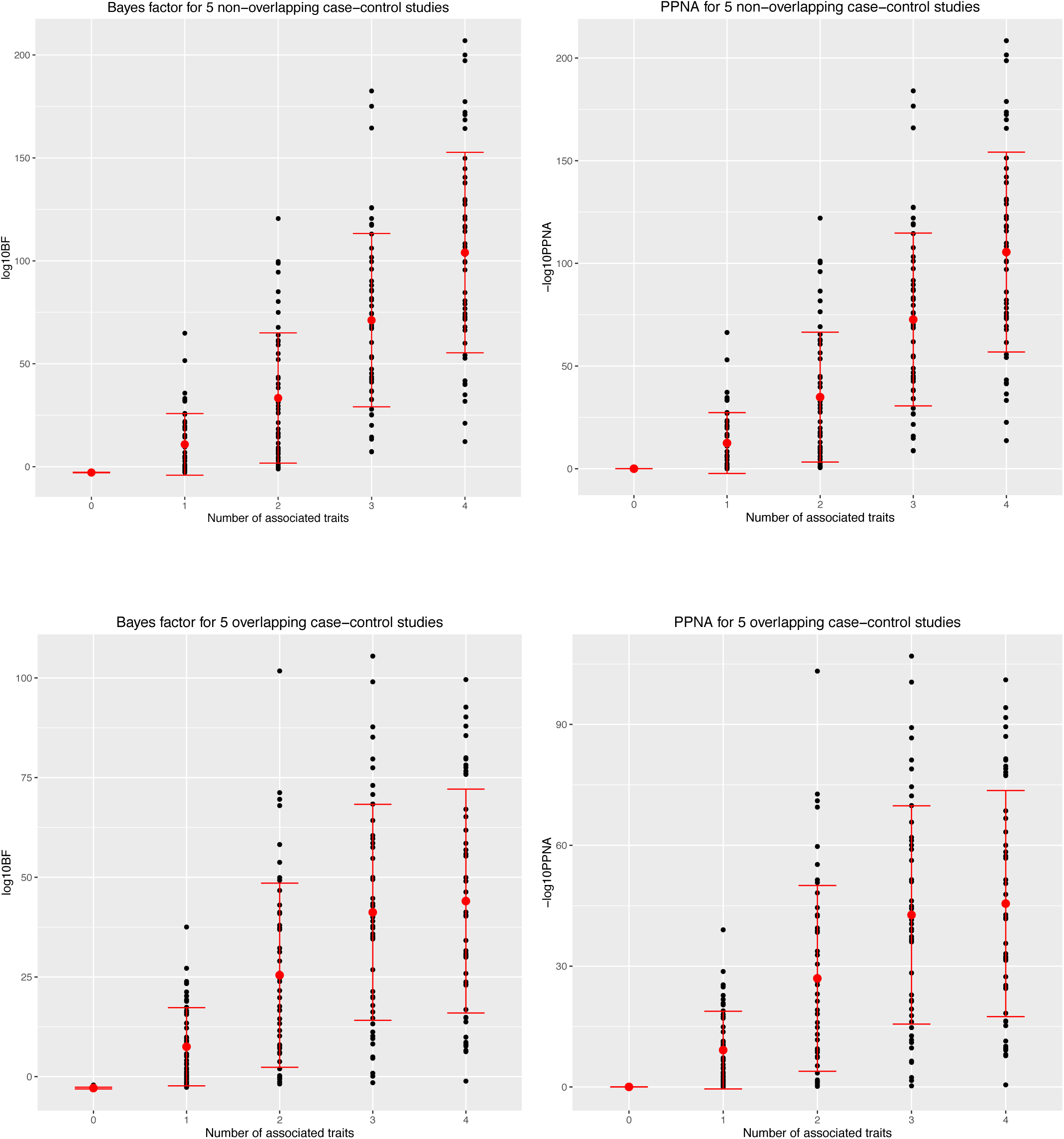
Simulation study results. Bayes factor and PPNA for five non-overlapping and overlapping case-control studies across 100 replications. The red colored band presents (mean – standard deviation, mean + standard deviation) of the Bayes factor or PPNA across the replications.

For overlapping case-control studies and cohort study, we implemented the combined strategy of CPBayes. For each simulation scenario in Table S8, S9, S10, S11, S12 (for overlapping case-control studies), and S13, S14, S15 (for a cohort study), we provide the percentage of replications in which the combined strategy chose the uncorrelated version (denoted by uncor%). From the tables, we observe that uncor% increases as *K*_1_ increases for a given choice of *K*. For example, in Table S11 (for 10 overlapping case control studies), for m = 0.3, uncor% increases from 3% to 6.4% as *K*_1_ increases from 6 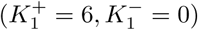 to 8 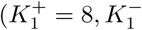 = 0). Similarly, in Table S15 (for a cohort study with 15 phenotypes), for *m* = 0.1, uncor% increases from 5.8% to 8.4% as *K*_1_ increases from 6 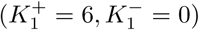 to 9 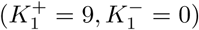. For a sparse scenario when less than half of the traits are associated (e.g., *K*_1_ = 3 and *K* = 15), uncor% is substantially smaller than that for a non-sparse scenario (e.g., *K*_1_ = 9 and *K* = 15). For example, in Table S12 (for 15 overlapping case-control studies), for *m* = 0.1, uncor% is 11% when *K*_1_ =3 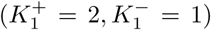, and it increases to 26.2% when *K*_1_ = 9 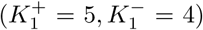. Similarly, in Table S15 (for a cohort study with 15 phenotypes), for *m* = 0.3, uncor% increases from 4.8% to 7.6% when *K*_1_ increases from 3 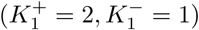 to 9 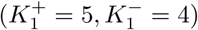.

To evaluate the accuracy of selection of associated traits in a simulation scenario, we computed the average specificity and sensitivity obtained across 500 replications. We present the comparison of the selection accuracy between CPBayes, ASSET, and BH_0.01_ in Figure 2 (for non-overlapping case-control studies), Figure 3(for overlapping case-control studies), and Figure S13 (for cohort study). First we note that CPBayes and BH_0.01_ offer similar selection accuracy. In Figure 2, 3, and S13, points plotting the average specificity and sensitivity for CPBayes and BH_0.01_ across various simulation scenarios cluster around each other. So next, we focus on comparing the selection accuracies between the two main competing methods CPBayes and ASSET.

**Figure 2:**
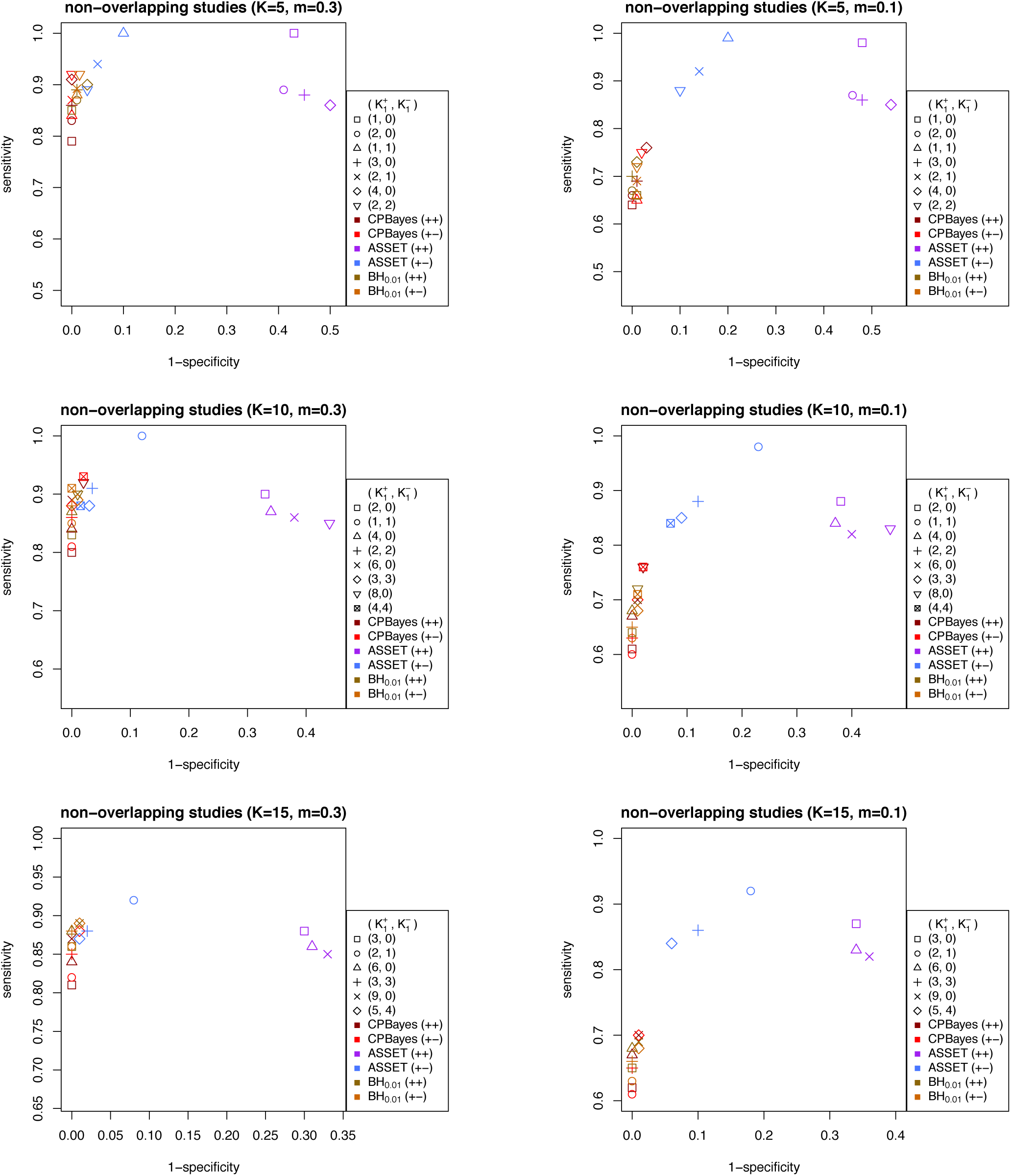
Simulation study results. Comparison of the accuracy of selection of associated traits by CPBayes and ASSET for multiple non-overlapping case-control studies. The total number of phenotypes/studies is denoted by *K* and *m* denotes the minor allele frequency at the risk SNP.

**Figure 3:**
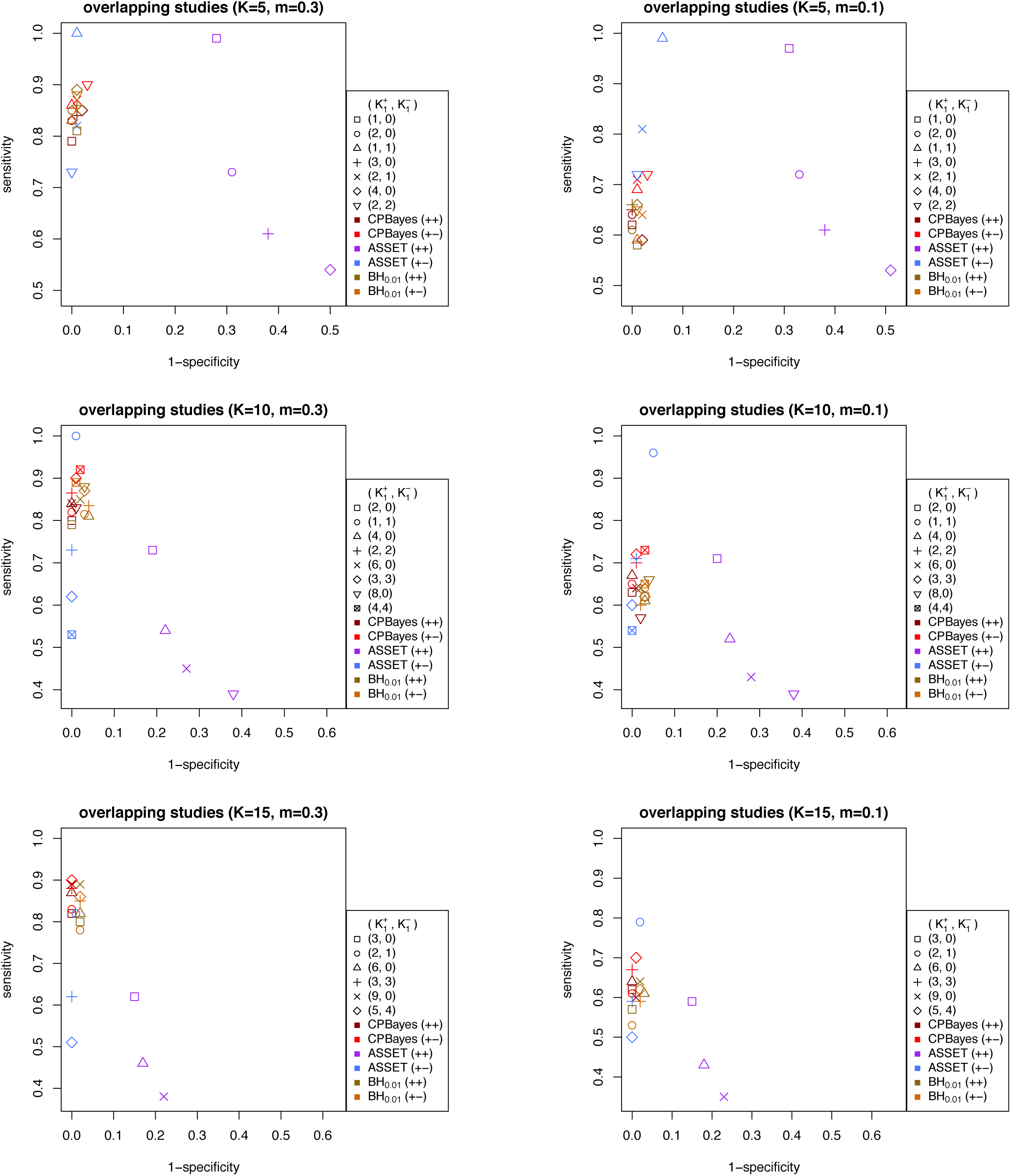
Comparison of the accuracy of selection of associated traits by CPBayes and ASSET for multiple overlapping case-control studies. The total number of phenotypes/studies is denoted by *K* and *m* denotes the minor allele frequency at the risk SNP.

CPBayes yielded a very good level of specificity (consistently more than 95%) which is substantially higher than that of ASSET. For example, in Figure 2 (for non-overlapping case-control studies), while CPBayes’ specificity is > 97%, the specificity of ASSET varies in the range 46% − 98% when *K* = 5, and in a range 53% − 99% when *K* = 10. Similary, in Figure 3 (for overlapping case-control studies), the specificity of ASSET varies in a range 49% − 99% when *K* = 5, and in 62% − 100% when *K* = 10.

CPBayes also offers an overall good level of sensitivity. For five non-overlapping case-control studies (Figure 2), CPBayes produced a sensitivity of 79% − 89% when MAF = 0.3 and 64% − 76% when MAF = 0.1; ASSET yielded slightly higher sensitivity of 86% − 100% (with 50% − 98% specificity) when MAF = 0.3, and a higher sensitivity of 85% − 98% (with 46% − 90% specificity) when MAF = 0.1. In Figure 2, we observe a similar pattern for *K* = 10 and *K* = 15. ASSET’s higher sensitivity than CPBayes appears to come at the expense of a substantially lower specificity.

For overlapping case-control studies (Figure 3), CPBayes gave a substantially better sensitivity than ASSET (along with substantially better specificity) for a majority of the simulation scenarios. For example, when *K* =10 and *m* = 0.3, the sensitivity was 82% − 89% for CPBayes, and 39% − 73% for ASSET, except for the case 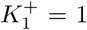, 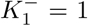 when ASSET had 100% sensitivity. Similarly, for *K* =10 and *m* = 0.1, the sensitivity was 57% − 73% for CPBayes and 39% − 71% for ASSET except for the choice 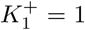, 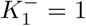 when ASSET had 96% sensitivity. For *K* = 15 and *m* = 0.3, the sensitivity was 82% − 87% for CPBayes and 38% − 82% for ASSET. For *K* =15 and *m* = 0.1, the sensitivity was 60% − 70% for CPBayes and 35% − 79% for ASSET.

For a cohort study, CPBayes and ASSET both consistently exhibited good levels of sensitivity. For some cases, ASSET had a higher sensitivity than CPBayes, though generally with lower specificity. We also observe that when the non-null effects are all positive, the specificity of ASSET is smaller compared to when the non-null effects are both positive and negative. However, CPBayes performed more robustly with respect to the direction of non-null effects.

We also carried out simulations for 50 traits. We considered the same set-up of non-overlapping and overlapping case-control studies considered above and *K*_1_ = 0, 5, 10. Since ASSET is computationally very slow for 50 traits due to an extremely large number of possible subsets of traits, we only implemented CPBayes. We also applied BH_0.01_ to select the non-null traits. Different summary measures of log_10_BF and -log_10_PPNA obtained across 200 replications are described in Table S16 for non-overlapping studies and in Table S17 for overlapping studies. The mean specificity and sensitivity of CPBayes and BH_0.01_ are provided in Table S18 for non-overlapping studies and Table S19 for overlapping studies. We observe that CPBayes performs similarly as for 5, 10, or 15 non-overlapping or overlapping studies (discussed above). While selecting the associated traits, both CPBayes and BH_0.01_ produced very high level of specificity. For non-overlapping studies, BH_0.01_ produced marginally higher sensitivity than CPBayes (Table S18). However, for overlapping studies, CPBayes produced higher sensitivity than BH_0.01_ (Table S19), in particular for lower minor allele frequency (MAF = 0.01).

## GERA cohort analysis

To investigate the performance of CPBayes using real data, we analyzed the large genome-wide association study from the Kaiser “Resource for Genetic Epidemiology Research on Adult Health and Aging” (GERA) cohort data obtained from dbGaP [dbGaP Study Accession: phs000674.v1.p1]. We also analyzed the data by ASSET. For simplicity’s sake, we restricted our analysis to 62,318 European-American individuals, who constitute more than 75% of the dbGaP data. We tested 657,184 SNPs genotyped across 22 autosomal chromosomes for their potential pleiotropic effects on 22 phenotypes in the GERA cohort (Table S20). Note that in the dbGaP data, the cancers are collapsed into a single variable (any cancer). Therefore, we could only use an overall cancer categorization even though the genetic architecture is likely heterogeneous across different cancers. We provide the trait-trait correlation matrix in Table S26. The phenotypes are correlated modestly with a maximum correlation of 0.36 observed between Hypertension and Dyslipidemia.

Before our analysis, we undertook the following QC steps. First, we removed individuals with: over 3% of genotypes missing; any missing information on covariates (described below); genotype heterozygosity outside six standard deviations; first degree relatives; or discordant sex information. This left us with 53,809 individuals. Next, we removed SNPs with: MAF < 0.01; 10% or more missingness; or deviation from HWE at a level of significance 10^−5^. This leaves 601,175 SNPs that were tested for pleiotropic association by CPBayes and ASSET. We adjusted the analysis for the following covariates: age, gender, smoking status, BMI category, and 10 principal components of ancestry (PCs). We tested the single-trait association for each of the 22 phenotypes by a logistic regression of the case-control status on the genotype incorporating the same set of adjusting covariates. We used SNP-trait effect estimates (log odds ratios) and their standard errors in CPBayes and ASSET. Since the summary statistics are correlated, we used the combined strategy of CPBayes and the correlated version of ASSET.

As we have environmental covariates in the GERA study, we estimated the correlation matrix of the effect estimates using the genome-wide summary statistics data [Zhu et al., 2015]. First, we extracted all of the SNPs for each of which the trait-specific univariate p-value across 22 traits are > 0.1. This ensures that each SNP is either weakly or not associated with any of the 22 phenotypes. Then we selected a set of 24,510 independent SNPs from the initial set of null SNPs by using a threshold of *r*^2^ < 0.01 (*r*: the correlation between the genotypes at a pair of SNPs). Finally, we computed the correlation matrix of the effect estimates as the sample correlation matrix of 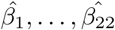 across the selected 24,510 independent null SNPs.

We apply the conventional genome-wide (GW) level of statistical significance 5 × 10^−8^ for ASSET (equivalent to -log_10_(ASSET p-value) > 7.30). It is tough to decide on an appropriate GW cut-off for log_10_BF of CPBayes. Of note, Bhattacharjee et al. [2012] and Majumdar et al. [2016] demonstrated by simulations that ASSET appropriately controls for the false positive rate. We observed in our simulation study that, under the global null hypothesis of no association, CPBayes always produced a negative log_10_BF (Table 1, 2, S2). Hence log_10_BF was always smaller than -log_10_AST_pv_ under the null hypothesis. So, for contrasting the results obtained by the two methods, we set the cut-off of log_10_BF as 7.30, the same as for -log_10_AST_pv_. This cut-off may be somewhat conservative for CPBayes, because in the simulation study, CPBayes produced substantially smaller log_10_BF than -log_10_AST_pv_ under the global null hypothesis (Table 1, 2, S2). Based on the GW cut-off, CPBayes identified 314 SNPs and ASSET detected 394 SNPs. By definition, each of these SNPs is associated with at least one of the 22 phenotypes. We note that CPBayes and ASSET identified a common set of 253 SNPs.

Many of the associated SNPs are expected to be in linkage disequilibrium (LD). On each chromosome, we identified the LD blocks by using a threshold of *r^2^* = 0.25. For CPBayes, we identified 49 associated LD blocks, and for ASSET, we detected 30 associated LD blocks. For each of the 394 SNPs detected by ASSET, the optimal subset of non-null traits always included more than one phenotype. So for ASSET, within each LD block, we chose the SNP having the minimum p-value of aggregate-level pleiotropic association. We present the results for these lead SNPs in Table S22, S23, S24, and S25. CPBayes selected more than one trait for 63 among 314 SNPs. Within each LD block identified by CPBayes, we chose the SNP associated with the maximum number of phenotypes. If multiple SNPs satisfy this criterion, we chose the one having the maximum log_10_BF. If every SNP in a block is associated with one trait, we chose the SNP with the maximum log_10_BF. In Table 3, we present the results for the independent pleiotropic SNPs at which CPBayes selected at least two phenotypes. In Table S21, we report the selected independent SNPs at which CPBayes detected one trait. In the tables for CPBayes, we present the estimated trait-specific posterior probability of association (PPA_*j*_) and the direction of association for the selected phenotypes (genotype was coded as the number of wild allele). In all the tables for CPBayes and ASSET, we also provide the trait-specific univariate association p-values. For the independent pleiotropic SNPs identified by both the methods, we applied BH_0.01_ and list the selected traits in Table 3, S21, S22, S23, S24 and S25 for contrast’s sake.

**Table 3:** Independent pleiotropic SNPs detected by CPBayes which are associated with at least two phenotypes.

| rsID | chrom<br>band | CPBayes<br>log<sub>10</sub>BF | CPBayes<br>PPNA | Subset of associated<br>traits selected by CPBayes | PPA<sub>j</sub> | Direction | Univariate<br>p-values | Subset of associated<br>traits selected by BH<sub>0.01</sub> |
| --- | --- | --- | --- | --- | --- | --- | --- | --- |
| rs6025 | 1q24.2 | 238.09 | 1.92E-245 | Dermatophytosis | 63.8% | positive | 0.0018 | Dermatophytosis |
|  |  |  |  | Hemorrhoids | 68.7% | positive | 0.0014 | Hemorrhoids |
|  |  |  |  | Iron Deficiency | 90.2% | positive | 0.0004 | Iron Deficiency |
|  |  |  |  | Osteoporosis | 91.9% | negative | 0.0002 | Osteoporosis |
|  |  |  |  | Peripheral Vascular Disease | 100% | negative | 6.81E-14 | Peripheral Vascular Disease |
| rs7601401 | 2p16.1 | 9.69 | 4.91E-17 | Abdominal Hernia | 100% | positive | 3.88E-12 | Abdominal Hernia |
|  |  |  |  | Osteoarthritis | 86.7% | positive | 3.46E-06 | Osteoarthritis |
| rs17661572 | 2p21 | 8.47 | 7.62E-16 | Allergic Rhinitis | 94.1% | negative | 4.32E-05 | Allergic Rhinitis |
|  |  |  |  | Insomnia | 98.4% | positive | 0.0001 | Insomnia |
| rs387608 | 6p21.33 | 29.06 | 2.07E-36 | Cancers | 54.6% | positive | 0.0001 | Cancers |
|  |  |  |  | Macular Degeneration | 100% | positive | 4.29E-12 | Macular Degeneration |
| rs10455872 | 6q25.3 | 26.20 | 1.49E-33 | Cardiovascular Disease | 72.9% | negative | 6.14E-05 | Cardiovascular Disease |
|  |  |  |  | Dyslipidemia | 100% | negative | 6.97E-15 | Dyslipidemia |
|  |  |  |  | Peripheral Vascular Disease | 66.8% | negative | 0.0002 | Peripheral Vascular Disease |
| rs3830123 | 6p21.32 | 22.31 | 1.16E-29 | Cancers | 99.9% | negative | 1.01E-07 | Cancers |
|  |  |  |  | Dyslipidemia | 100% | negative | 5.55E-13 | Dyslipidemia |
| rs9275476 | 6p21.32 | 22.10 | 1.88E-29 | Asthma | 100% | negative | 5.61E-08 | Asthma |
|  |  |  |  | Type 2 Diabetes | 100% | negative | 4.27E-06 | Type 2 Diabetes |
|  |  |  |  | Macular Degeneration | 56% | positive | 0.0011 | Macular Degeneration |
| rs651007 | 9q34.2 | 24.72 | 4.57E-32 | Dyslipidemia | 100% | negative | 2.89E-15 | Dyslipidemia |
|  |  |  |  | Peripheral Vascular Disease | 100% | negative | 9.55E-08 | Peripheral Vascular Disease |
| rs4506565 | 10q25.2 | 117.08 | 1.96E-124 | Type 2 Diabetes | 100% | negative | 2.02E-55 | Type 2 Diabetes |
|  |  |  |  | Dyslipidemia | 60.6% | negative | 1.33E-06 | Dyslipidemia |

Legend: The chromosome band of a SNP is denoted by ‘chrom band’. Direction means whether the SNP is positively or negatively associated with the phenotype.

Even though CPBayes detected a smaller number of GW-significant SNPs (314) than ASSET (394), the former identified a substantially larger number of LD blocks than the later (49 versus 30). For example, CPBayes identified one SNP on chromosome 18 associated with Peripheral Vascular Disease, but ASSET did not detect any SNP on this chromosome. Specifically, CPBayes detected rs8092654 (log_10_BF = 12.24) at which ASSET yielded a p-value of 0.13. In the NHGRI-EBI GWAS catalog, rs8092654 is reported as an eQTL hit for the ZNF611 gene in the peripheral blood monocytes tissue. As another example, on chromosome 11, CPBayes detected three LD blocks. From Table S21 for CPBayes, we observe that the lead SNPs of the blocks are rs1799963 (11p11.2), rs964184 (11q23.3), and rs55975204 (11q13.2) which are associated with Peripheral Vascular Disease (univariate p-value: 1.19 × 10^−8^), Dyslipidemia (univariate *p*-value: 5.49 × 10^−28^), and Osteoporosis (univariate p-value: 2.0 × 10^−9^), respectively. But, ASSET detected only one LD block on chromosome 11, which contains SNPs that are mainly associated with Dyslipidemia. In this block, the lead SNP for ASSET (rs964184) also turned out to be the lead SNP for CPBayes in the corresponding LD block containing the SNPs associated with Dyslipidemia. So, ASSET missed the signals for Peripheral Vascular Disease and Osteoporosis. We note that, in the NHGRI-EBI GWAS catalog, rs1799963 is reported to be associated with venous thromboembolism, and rs964184 is reported as associated with LDL cholesterol, triglycerides, and total cholesterol. And, rs55975204 is in LD (*r^2^* = 0.9) with rs12286536 which is an eQTL hit for the CPT1A and MTL5 genes in the whole blood tissue. These findings suggest that in certain situations CPBayes may detect associations not detected by ASSET. We also present the Manhattan plot for CPBayes in Figure S15 and ASSET in Figure S16 which clearly show that CPBayes detected larger number of pleiotropic regions than ASSET.

For each SNP detected by ASSET, the selected subset of non-null traits always contained more than one phenotype (Table S22, S23, S24, S25). For a majority of the SNPs, the subset of non-null traits included many phenotypes that have large univariate association p-values (so, weak effects). For example, rs2300430 (1q31.3) was detected by both the methods (first SNP in Table S22 and S21); ASSET selected Allergic Rhinitis, Depressive Disorder, Dermatophytosis, Hemorrhoids, Insomnia, Macular Degeneration, Osteoporosis, and Peptic Ulcer, which have trait-specific univariate p-values equal to: 0.34, 0.18, 0.09, 0.36, 0.47, 2.57×10^−77^, 0.09, and 0.74, respectively. In contrast, CPBayes only selected Macular Degeneration (Table S22). This suggests that CPBayes selects only those phenotypes having substantially strong genetic effects, while ASSET may select many more traits with lower specificity as seen in our simulation study. BH_0.01_ only selected Macular Degeneration which is consistent with CPBayes. Among 30 independent pleiotropic SNPs detected by ASSET, BH_0.1_ selected more than one trait only for nine SNPs, whereas ASSET selected multiple traits for each of them (Table S22, S23, S24, S25). This again indicates lower specificity of ASSET.

CPBayes detected nine independent GW significant pleiotropic SNPs, for which it selected at least two phenotypes as non-null (Table 3). For example, at rs6025 (1q24.2), it selected a maximum of 5 phenotypes: Dermatophytosis, Hemorrhoids, Iron Deficiency, Osteoporosis, and Peripheral Vascular Disease, which have univariate p-values equal to 0.0018, 0.0014, 0.0004, 0.0002, and 6.81 × 10^−14^, respectively (Figure S17). ASSET also identified this SNP and selected the same five traits as CPBayes. Interestingly, the SNP was positively associated with Dermatophytosis, Hemorrhoids, and Iron Deficiency, but negatively associated with Osteoporosis and Peripheral Vascular Disease (Figure S17). At rs10455872 (6q25.3), CPBayes selected Cardiovascular Disease, Dyslipidemia, and Peripheral Vascular Disease (Figure S21). ASSET selected these three traits and five more phenotypes with large univariate p-values (Figure S21). At rs651007 (9q34.2), CPBayes detected pleiotropy with Dyslipidemia and Peripheral Vascular Disease, whereas ASSET detected both of these and other phenotypes with weak genetic effects (Figure S24). CPBayes detected two independent pleiotropic SNPs in the chromosomal region 6p21.32: rs3830123 and rs9275476 (Table 3). The *r*^2^ value between rs3830123 and rs9275476 was 0.016. We also found that a disjoint set of phenotypes are associated with the two SNPs: Asthma, Type 2 Diabetes, and Macular Degeneration are associated with rs9275476; Cancers and Dyslipidemia are associated with rs3830123. CPBayes also detected four other pleiotropic loci: rs7601401 (2p16.1) [Abdominal Hernia and Osteoarthritis], rs17661572 (2p21) [Allergic Rhinitis and Insomnia], rs387608 (6p21.33) [Cancers and Macular Degeneration], and rs4506565 (10q25.2) [Type 2 Diabetes and Dyslipidemia]. For these nine independent pleiotropic SNPs identified by CPBayes, BH_0.01_ selected the same subset of traits as CPBayes (Table 3), which is expected because the two approaches offered similar selection accuracy in our simulation study. We present a forest plot for each independent pleiotropic signal detected by CPBayes in Figure S17 – S25. We also provide a circular plot in Figure 7 presenting 22 pair-wise trait-trait pleiotropic association signals detected by CPBayes.

**Figure 4:**
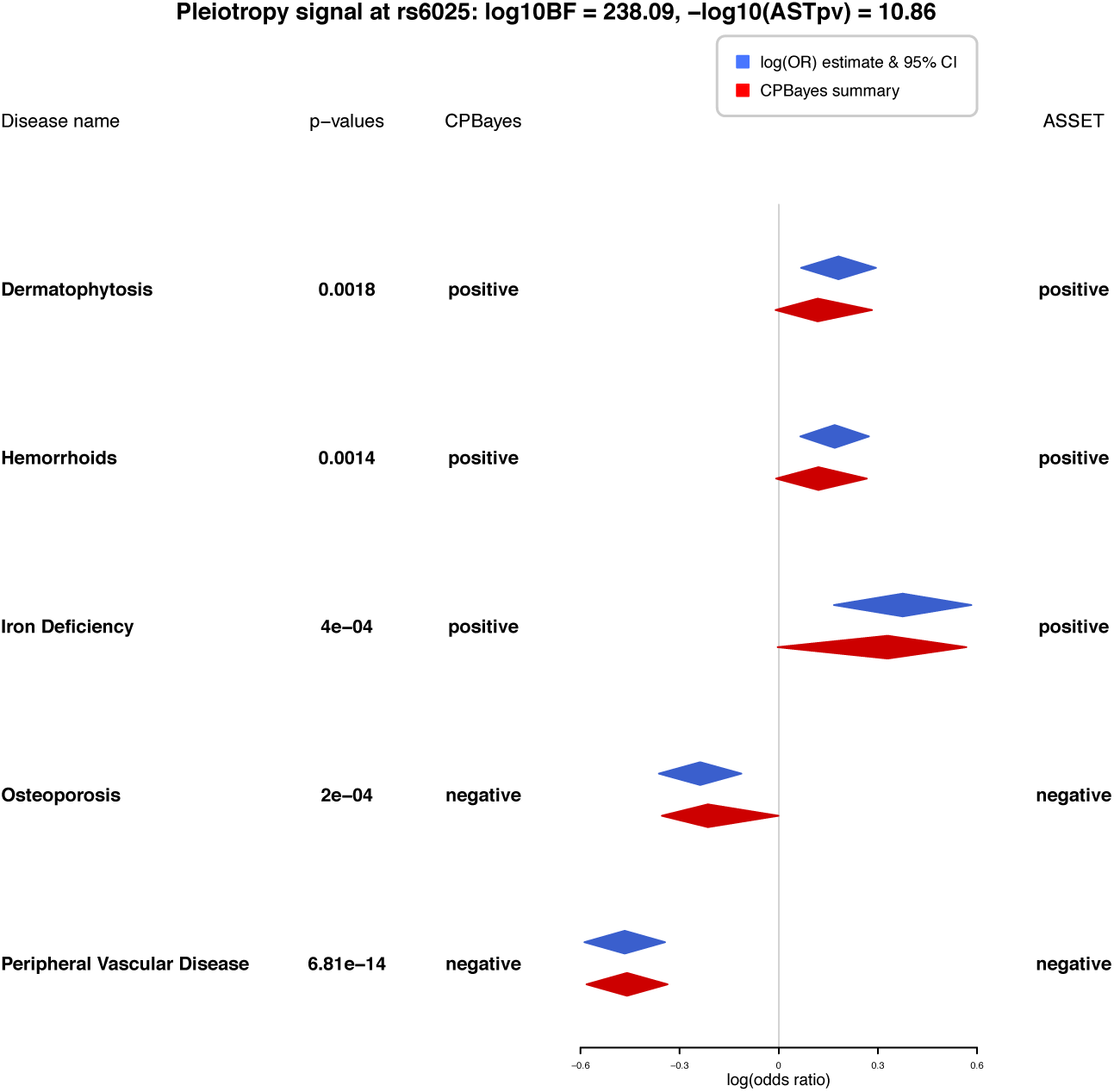
Forest plot for pleiotropic signal at rs6025 on chromosome 1 contrasting the selection of traits by CPBayes and ASSET. Phenotypes selected by either of the two methods are plotted. Blue colored bands present the trait-specific univariate log(odds ratio) estimate with the corresponding 95% confidence interval. Red colored bands present the posterior mean and 95% credible interval of the trait-specific log(odds ratio) obtained by CPBayes. The *y*-axis represents the value of log(odds ratio) as zero (null association). The log_10_BF produced by CPBayes and -log_10_(ASSET p-value) (-log_10_AST_pv_) are provided. The association status of a phenotype detected by a method is denoted by null (not associated), positive or negative (associated). The trait-specific univariate association p-values are also provided.

**Figure 5:**
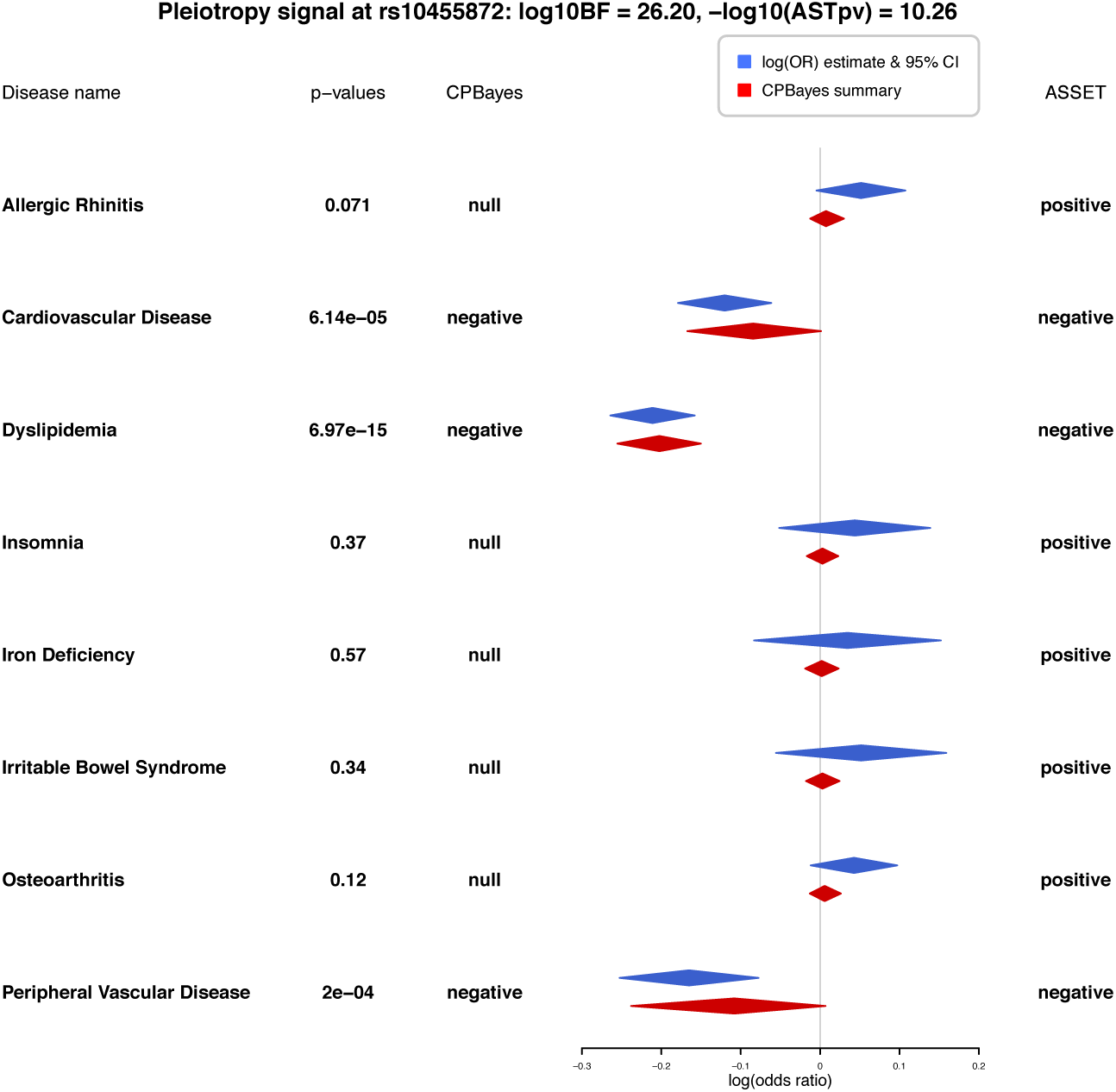
Forest plot for pleiotropic signal at rs10455872 on chromosome 6 contrasting the selection of traits by CPBayes and ASSET. Phenotypes selected by either of the two methods are plotted. Blue colored bands present the trait-specific univariate log(odds ratio) estimate with the corresponding 95% confidence interval. Red colored bands present the posterior mean and 95% credible interval of the trait-specific log(odds ratio) obtained by CPBayes. The *y-*axis represents the value of log(odds ratio) as zero (null association). The log_10_BF produced by CPBayes and -log_10_(ASSET p-value) (-log_10_ASTpv) are provided. The association status of a phenotype detected by a method is denoted by null (not associated), positive or negative (associated). The trait-specific univariate association p-values are also provided. The trait-specific univariate association p-values are also provided.

**Figure 6:**
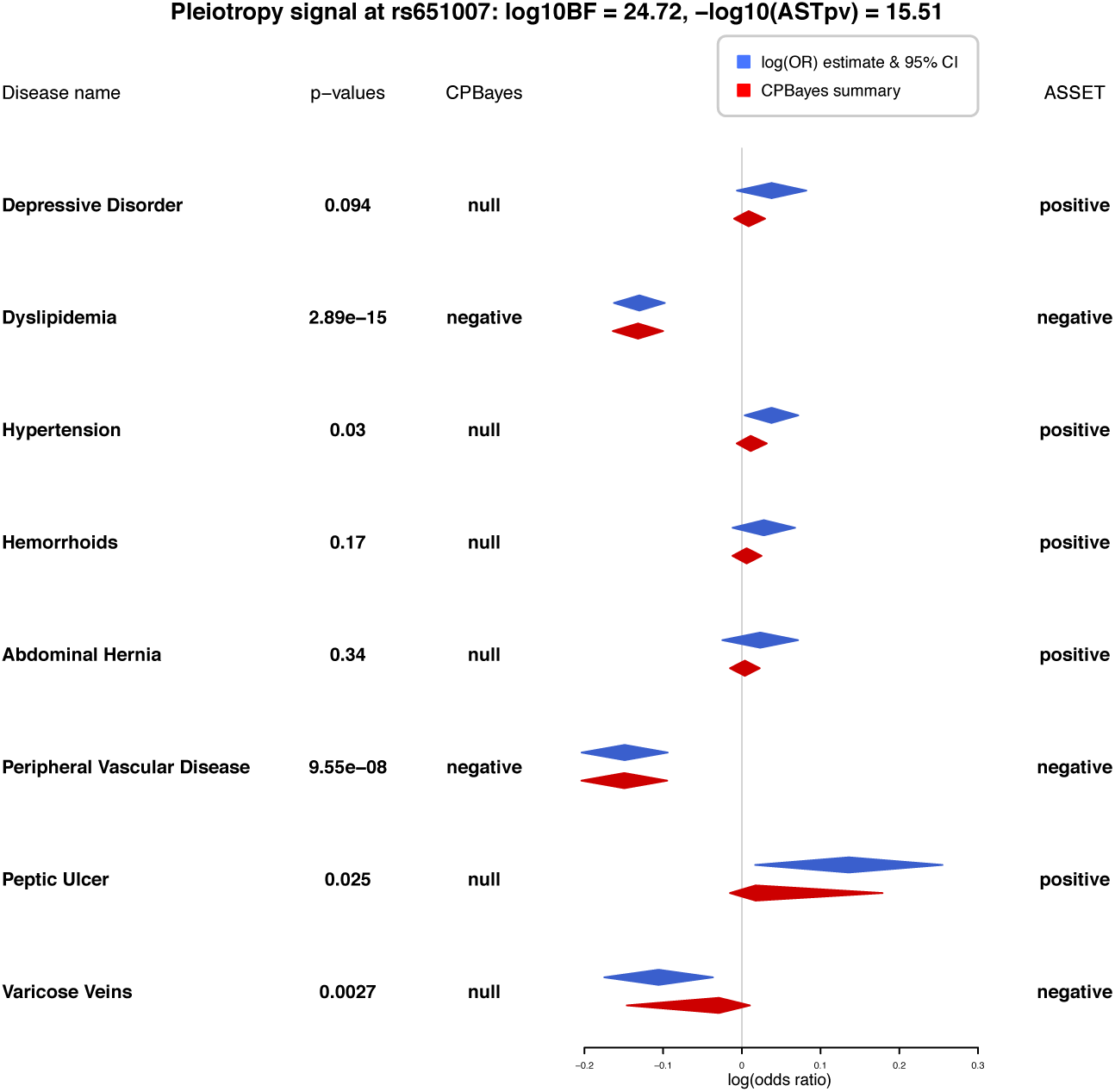
Forest plot for pleiotropic signal at rs651007 on chromosome 9 contrasting the selection of traits by CPBayes and ASSET. Phenotypes selected by either of the two methods are plotted. Blue colored bands present the trait-specific univariate log(odds ratio) estimate with the corresponding 95% confidence interval. Red colored bands present the posterior mean and 95% credible interval of the trait-specific log(odds ratio) obtained by CPBayes. The *y*-axis represents the value of log(odds ratio) as zero (null association). The log_10_BF produced by CPBayes and -log_10_(ASSET p-value) (-log_10_ASTpv) are provided. The association status of a phenotype detected by a method is denoted by null (not associated), positive or negative (associated). The trait-specific univariate association p-values are also provided.

**Figure 7:**
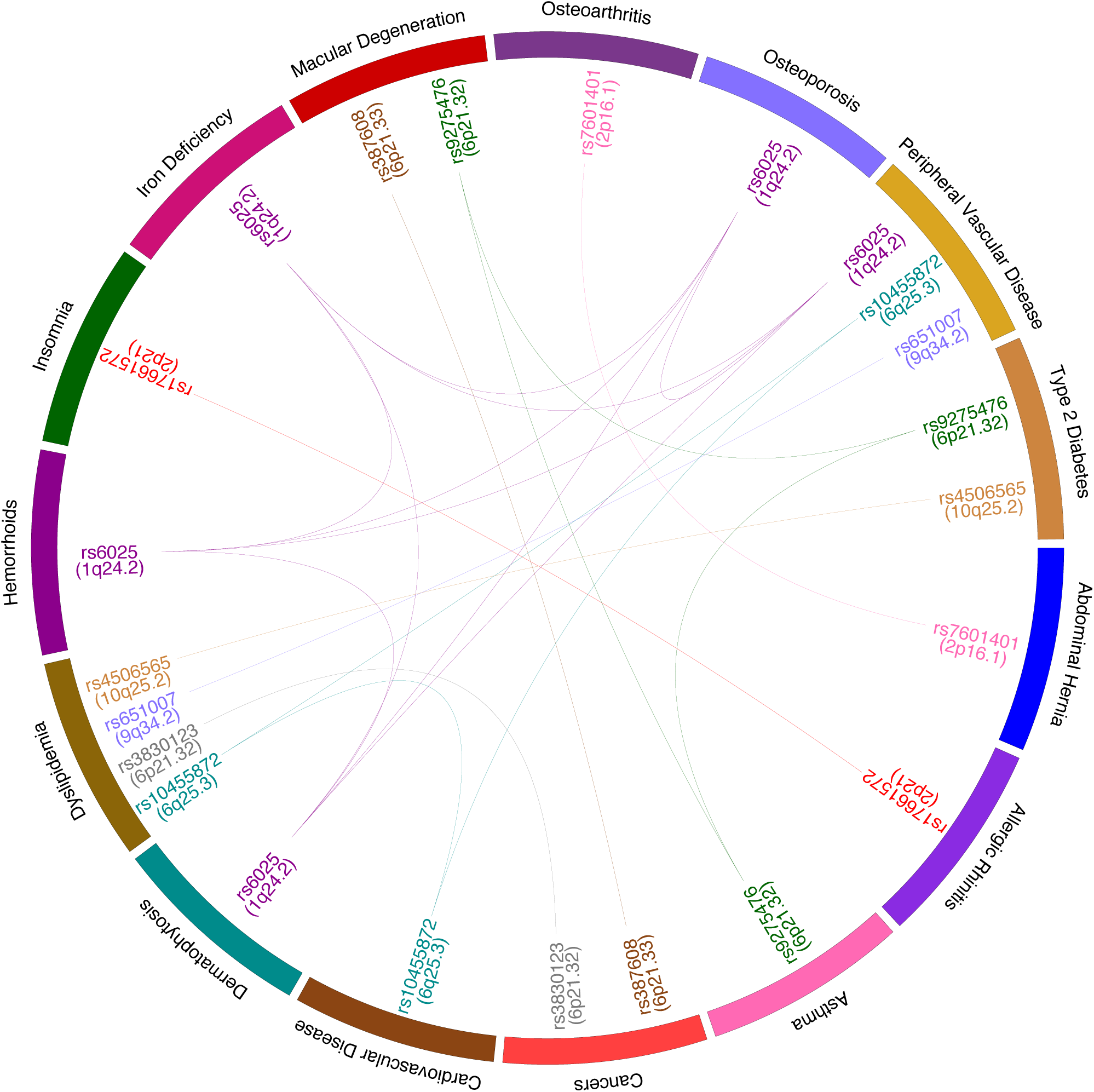
A circular diagram presenting the pairwise trait-trait pleiotropic signals detected by CPBayes.

For CPBayes, the trait-specific posterior probability of association (PPA_*j*_) provides a better insight into the relative strength of association between a pleiotropic variant and the selected non-null traits. For example, at rs6025, PPA_*j*_ for Dermatophytosis, Hemorrhoids, Iron Deficiency, Osteoporosis, and Peripheral Vascular Disease are 63.8%, 68.7%, 90.2%, 91.9%, and 100%, respectively (Table 3). This implies that the association with Peripheral Vascular Disease is the strongest among the five selected phenotypes. Even though the order of the strength of association in terms of PPA_*j*_ appeared to be the same as that given by the univariate association p-values, PPA_*j*_ provides a better interpretation. At some of the GW significant SNPs detected by CPBayes (Table 3 and S21), a few phenotypes produced a non-negligible value of PPA_*j*_ but were left out from the optimal subset of non-null traits. In Table 4, we list these SNPs and the corresponding phenotypes having a PPA_*j*_ larger than 25%. For example, at rs849135 (7p15.1), CPBayes only selected Type 2 Diabetes (Table S21), but Asthma also produced a PPA_*j*_ of 40.2%. Thus, even though the effect of rs849135 on Asthma was not strong enough to make into the optimal subset, a further consideration of the pleiotropic effect of rs849135 on Type 2 Diabetes and Asthma looks promising. At rs76075198 (19q13.31), CPBayes only selected Dyslipidemia (Table S21), but Peripheral Vascular Disease also produced a PPA_*j*_ of 44.9%. Similarly at rs115946033, CPBayes only selected Type 2 Diabetes (Table S21), but it also estimated PPA_*j*_ for Depressive Disorder as 37.2% (Table 4). At all the SNPs in Table S21, BH_0.01_ selected the same single trait selected by CPBayes. But for three SNPs: rs115946033, rs849135, and rs55975204, BH_0.01_ selected one additional trait (Table S21). Of note, at rs115946033 and rs849135, CPBayes also produced a non-negligible value of PPA_*j*_ for the additional trait selected by BH_0.01_ (Table 4) as discussed above. Thus, CPBayes and BH_0.01_ broadly agree with each other while selecting the non-null traits.

**Table 4:** Pleiotropy results by CPBayes for those SNPs in Table 3 and S21, at which some phenotypes (colored blue) were not selected in the optimal subset of non-null traits but produced a non-negligible value of trait-specific posterior probability of association (PPA_*j*_).

| rsID | chrom<br>band | CPBayes<br>log10BF | CPBayes<br>PPNA | Important<br>phenotypes | $PPA_j$ | Univariate<br>p-values |
| --- | --- | --- | --- | --- | --- | --- |
| rs115946033 | 3q25.32 | 8.59 | 6.07E-16 | Depressive Disorder | 37.2% | 0.0008 |
|  |  |  |  | Type 2 Diabetes | 100% | 3.83E-07 |
| rs849135 | 7p15.1 | 7.66 | 5.22E-15 | Asthma | 40.2% | 2.19E-05 |
|  |  |  |  | Type 2 Diabetes | 100% | 1.79E-14 |
| rs76075198 | 19q13.31 | 41.56 | 6.53E-49 | Dyslipidemia | 100% | 5.29E-11 |
|  |  |  |  | Peripheral Vascular Disease | 44.9% | 0.0067 |

At rs3791679 (Table S22), BH_0.01_ selected Abdominal Hernia (univariate p-value: 2.66 × 10^−14^) and Osteoarthritis (univariate p-value: 9.85 × 10^−5^), but ASSET missed Osteoarthritis and selected Abdominal Hernia and other traits with larger p-values (Table S22). At rs7601401 which is the lead SNP for CPBayes in the corresponding LD block containing rs3791679, both BH_0.01_ and CPBayes selected Abdominal Hernia (p-value: 3.88 × 10^−12^) and Osteoarthritis (p-value: 3.46 × 10^−6^). Similarly at rs4506565, both CPBayes and BH_0.01_ selected Type 2 Diabetes (p-value: 2.02 × 10^−55^) and Dyslipidemia (p-value: 1.33 × 10^−6^), but ASSET missed Dyslipidemia and selected Type 2 Diabetes and other traits with larger p-values (Table S24).

This indicates lower sensitivity of ASSET observed in some scenarios of our simulation study.

We note that a majority of the SNPs detected by the two methods (Table 3, S21, S22, and S23) are already reported in the NHGRI-EBI GWAS catalog. For example, rs6025 (1q24.2) has been associated with inflammatory bowel disease and venous thromboembolism. rs10455872 (6q25.3) has been associated with myocardial infarction, response to statins (LDL cholesterol change), coronary artery disease, and aortic valve calcification. rs651007 (9q34.2) has been associated with iron status biomarkers (ferritin levels), blood metabolite levels, serum alkaline phosphatase levels, and end-stage coagulation. We also note that the combined strategy of CPBayes used the uncorrelated version only for 19 SNPs among all the 601,175 SNPs analyzed, and for none of the 314 SNPs identified in the primary genome-wide screening.

## Discussion

We have proposed a Bayesian meta-analysis approach CPBayes for pleiotropic association analysis based on summary-level data. It simultaneously evaluates the evidence of aggregate-level pleiotropic association and estimates an optimal subset of traits associated with the risk locus under a unified Bayesian statistical framework. The method is implemented by Gibbs sampling designed for both uncorrelated and correlated summary statistics. We have conducted extensive simulation study and analyzed the large GERA cohort for evaluating the performance of CPBayes.

An appealing feature of CPBayes is that, in addition to log_10_BF, PPNA, and an optimal subset of non-null traits, it simultaneously provides other interesting insights into an observed pleiotropic signal. For example, it estimates a trait-specific posterior probability of association (PPA_*j*_), the direction of association, posterior mean/median, and the credible interval of the unknown true genetic effect across traits. PPA_*j*_ quantifies the marginal probability of each trait being associated with a pleiotropic variant. As demonstrated in the real data application, even if CPBayes does not select a phenotype in the optimal subset of non-null traits which is defined as the MAP estimate (see the methods section), PPA_*j*_ for the phenotype may not be negligible. It may help an investigator to better explain a pleiotropic signal. One can also define the optimal subset of associated traits as {*Y_j_*: PPA*_j_* > *p*}, where *p* can be chosen as 0.5 (known as the median model), or other values. One can evaluate the joint posterior probability of a particular subset of traits being associated. Such flexibility in making inference on pleiotropy are mainly due to the MCMC construction underlying CPBayes.

In contrast to ASSET, the major advantage of CPBayes is that it selects the non-null traits underlying a pleiotropic signal with a substantially higher accuracy. A possible reason behind this is that CPBayes performs the selection probabilistically through updating the latent association status by MCMC. ASSET selects that subset of traits as non-null which maximizes the observed value of a weighted linear combination of the normalized univariate association statistics corresponding to the phenotypes belonging to a subset. So, given the summary statistics, ASSET does not select the non-null traits probabilistically based on the distribution of the summary statistics. We also note that ASSET is based on the framework of a fixed effects meta-analysis and assumes that the effects in a given direction (positive/negative) are the same in size. But we observed in our real data application that, in a given direction, the effects of a variant across phenotypes may often be heterogeneous. CPBayes allows heterogeneity simultaneously in the direction and size of the effects. Table S1 summarizes key features of CPBayes and ASSET.

While assessing the selection accuracy, we have placed more emphasis on specificity than sensitivity. This was because a higher sensitivity at the expense of a lower specificity can lead to a false selection of too many traits as non-null. CPBayes consistently maintained a very good level of specificity while offering a good level of sensitivity across a wide range of simulation scenarios. The overall selection accuracy of BH_0.01_ was also comparable with that of CPBayes. While CPBayes produced a limited number of pleiotropic SNPs associated with more than one phenotype in the analysis of GERA cohort, these pleiotropic signals seem very promising with high specificity. Hence, the non-null traits for a pleiotropic variant selected by CPBayes are substantially more reliable than those detected by ASSET. The subsets of non-null traits selected by BH_0.01_ in the GERA cohort were consistent with CPBayes but not with ASSET which also indicates that CPBayes is a more robust alternative than ASSET. Here we note that, BH_0.01_ only facilitates the selection of associated traits underlying a pleiotropic signal but can not test for the evidence of overall pleiotropic association as CPBayes and ASSET. While CPBayes and ASSET estimate the measure of aggregate-level pleiotropic association and subset of non-null traits simultaneously under the same framework, BH_0.01_ has to be implemented in a separate step for a GW significant pleiotropic variant. Hence, CPBayes is a substantially more complete statistical tool for pleiotropy analysis than BH_0.01_.

We contrasted CPBayes with GPA for a pair of traits by simulations. We observed that the two methods perform similarly with respect to estimating the joint posterior probabilities of different configurations of association with the two traits. Both approaches use a mixture of two probability distributions – one to model null effects and the other to model non-null effects. While CPBayes directly models the effect estimates by a scale mixture of two normal distributions with mean zero (one with small variance to model the null effects), GPA models the univariate association p-values by a mixture of Uniform(0,1) [Beta(1,1)] distribution (modeling null effects) and a Beta distribution with its first shape parameter smaller than the second shape parameter (modeling non-null effects). This connection in the probabilistic modeling is a reason behind the similar performance of the two approaches.

The conditional FDR approach and GPA are mainly suited for analyzing a pair of traits at a time. For more than two traits, all possible pairs of traits have to be separately analyzed and then combined which is not a simultaneous analysis of multiple traits. [Chung et al., 2014] discussed that it is theoretically possible but challenging to extend GPA for simultaneous analysis of multiple traits. They also suggested in the R-package implementing GPA to analyze a pair of traits at one time. Since they demonstrated that GPA performs better than the conditional FDR approach with respect to accurately prioritizing risk SNPs, we compared CPBayes with GPA.

CPBayes and ASSET are explicitly designed to analyze two or more traits simultaneously. Like the Bayes factor and PPNA in CPBayes or the global p-value in ASSET, GPA does not provide a measure of aggregate-level pleiotropic association for individual SNPs. Rather it performs a statistical test to evaluate whether two traits are overall pleiotropic using GW summary statistics data. GPA fits the model by EM algorithm and CPBayes employs MCMC that allows for estimating the Bayes factor and PPNA along with the marginal or joint posterior probabilities of a specific trait or a subset of traits being associated. While CPBayes and ASSET explicitly adjust for correlation between summary statistics, GPA does not allow for such correlation. For model fitting, GPA requires summary-level data for a sufficiently large number of genome-wide variants which should include a substantial number of risk SNPs for better fitting. However, CPBayes and ASSET can be implemented for any collection of SNPs individually. Hence we conducted our main comparison study between CPBayes and ASSET which address similar objectives in a pleiotropic association study.

Note that the continuous spike inherits the infinitesimal-model assumption that every SNP contributes to the variation of a trait, and the distinction is made between a negligible and a significant contribution, whereas the Dirac spike assigns the null effects explicitly to zero. From the perspective of heritability estimation, the infinitesimal-model assumption is more relevant since many SNPs with small effects underlie the variation of a phenotype. We conducted a simulation study (provided in the supplementary material) to compare the continuous spike and the Dirac spike. We found that the continuous spike offers better accuracy in the selection of non-null traits than the Dirac spike. The continuous spike is also computationally much faster (2-3 times) than the Dirac spike. Hence, we adopted the continuous spike for constructing CPBayes. We note that, the latent association status (*Z*) could only be used in the model for the continuous spike. For the Dirac spike, the inclusion of *Z* in the model makes the corresponding MCMC reducible, and hence non-convergent to its stationary distribution (details not provided). Also, for the continuous spike, the full conditional posterior distributions of *z*_1_….,*z_K_* are independent which leads to an explicit estimation of the Bayes factor based on the MCMC sample. But, for the Dirac spike, the explicit estimation of the Bayes factor appears to be extremely difficult in the correlated case, because the full conditional posterior distributions of *β*_1_,…, *β_K_* are not independent for correlated summary statistics.

In a related work, Han and Eskin [2012] proposed a modified random effects meta-analysis for combining heterogeneous studies coupled with a Bayesian approach to provide a better interpretation of an observed signal of aggregate-level association. They investigated how to combine heterogeneous genetic studies across different populations/ethnicities. However, they did not address how to account for a possible correlation between the summary statistics while selecting the most important studies underlying an observed signal of aggregate-level association. Moreover, they assumed that the non-null effects are similar across studies which is less likely to hold in the context of pleiotropy. Hence we compared CPBayes with ASSET, GPA, and BH_0.01_.

We note that we did not explicitly compare CPBayes and ASSET with respect to power, because the Bayes factor and the p-value are not directly comparable. Computing a p-value based on the Bayes factor is computationally very expensive, and moreover, the approximation may not be accurate at a genome-wide scale. On the other hand, determining the cut-off of the Bayes factor corresponding to a given choice of the false positive rate is also time consuming and may become computationally infeasible for a false positive rate at the genome-wide scale, e.g., 5 × 10^−8^. Hence, we preserved the fully Bayesian essence of CPBayes. In the GERA cohort analysis, CPBayes primarily detected smaller number of SNPs than ASSET (314 versus 394) by using a seemingly conservative cut-off of log_10_BF (discussed in the real data application section). But, among the primarily identified genome-wide significant SNPs, CPBayes detected substantially larger number of associated LD blocks than ASSET (49 versus 30). ASSET completely missed some loci which were detected by CPBayes. These findings indicate that CPBayes is powerful to identify pleiotropic variants. Of note, we did not conduct a replication study and all the pleiotropic association signals are reported based on the analysis only in the discovery sample.

We have evaluated CPBayes primarily for SNP by SNP analysis. However, one possible approach to implement the method for a gene-based association analysis is as follows. First, implement PrediXcan [Gamazon et al., 2015] to impute the expression level of a gene. Next, regress each phenotype individually on the imputed gene expression level and compute the estimate of the association parameter (*β*) along with the corresponding standard error. Finally, we can implement CPBayes on these summary statistics to conduct a gene-level pleiotropy analysis. We note that the method can also be applied to observational studies of non-genetic exposures.

For a larger number of phenotypes, CPBayes is computationally faster than ASSET. For example, in the analysis of 22 traits in the GERA cohort, CPBayes took an average run time of 3.5 hours for 1,000 SNPs, and ASSET took an average run time of 9 hours for 1,000 SNPs. However, as the number of traits decreases, ASSET gradually becomes faster due to the reduction in the number of all possible subsets of traits. That said, CPBayes is computationally feasible and can be implemented at a genome-wide scale. As expected, the uncorrelated version of CPBayes is at least twice as fast as the correlated version of CPBayes. In future work, we aim to investigate whether the computing speed of CPBayes can be increased by using a variational Bayes approach or by using an optimization technique (e.g., EM algorithm or its variants) instead of using MCMC, while preserving the efficiency of the method. Also, we want to explore how to relax the assumption in CPBayes that 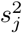 is a reasonably accurate estimate of 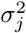 which requires a large sample size to be satisfied.

In summary, CPBayes is a sensitive and specific approach for detecting associated traits underlying a pleiotropic signal. CPBayes has a strong theoretical foundation and allows for heterogeneity in both the direction and size of effects. In addition to parameters of primary interest (e.g., the measures of overall pleiotropic association, the optimal subset of associated traits), it provides other interesting insights into a pleiotropic signal (e.g., the trait-specific posterior probability of association, the direction of association, the credible interval of unknown true genetic effect across traits). It is computationally feasible and faster than ASSET for a larger number of traits. A user-friendly R-package ‘CPBayes’ is provided for general use by other investigators.

## Appendix A

Here we state the Gibbs sampling algorithm for the continuous spike described in Equation 1. It is a desirable practice to provide a MCMC with a good initial value of the model parameters for faster convergence to its stationary distribution. Hence we use the false discovery rate controlling procedure proposed by Benjamini and Yekutieli [2001] (BY procedure) which is robust to arbitrary correlation structure of multiple test statistics. We apply the BY procedure on the univariate association p-values of *K* traits at an FDR level of 0.05 and assign *z_j_* = 1 if *Y_j_* is found to be significantly associated, otherwise set *z_j_* = 0; *j* = 1,…, *K*. We also choose an initial value of *q* as the proportion of non-null traits detected by the BY procedure (the boundary situations of no/all non-null traits are taken care of appropriately).

Define Σ_2_ = *diag*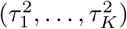 (a diagonal matrix with diagonal elements 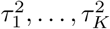), where *τ_j_* = *τ* if *z_j_* = 0; and 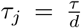 if *z_j_* = 1; *j* = 1,…, *K*. So 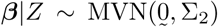. Let Σ_1_ = *S*. Also, let *β*_−*j*_ = (*β*_1_,…, *β* _*j*−1_, *β* _*j*+1_,…, *β_K_*), and *Z*_−*j*_ = (*z*_1_,…, *z*_*j*−1_,… *z*_*j*+1_,…, *z_K_*).

### Algorithm 1 Gibbs sampling for continuous spike in correlated case

1: *Start*:

2: Assign the initial values of *Z* and *q* as described above.

3: *loop*:

4: Simulate *β* from its full conditional posterior distribution: 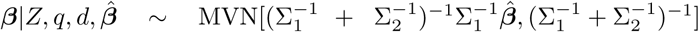.

5: For *j* = 1,…, *K*, update *z_j_* using the full conditional posterior probability: 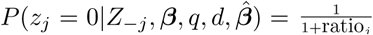, where 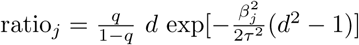.

6: Let 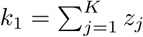, *k*_0_ = *K* − *k*_1_. Update *q* using 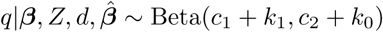.

7: We assume that *e*_1_ = *e*_2_ = 1. Update *d* from its full conditional posterior distribution in a fixed range so that the slab variance 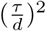 varies in a given range (*υ*_0_, *υ*_1_), and let the corresponding range of *d* be given by: *d*_0_ < *d* < *d*_1_. If 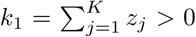, then 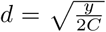, where 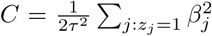, and *y* follows a truncated 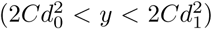 central 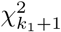 distribution. If *k*_1_ = 0, *d* is updated from the truncated (*d*_0_ < *d* < *d*_1_) Beta(1,1) distribution.

8: **goto** *loop* until all the MCMC iterations are finished.

We note that, *d* can be updated using the truncated central *χ^2^* distribution as long as the second shape parameter of its Beta prior (*e*_2_) is 1.

If the summary statistics are uncorrelated, step 4 of Algorithm 1 is modified as: for *j* = 1,…, *K*, update *β_j_* by sampling from its full conditional posterior distribution: 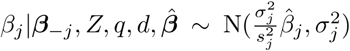, where 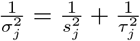 All the other steps remain the same as in the Algorithm 1.

## Author Contributions

Conceived and designed the experiments: AM SB JSW. Conducted the analysis: AM TH. Wrote the paper: AM TH JSW.

## Acknowledgements

This work was supported by the National Institutes of Health grants R01CA088164, U01CA127298, R25CA112355, and the UCSF Goldberg-Benioff Program in Cancer Translational Biology. We thank Thomas Hoffmann, Prasenjit Ghosh, and Moumita Das for important discussions relating to this work. The GERA cohort data came from a grant, the Resource for Genetic Epidemiology Research in Adult Health and Aging (RC2 AG033067; Schaefer and Risch, PIs) awarded to the GERA Permanente Research Program on Genes, Environment, and Health (RPGEH) and the UCSF Institute for Human Genetics. The RPGEH was supported by grants from the Robert Wood Johnson Foundation, the Wayne and Gladys Valley Foundation, the Ellison Medical Foundation, GERA Permanente Northern California, and the GERA Permanente National and Northern California Community Benefit Programs. The RPGEH and the Resource for Genetic Epidemiology Research in Adult Health and Aging are described in the following publication, Schaefer C. et al., The GERA Permanente Research Program on Genes, Environment and Health: Development of a Research Resource in a Multi-Ethnic Health Plan with Electronic Medical Records, In preparation. The authors do not have any conflict of interest.

## 1 Outline of mathematical derivation of the full conditional posterior distributions for continuous spike

### 1.1 Correlated case

Here we derive the full conditional posterior distributions of the model parameters to perform Gibbs sampling for correlated summary statistics. If *S* is the covariance matrix of 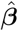,

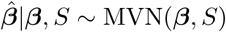

For *j* = 1,…, *K*,

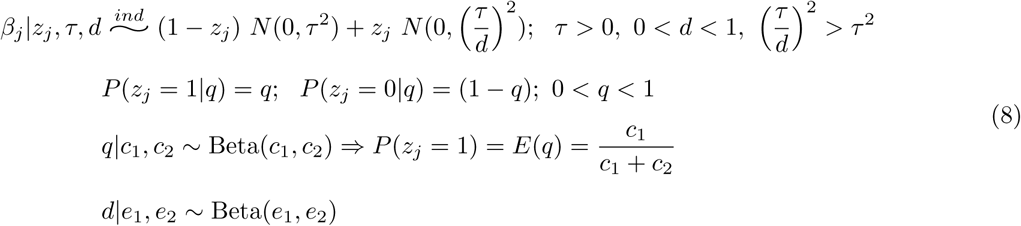

So, *β_j_*|*z_j_* = 0, *τ* ∼ N(0, *τ*^2^) and 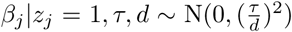. Let Σ_1_ = *S*. Define, 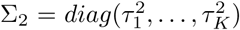 (a diagonal matrix with the diagonal elements 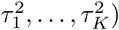, where *τ_j_* = *τ* if *z_j_* = 0, and 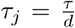 if *z_j_* = 1, *j* = 1,…, *K*. So,

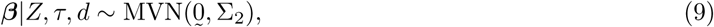

#### 1.1.1 Full conditional posterior distribution of *β*

Let [*U*] denote a generic notation of the probability distribution of a random variable *U*, and [*U*_1_|*U*_2_] denote a generic notation of the conditional probability distribution of *U*_1_ given *U*_2_. We have considered fixed choice of *τ*. Hence we drop it from the set of conditional parameters while writing the expressions of the full conditional distributions. For example, we write 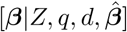 instead of 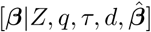. Note that,

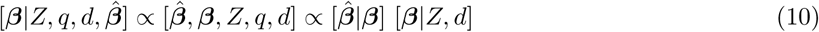

Applying standard techniques from linear algebra and distribution theory of multivariate normal, one can obtain that:

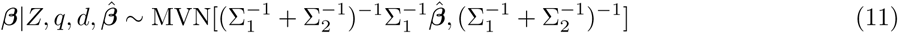

Note that Σ_2_ is dependent on *Z*, and *Z* influences the full conditional posterior distribution of *β* through the specification of Σ_2_.

#### 1.1.2 Full conditional posterior distribution of *Z*

Let *Z_−j_*, = (*z*_1_,…, *z_j−1_*, *z_j+1_*,…, *z_K_*). Note that, 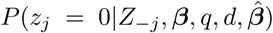 ∝ [*β_j_*|*z_j_* = 0] [*z_j_* = 0|*q*]. Similarly, 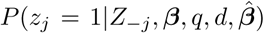 ∝ [*β_j_*|*z_j_* = 1, *d*] [*z_j_* = 1|*q*]. Combining these two equations, we obtain that:

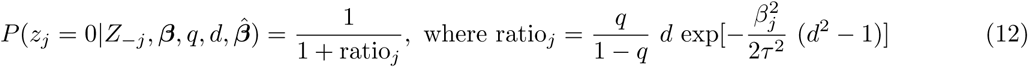

We note that the full conditional posterior distribution of *z_j_* does not depend on *Z_−j_*. Hence, the full conditional distributions of *z*_1_,…, *z_K_* are independent.

#### 1.1.3 Full conditional posterior distribution of *q*

Note that, 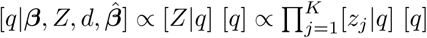. Let 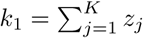, and *k*_0_ = *K* − *k*_1_. Then it can easily be derived that:

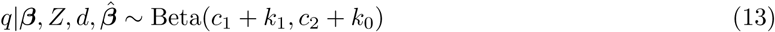

#### 1.1.4 Full conditional posterior distribution of *d*

We assume that *e*_2_ = 1 and derive a closed-form full conditional posterior distribution of *d* under this restriction. Let 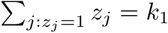.

<u>Case 1</u>: *k*_1_ > 0

Of note, 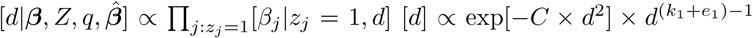, where 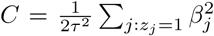. We consider the following transformation of variable: *y* = 2*Cd*^2^. It can be derived that: 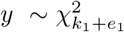 under the assumption that *y* > 0.

Suppose that, we want to update the slab variance 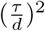 in a range (say, *υ*_0_ – *υ*_1_) such that the corresponding range of *d* is given by: *d*_0_ < *d* < *d*_1_. Using the above transformation of variable, the corresponding range of *y* is given by: 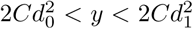. Hence, *y* ∼ truncated central 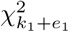, where 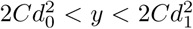. Finally, the updated *d* can be obtained by using the transformation 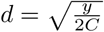.

<u>Case 2</u>: *k*_1_ =0

It can easily be shown that: 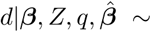 truncated Beta(*e*_1_, 1), where *d*_0_ < *d* < *d*_1_.

In the Algorithm 1 in main text, we considered *e*_1_ = 1, which is a natural choice under the absence of any prior information.

### 1.2 Uncorrelated case

When the summary statistics are uncorrelated, the full conditional posterior distributions of all the parameters except ***β*** remain the same as in the correlated case (described above). Now the full conditional distributions of *β*_1_,…, *β_κ_* become independent. For *j* = 1,…, *K*,

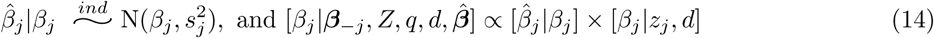

Using the above equation, it’s easy to derive the full conditional distribution as: for *j* = 1,…, *K*,

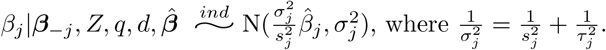

## 2 Gibbs sampling algorithm for Dirac spike

Here we outline the Gibbs sampler for the Dirac spike. We apply the BY procedure [Benjamini and Yekutieli, 2001] on the univariate association p-values of *K* traits at an FDR level of 0.05 and assign 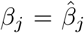 (since 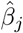 is a consistent estimator of *β_j_*) if *Y_j_* is found to be associated, otherwise we set *β_j_* = 0; *j* = 1,…, *K*. We also choose an initial value of *q* as the proportion of non-null/associated traits detected by the BY procedure (the boundary situations of no/all non-null traits are taken care of appropriately).

Let *β_−j_* = {*β*_1_,…, *β*_*j*−1_, *β*_*j*+1_,…, *β_K_*}. Consider the following partition of *S*:

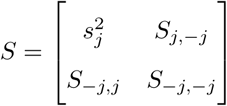

Let 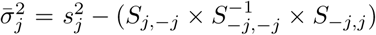, and 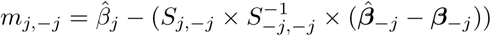.

### Algorithm S1 Gibbs sampling for the Dirac spike for correlated summary statistics

1: *Start*:

2: Assign the initial values of *β* and *q* as discussed above.

3: *loop*:

4: For *j* = 1,…, *K*, update *β*_*j*_ as follows: set *β* = 0 with probability 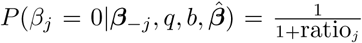, where ratio_*j*_ = 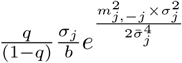. If *β_j_* is selected to be non-zero, simulate it from 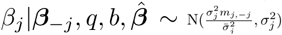, where 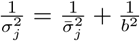.

5: Let *k*_0_ = #{*β_j_*: *β_j_* = 0, *j* = 1,…, *K*}. Update *q* using it’s full conditional posterior distribution which is a mixture of (*k*_0_ + 1) Beta distributions as follows: for *j* = 0, 1,…, *k*_0_, 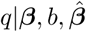 ∼ Beta(*c*_1_ + *K* − *j*, *c*_2_ + *j*) with probability 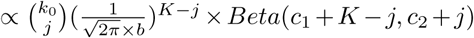. Here *Beta*(*r*_1_, *r*_2_) denotes the normalizing constant of a Beta(*r*_1_, *r*_2_) distribution.

6: Let 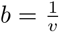, where *υ* > 0. Suppose, we want to update *b* in a given range (*b*_0_, *b*_1_), and the corresponding range of *υ* is given by: (*υ*_0_, *υ*_1_). We consider a uniform prior on *υ*. Let *k*_1_ = = *K* − *k*_0_ and *t_j_* = *K* − *j* + 1.

If *k*_1_ > 0, let 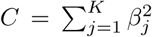. We update *υ* using the transformation: 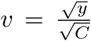, where *y* follows a mixture of (*k*_0_ + 1) distributions – the *j^th^* distribution is a truncated (between 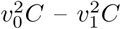) central chi-square distribution with degree of freedom *t_j_*, *j* = 0, 1,…, *k*_0_. The *j^th^* distribution is selected with probability 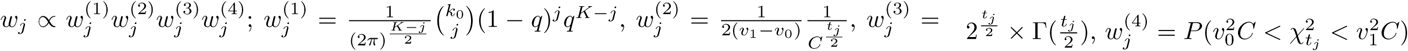, where 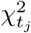 is a central chi-square distribution with d.f. *t_j_*.

If *k*_1_ = 0, the full conditional posterior distribution of *υ* is a mixture of (*K* + 1) distributions. For *j* = 0, 1,…, *K*, the *j^th^* distribution of the mixture is given by: 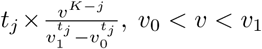; and it is selected with probability 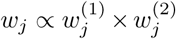, where 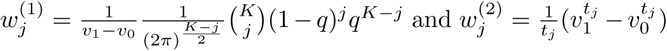.

7: **goto** *loop* until all MCMC iterations are finished.

If the summary statistics are uncorrelated, step 4 of Algorithm 2 is modified as: for *j* = 1,…, *K*, set *β*_*j*_ = 0 with probability 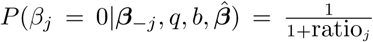, where ratio*_j_* = 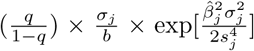, and 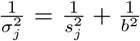; if *β_j_* is selected to be non-zero, simulate it from 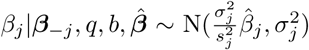. All the other steps of the algorithm remain the same.

## 3. Outline of mathematical derivation of the full conditional posterior distributions for Dirac spike

### 3.1 Correlated case

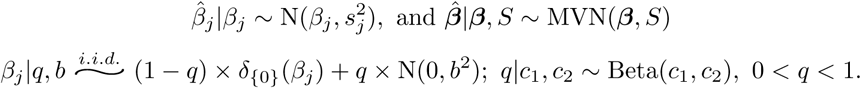

*δ*_{0}_(*β_j_*) is defined as: *δ*_{0}_(*β_j_*) = 1 when *β_j_* = 0, and *δ*_{0}_(*β_j_*) = 0 when *β_j_* ≠ 0. The full likelihood of the model is given by: 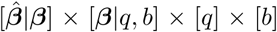 where 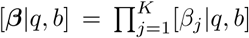. Next we derive the full conditional posterior distributions of different parameters. Let 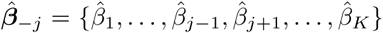 and 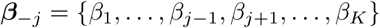.

#### 3.1.1 Full conditional posterior distribution of *β*

For *j* = 1, it can be shown that:

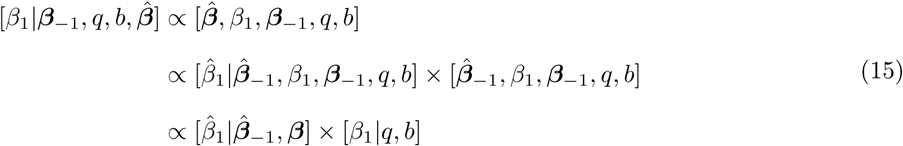

A general result: Suppose, *Y* ∼ MVN(*μ*, Σ) and (*Y*_1_, *Y*_2_) is a partition of *Y* with the corresponding partition of the mean vector and the covariance matrix as: *μ* = (*μ*_1_, *μ*_2_) and

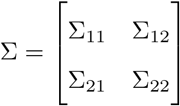

Then, 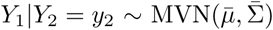 where 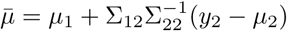 and 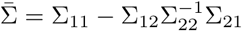.

Using the above general result, we can obtain 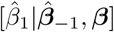 Let the partition of *S* according to the partition of *β* = (*β*_1_, *β*_−1_) be given by:

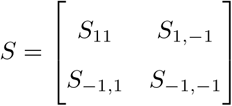

Here, 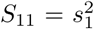 is a scalar, 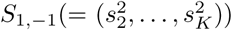 is a vector of length (*K* − 1), *S*_−1_, _− 1_ is a matrix of order (*K* − 1) × (*K* − 1). Thus 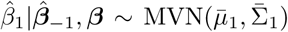, where 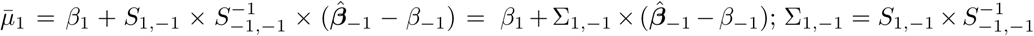. And, 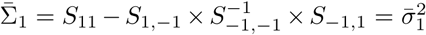, say. Thus,

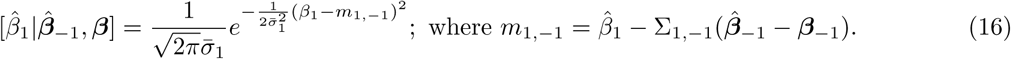

Hence,

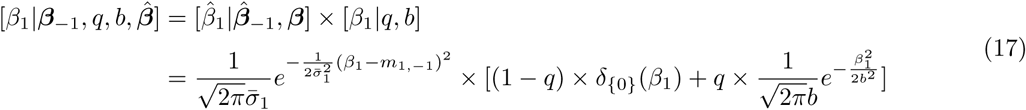

It is straightforward to derive from the above equation that:

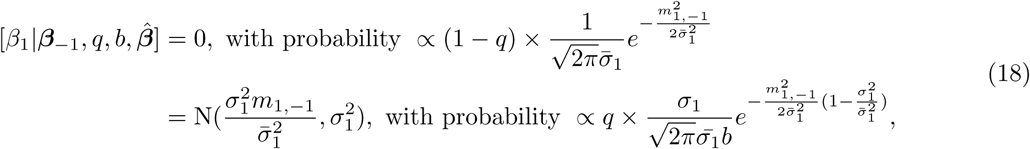
where 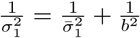. Hence, 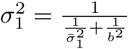.

More explicitly,

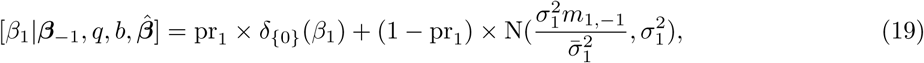
where 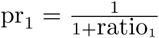, and 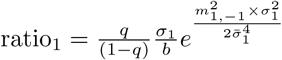

#### 3.1.2 Full conditional posterior distribution of *q*

Next we derive the full conditional distribution of *q*. Let *k*_0_ be the number of zeros in *β*, and *k*_1_ (= *K* − *k*_0_) be the number of non-zero elements in *β*. Let *dnorm*(*x*, *μ*, *σ*) denote the probability density function of a normal distribution at *x* with mean *μ* and variance *σ*^2^. Under the Dirac spike, since *β_j_* has a positive mass (1 − *q*) at 0, [*β_j_* = 0|*q*, *b*] = (1 − *q*) + *q* × *dnorm*(0, 0, *b*). Let *β_obs_* denote an observed value of *β* in a MCMC iteration.

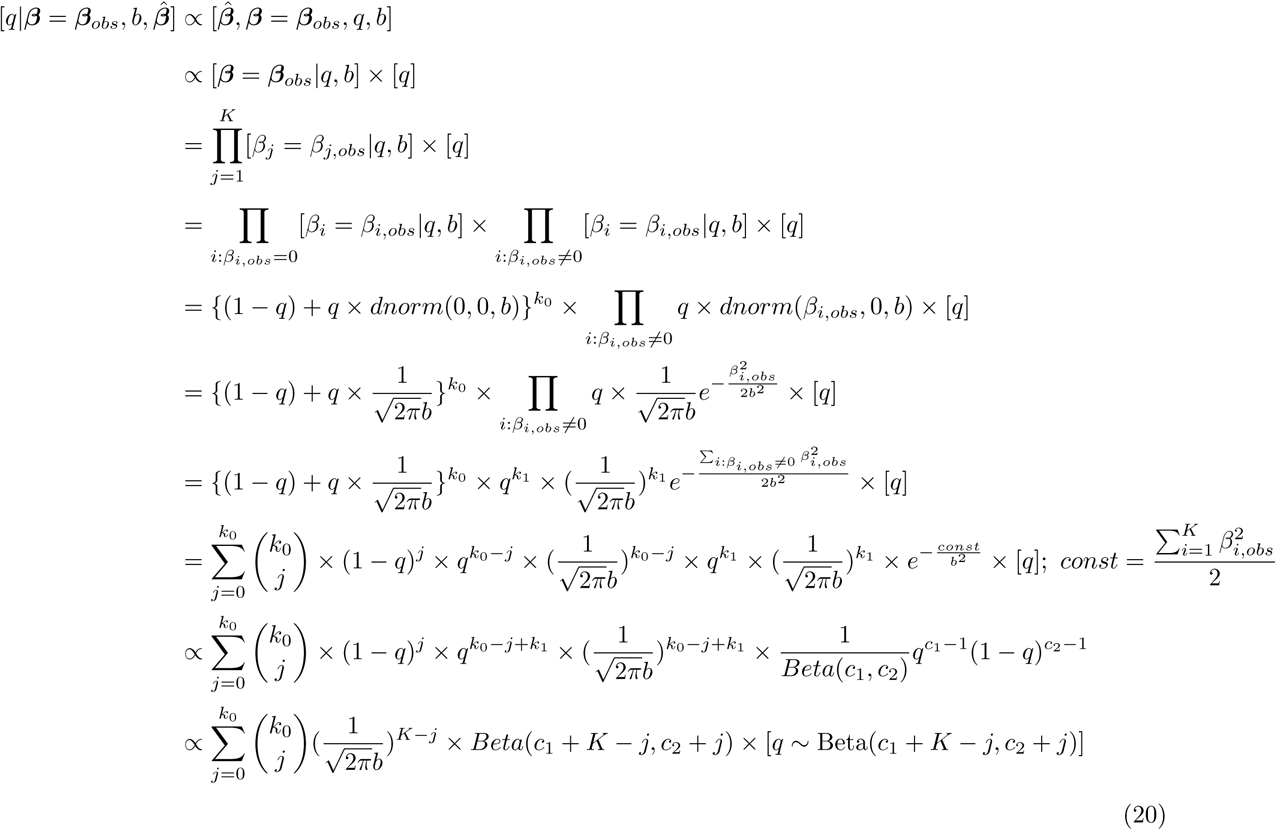

Thus, the full conditional posterior distribution of *q* is a mixture of (*k*_0_ + 1) Beta distributions as follows: for 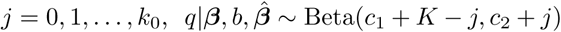 with probability 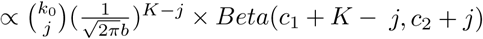. Here *Beta*(*r*_1_, *r*_2_) denotes the normalizing constant of the Beta(*r*_1_, *r*_2_) distribution.

#### 3.1.3 Full conditional posterior distribution of *b*

Since *b* > 0, let 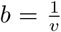 where *υ* > 0. Thus, for 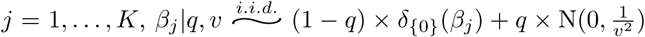. Suppose, we want to update *b* in a given range (*b*_0_, *b*_1_). Let the corresponding range of *υ* be given by (*υ*_0_, *υ*_1_). We assume a uniform prior on *υ*. So, 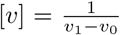, where *υ*_0_ < *υ* < *υ*_1_. Suppose, (*β*_1, *obs*_,…, *β*_*K*, *obs*_) denote an observed value of (*β*_1_,…, *β_K_*) in a MCMC iteration. Let *k*_1_ = #{*β_j_* ≠ 0: *j* = 1,…, *K*}.

<u>Case 1</u>: *k*_1_ > 0

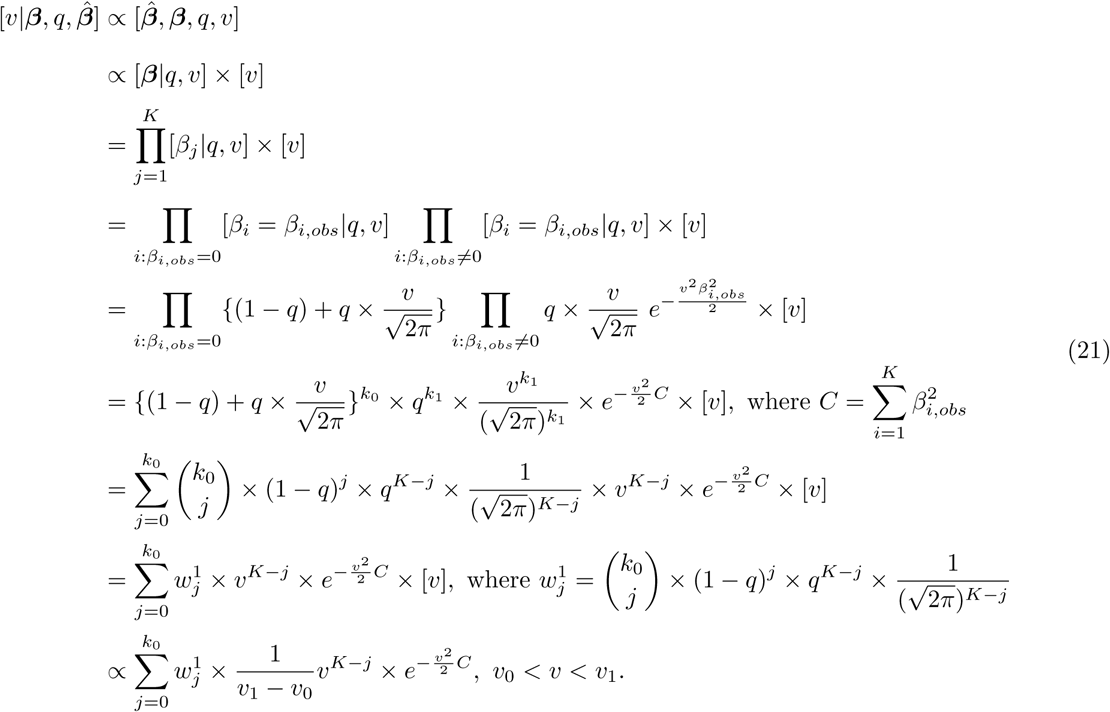

Now we consider the transformation: 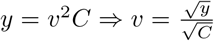, and 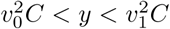. Using the above equation, we obtain that *y* follows a mixture of (*k*_0_ + 1) truncated central chi-square distributions as follows: for *j* = 0,…, *k*_0_, *y* ∼ truncated central 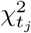 with probability *ω_j_*, where 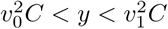 and *t_j_*, = *K* − *j* + 1. The mixture weight *ω_j_*, is given by: 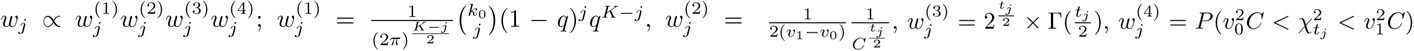. The updated *υ* is obtained from updated *y* using the transformation: 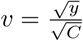.

<u>Case 2</u>: *k*_1_ =0

Similarly, if *k*_1_ = 0, we can derive the full conditional posterior distribution of *υ* which appears to be a mixture of (*K* + 1) distributions. For *j* = 0, 1,…, *K*, the *j^th^* distribution of the mixture is given by: 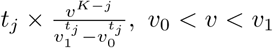. Here *t_j_*, = *K* − *j* + 1. The probability of the *j^th^* mixture component is given by: 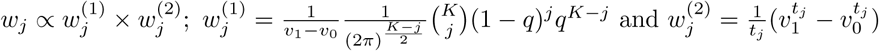.

### 3.2 Uncorrelated case

For uncorrelated summary statistics, the full conditional posterior distribution of all the parameters except *β* remain the same. The derivation of full conditional posterior distribution of *β* for uncorrelated summary statistics is straightforward and will easily follow from the derivation for correlated summary statistics.

## 4. CPBayes model diagram

Next we present two diagrams that explain the Bayesian model underlying CPBayes.

**Figure S1:**
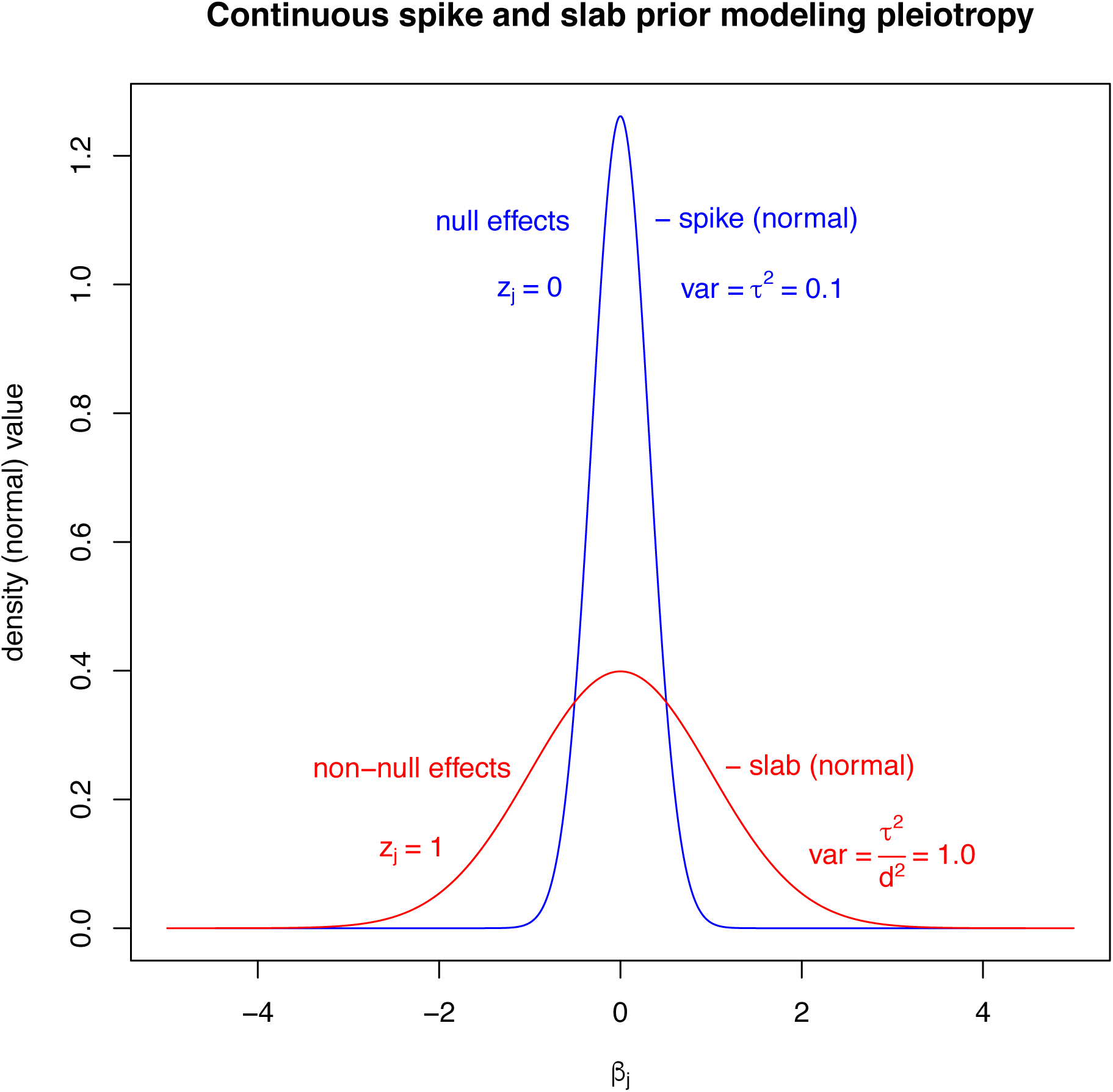
An example diagram of the continuous spike and slab prior used by CPBayes to model pleiotropy. In this diagram, the spike variance is chosen as 0.1. However, we set this value to 10^−4^ in our simulation study and real data analysis (a diagram corresponding to this choice is presented in Figure S2.)

**Figure S2:**
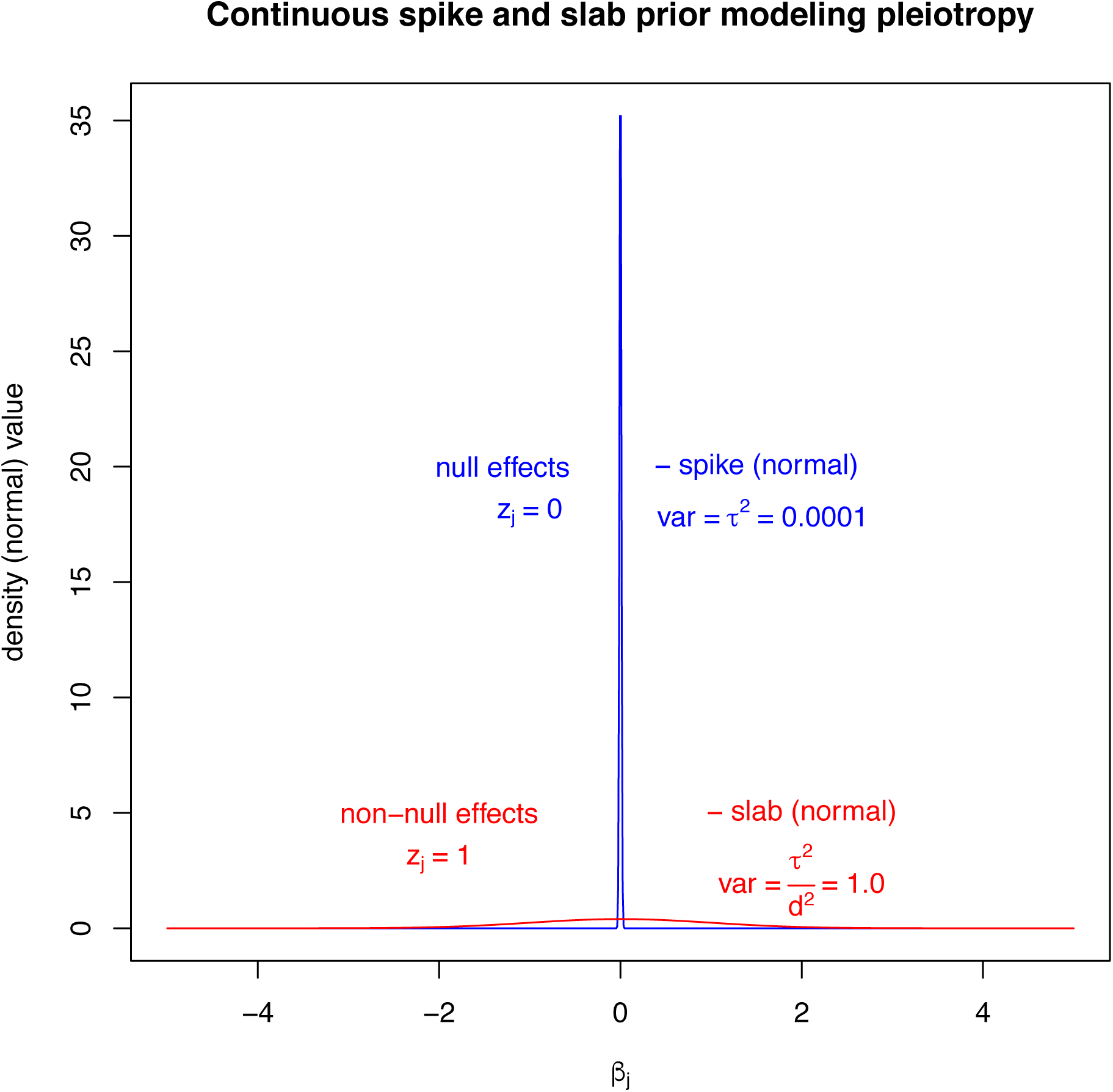
A diagram presenting the continuous spike and slab prior modeling pleiotropy with the spike variance *τ*^2^ = 10^−4^.

## 5 Comparison between main features of CPBayes and ASSET

Next we provide a table presenting a comparison between the key features of CPBayes and ASSET.

**Table S1:** Main features of CPBayes and ASSET

| Features | CPBayes | ASSET |
| --- | --- | --- |
| Paradigm | Bayesian | Frequentist |
| Measure of overall pleiotropic association | Bayes factor<br>PPNA | P-value |
| Simultaneous selection of non-null traits | Yes | Yes |
| Specificity of selection | High | Low-moderate |
| Sensitivity of selection | Moderate-high | Low-high |
| Heterogeneity in direction of effects | Yes | Yes |
| Heterogeneity in size of effects | Yes | No |

## 6 Comparison between CPBayes and GPA

We compare CPBayes with GPA for two non-overlapping case-control studies since GPA does not account for shared controls. As the summary statistics are uncorrelated here, we implement the uncorrelated version of CPBayes. We consider 1000 SNPs of which *r*% (*r* = 1, 2) are risk SNPs and (100 − *r*)% are null SNPs (not associated with both traits). SNPs are considered to be in linkage equilibrium (a modeling assumption in GPA). We simulate the minor allele frequency (MAF) at all the SNPs randomly from a Uniform(0.05, 0.5) distribution. For a risk SNP, we consider that one or both of the traits are associated. The odds ratio (OR) for a non-null trait is randomly simulated from Uniform(1.05, 1.25). We assign a larger OR to a risk SNP having a lower MAF and a smaller OR to a risk SNP having a larger MAF. In a given simulation scenario, we implement both the methods and estimate the joint posterior probability of four possible configurations of association with two traits: none of the traits (*p*_00_), 1^st^ trait but not the 2^nd^ trait (*p*_10_), 2^nd^ trait but not the 1^st^ trait (*p*_01_), both the traits (*p*_11_) are associated. Note that, this joint posterior probability is a key component of GPA to evaluate pleiotropy at a particular SNP. We present the estimates of the joint posterior probabilities obtanied by CPBayes and GPA for the risk SNPs and first 10 null SNPs in each simulation scenario in Figure S3 – S12.

In Figure S3 – S6, we present the posterior probabilities for first 10 null SNPs in four different simulation scenarios. For the null SNPs, the estimate of *p*_00_ should be high and the estimates of *p*_10_, *p*_01_, *p*_11_ should be close to zero. In Figure S3, 1% of 1000 SNPs are risk SNPs and associated only with the second trait. We observe that *p*_00_ is high for both CPBayes and GPA for the null SNPs. However, *p*_10_ for GPA was observed to be consistently larger than zero (Figure S3). But in Figure S4, this pattern for GPA is eliminated when the percentage of risk SNPs increases from 1% to 2%. Here, both CPBayes and GPA estimated *p*_00_ to be high. Chung et al. [2014] mentioned that the model fitting of GPA improves with an increase in the proportion of risk SNPs. However, the model fitting of CPBayes does not depend on the proportion of risk SNPs, because it works on each SNP individually. For SNP10 in Figure S5, CPBayes estimated a larger value of *p*_10_. We could not figure out a specific reason behind this, and suspect it to be a consequence of random fluctuation of the MCMC underlying CPBayes. In practice, MCMC can sometimes depend on the choice of the seed for random number generation and other computational issues. We also performed simulations for 1000 SNPs with all of them being null and observed that CPBayes consistently estimates *p*_00_ to be substantially higher than GPA (results not presented for brevity of space). In overall, CPBayes seems to be more robust than GPA with respect to correctly reflecting no association between a SNP and a pair of traits.

In Figure S7 − S12, we present the results for risk SNPs. In Figure S7, 1% of 1000 SNPs are risk SNPs and associated only with the second trait. So for these risk SNPs, estimated *p*_01_ should be high and *p*_00_, *p*_10_, *p*_11_ should be small. Here, CPBayes and GPA produce very similar estimates (Figure S7). For SNP1 – SNP7, most of the total posterior probability is placed on the ‘01’ association configuration of the traits. However, for SNP8 – SNP10 with larger MAF and smaller OR for the non-null trait, both the methods placed higher mass on the ‘00’ configuration which is expected from the perspective of power and sample size.

In Figure S8, 1% of 1000 SNPs are risk SNPs and associated with both the traits. So the estimate of *p*_11_ should be larger than that of *p*_00_, *p*_10_, and *p*_01_. For SNP1 – SNP6, both the methods estimated *p*_11_ very close to one (Figure S8). For SNP3, CPBayes estimated *p*_11_ to be marginally smaller than GPA. For SNP7 and SNP8, GPA placed slightly higher probability on the null association configuration ‘00’ than CPBayes (Figure S8). But, GPA estimated *p*_11_ to be higher than CPBayes while CPBayes assigned a bit of mass to the ‘10’ and ‘01’ configurations. For SNP9, both the methods estimated higher probability for the null association with CPBayes estimating *p*_00_ marginally smaller (Figure S8). For SNP10, CPBayes performed a bit conservatively and estimated *p*_10_ to be high, whereas GPA estimated *p*_11_ to be high. Here, both the traits are associated and CPBayes estimated *p*_10_ to be large instead of *p*_01_, because OR1 > OR2. In Figure S9 – S12, we observed a similar comparative performance of CPBayes and GPA as in Figure S7 and S8.

In summary, CPBayes and GPA produced similar estimates of the joint posterior probabilities across various simulation scenarios. For one SNP in Figure S8 and S12, CPBayes performed a bit conservatively missing the signal of association with the second trait having a small OR of 1.05 in both cases. We argue that this is an indirect consequence of the feature of CPBayes that it is more robust than GPA with respect to correctly identifying the global null association (both traits are unassociated). It is possible to increase the sensitivity of CPBayes by choosing a smaller value of the slab variance. However, we emphasized more on higher specificity and robustness of CPBayes in a wide range of scenarios of pleiotropy across two or more number of phenotypes. We also did simulations for one negatively associated trait and both positively and negatively associated traits. We observed similar behavior of CPBayes and GPA (results not shown to save space) as discussed above.

**Figure S3:**
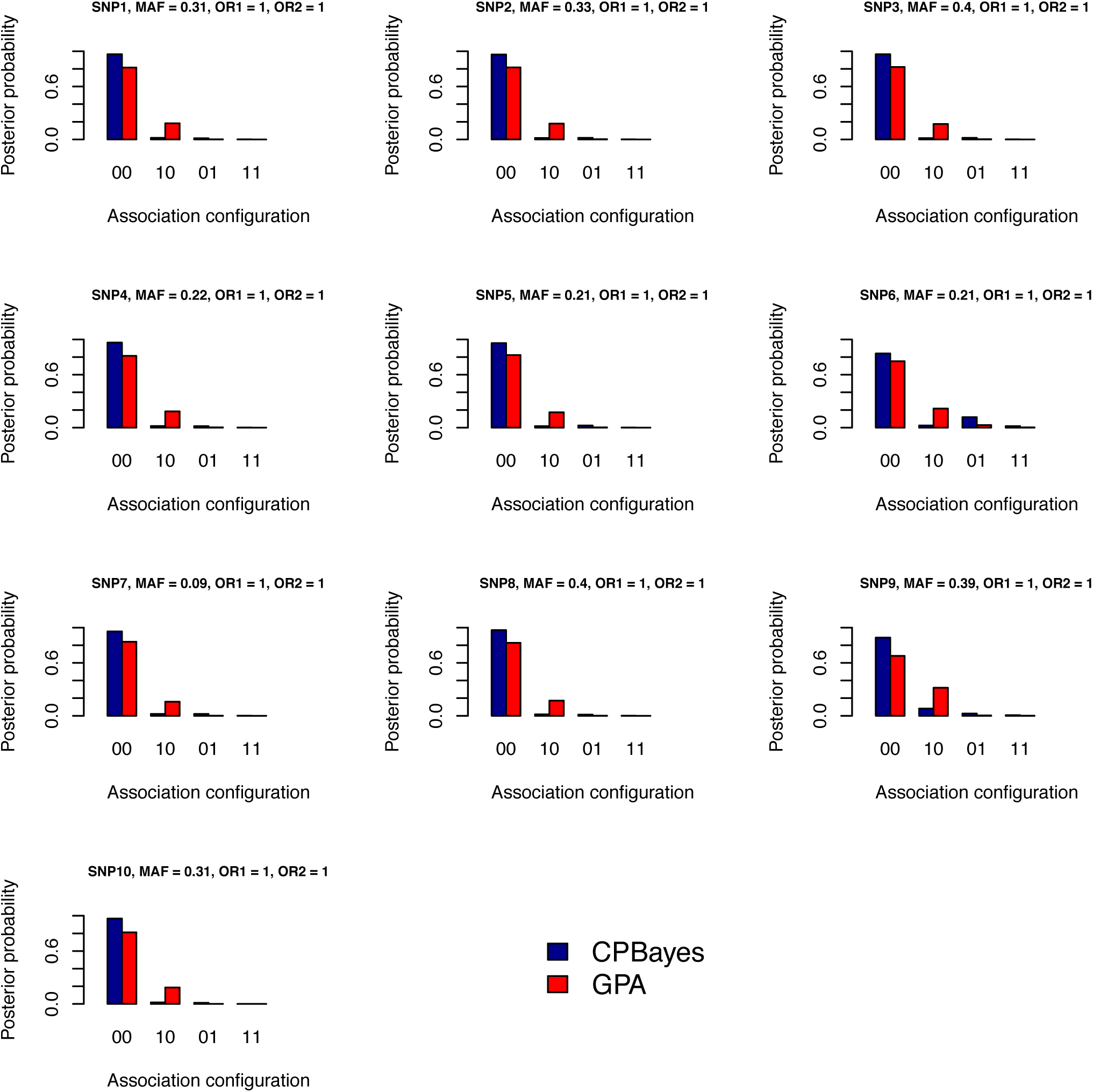
Estimated joint posterior probabilities of the trait configurations obtained by CPBayes and GPA for first 10 null SNPs. Here 1% of 1000 SNPs are risk SNPs and associated only with the second trait and 99% SNPs are null.

**Figure S4:**
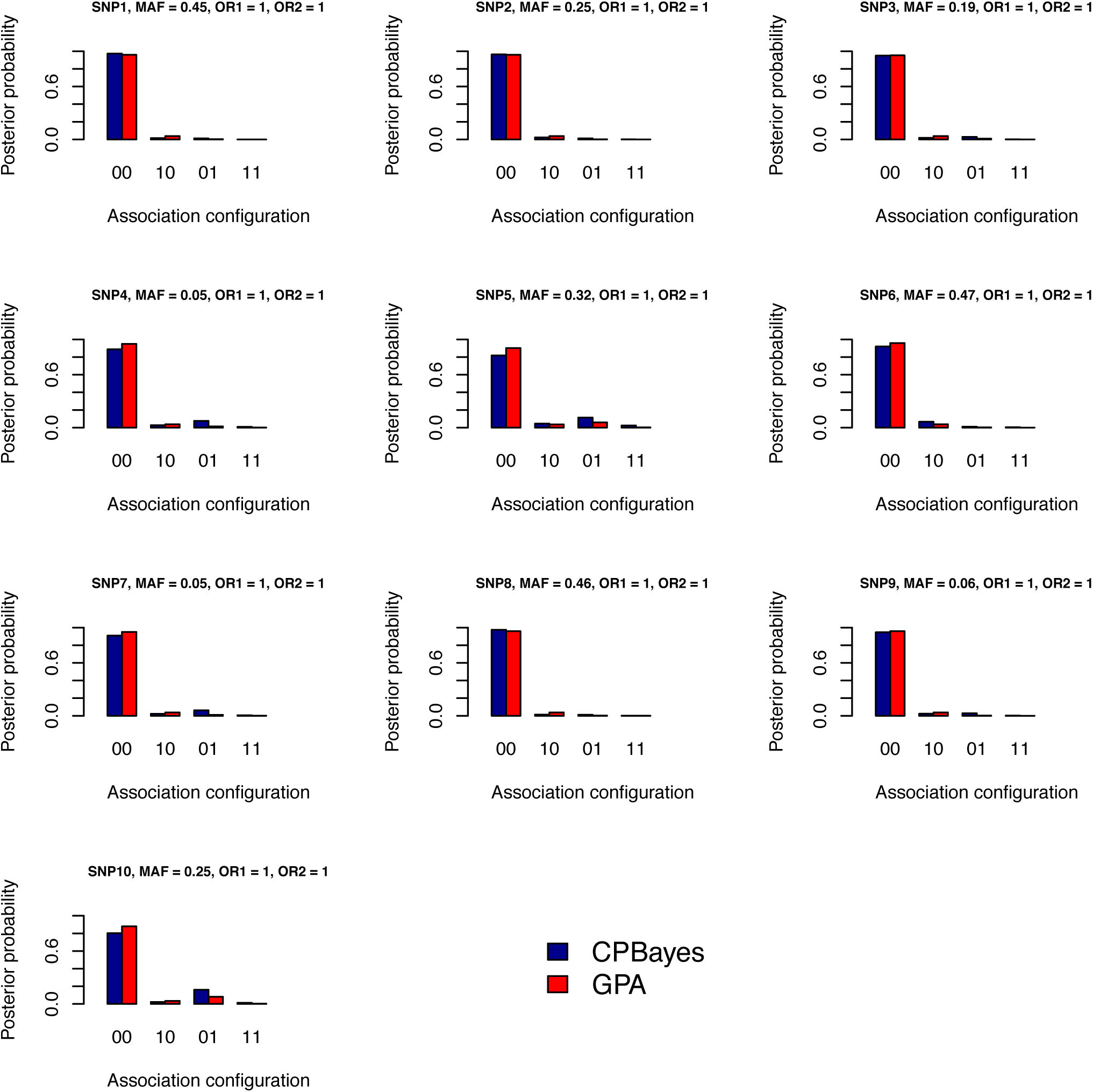
Estimated joint posterior probabilities of the trait configurations obtained by CPBayes and GPA for first 10 null SNPs. Here 2% of 1000 SNPs are risk SNPs and associated only with the second trait and 98% SNPs are null.

**Figure S5:**
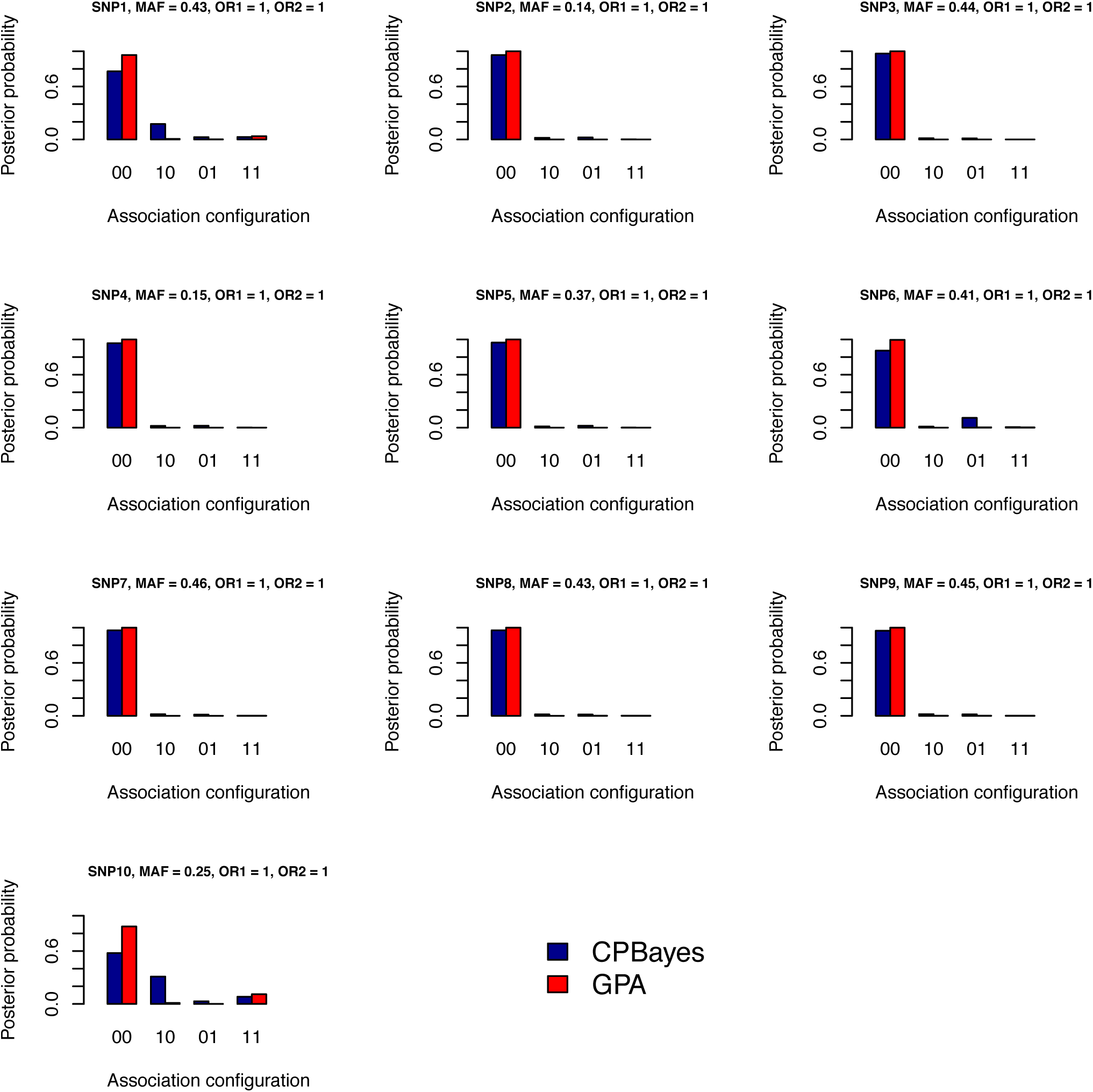
Estimated joint posterior probabilities of the trait configurations obtained by CPBayes and GPA for first 10 null SNPs. Here 1% of 1000 SNPs are risk SNPs and associated with both the traits and 99% SNPs are null.

**Figure S6:**
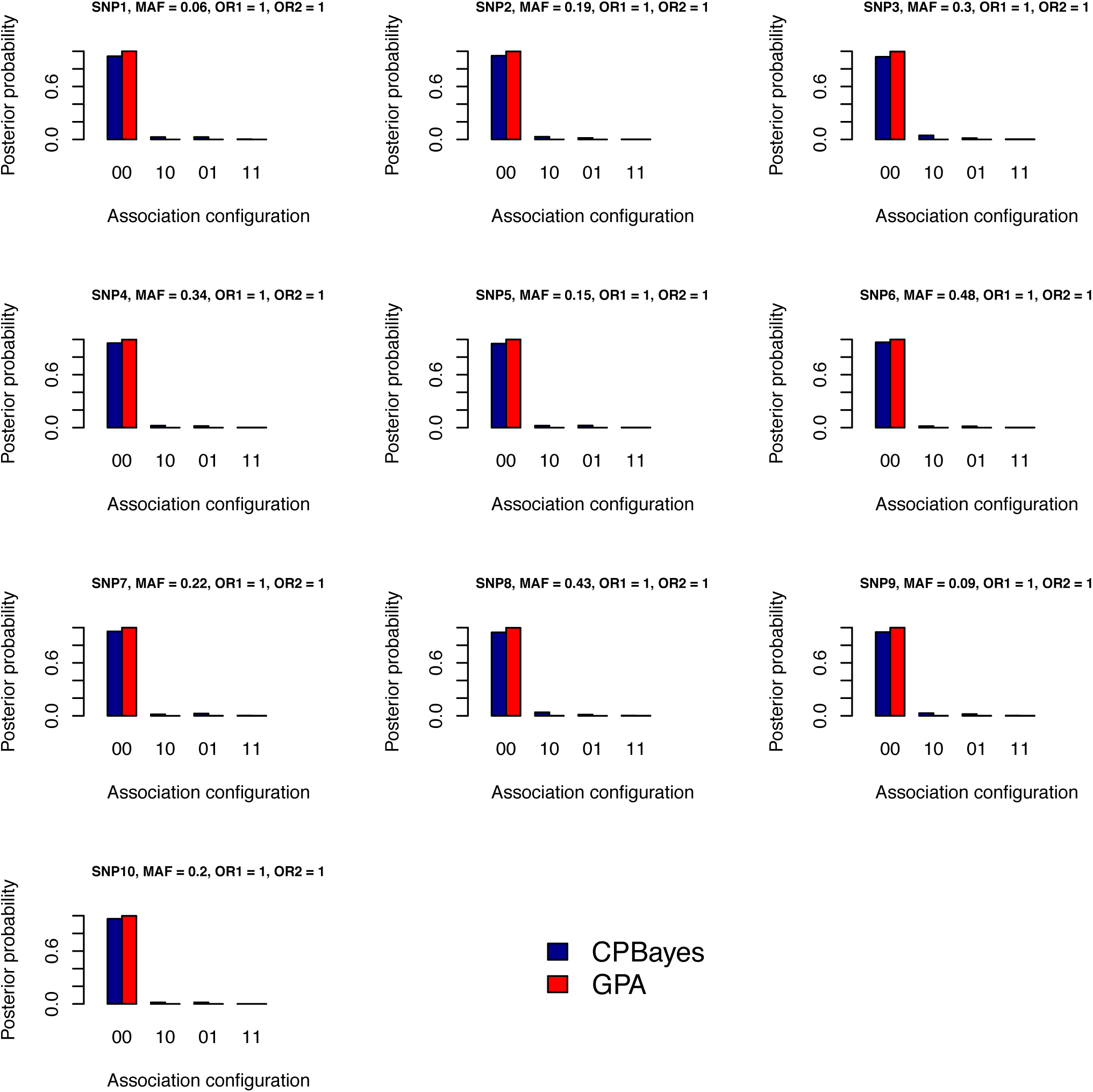
Estimated joint posterior probabilities of the trait configurations obtained by CPBayes and GPA for first 10 null SNPs. Here 2% of 1000 SNPs are risk SNPs and associated with both the traits and 98% SNPs are null.

**Figure S7:**
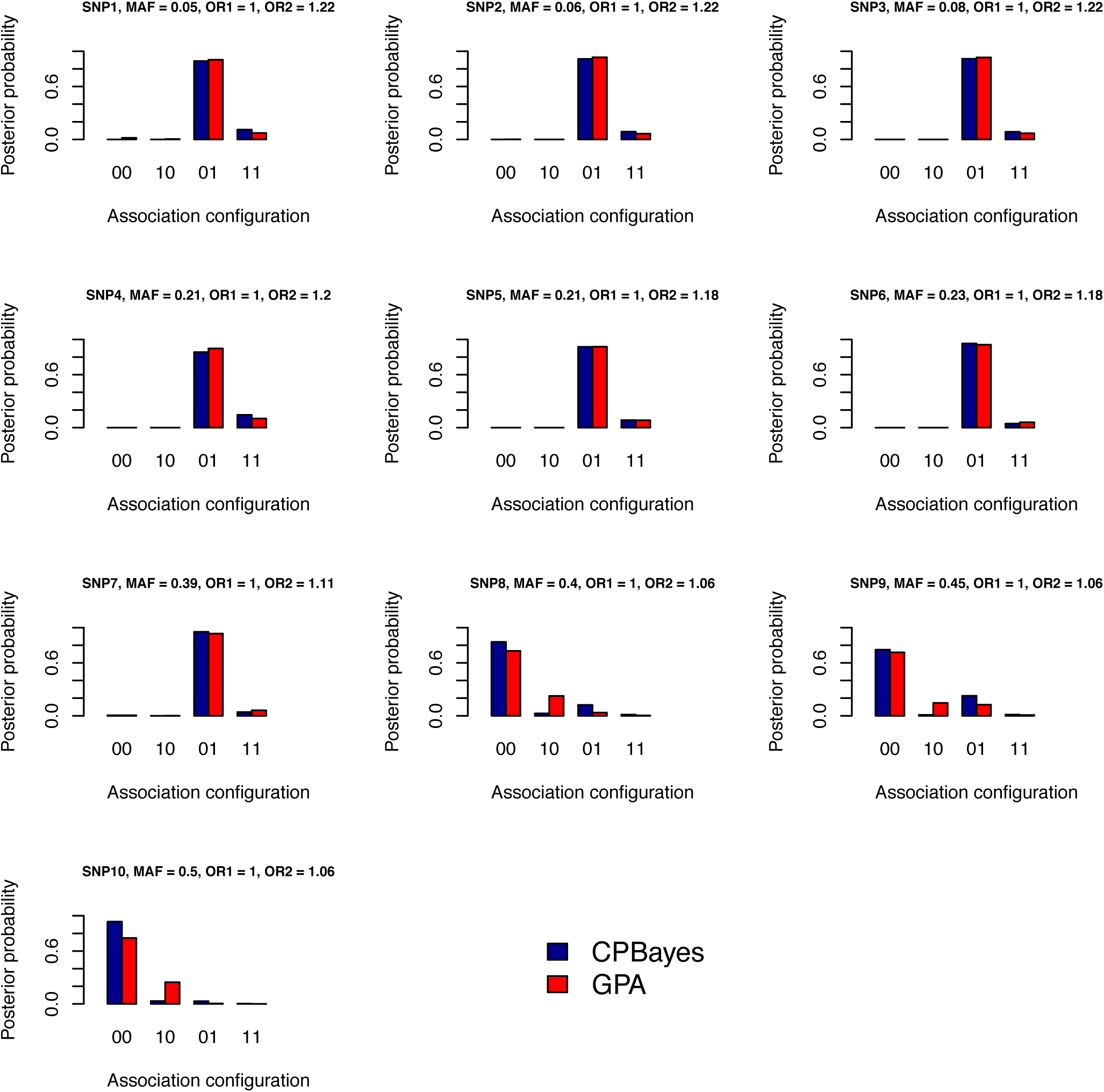
Estimated joint posterior probabilities of the trait configurations obtained by CPBayes and GPA for the risk SNPs. Here 1% of 1000 SNPs are risk SNPs and associated only with the second trait and 99% SNPs are null.

**Figure S8:**
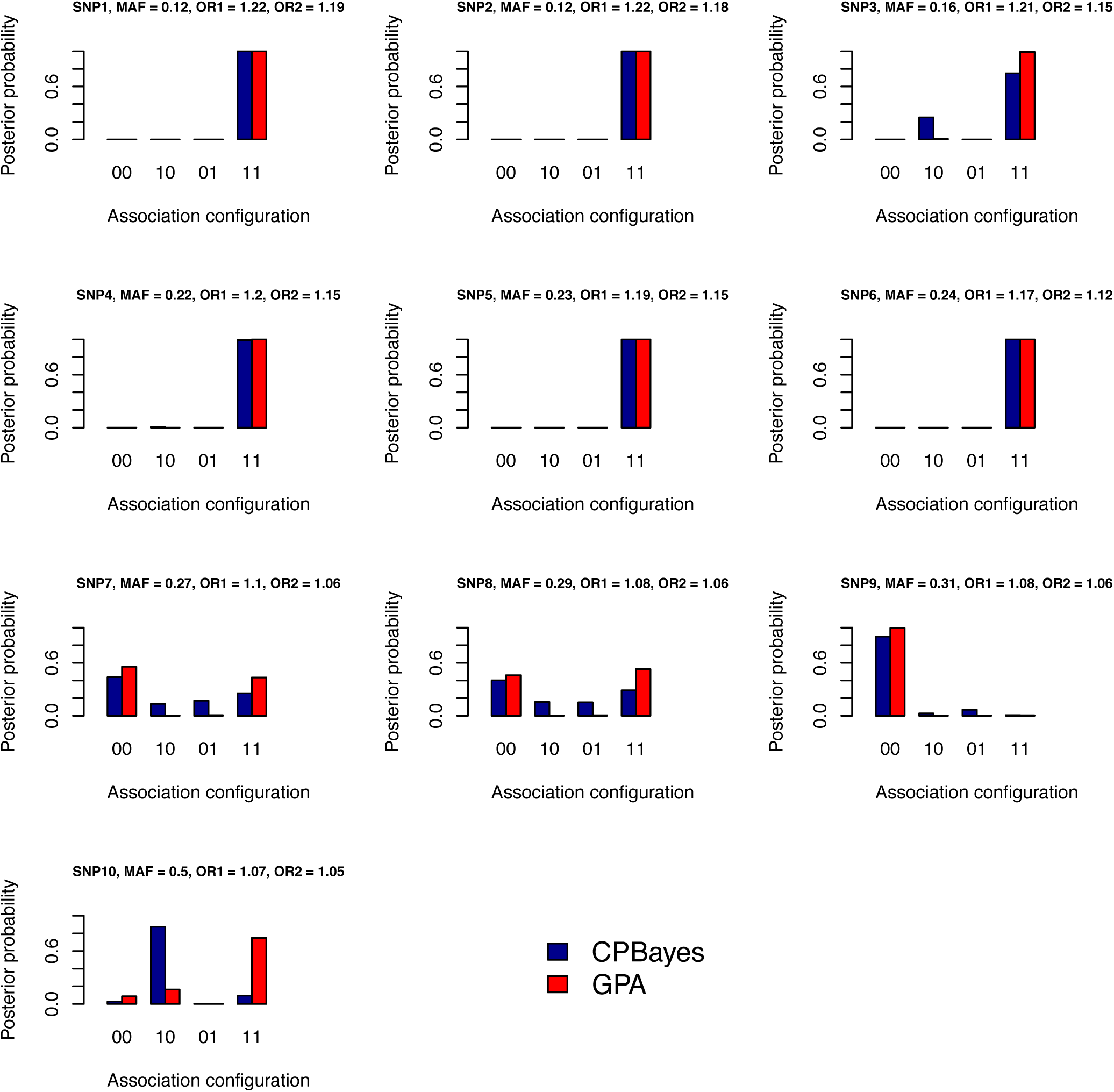
Estimated joint posterior probabilities of the trait configurations obtained by CPBayes and GPA for the risk SNPs. Here 1% of 1000 SNPs are risk SNPs and associated with both the traits and 99% SNPs are null.

**Figure S9:**
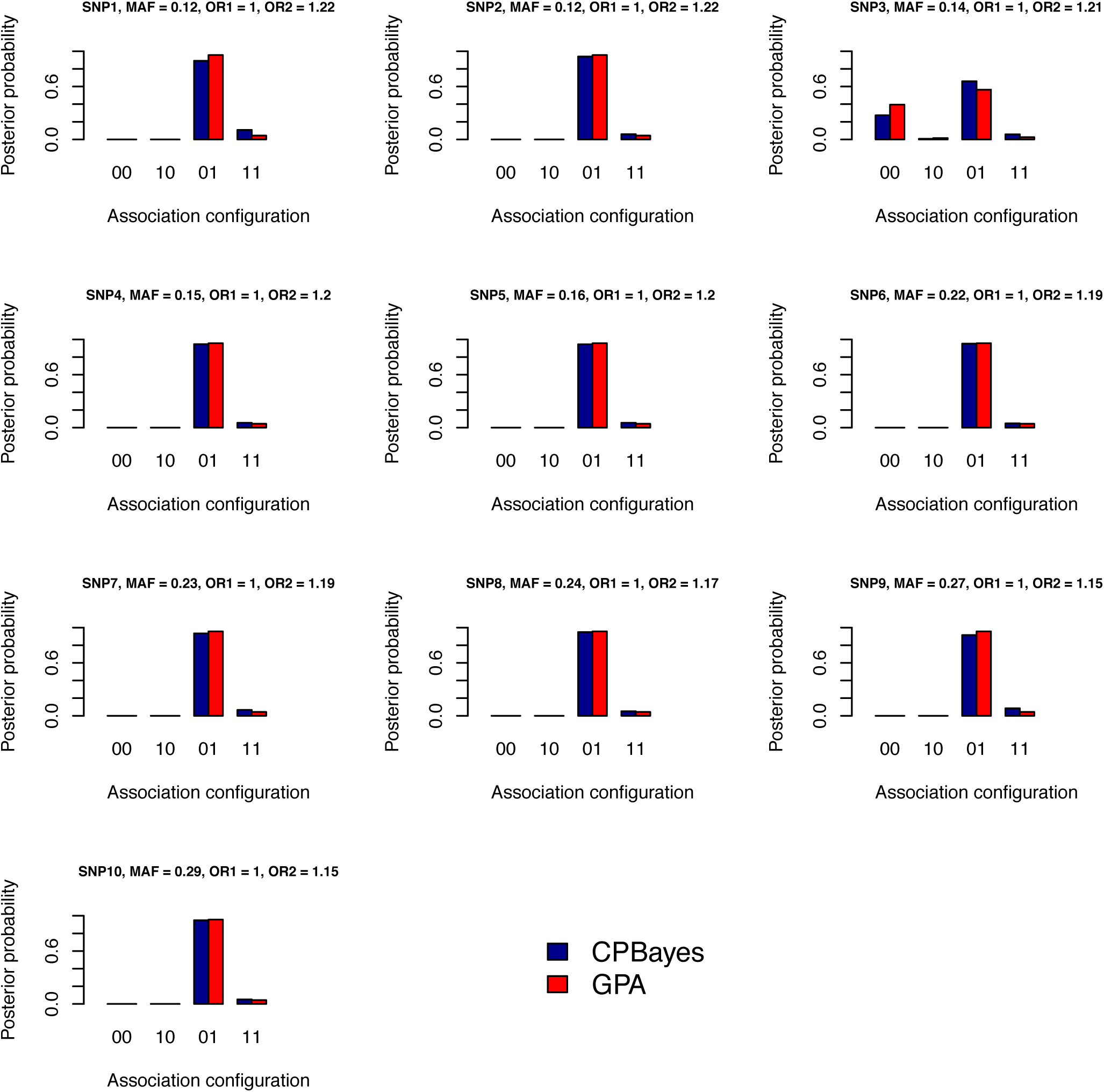
Estimated joint posterior probabilities of the trait configurations obtained by CPBayes and GPA for the first 10 risk SNPs. Here 2% of 1000 SNPs are risk SNPs and associated only with the second trait and 98% SNPs are null.

**Figure S10:**
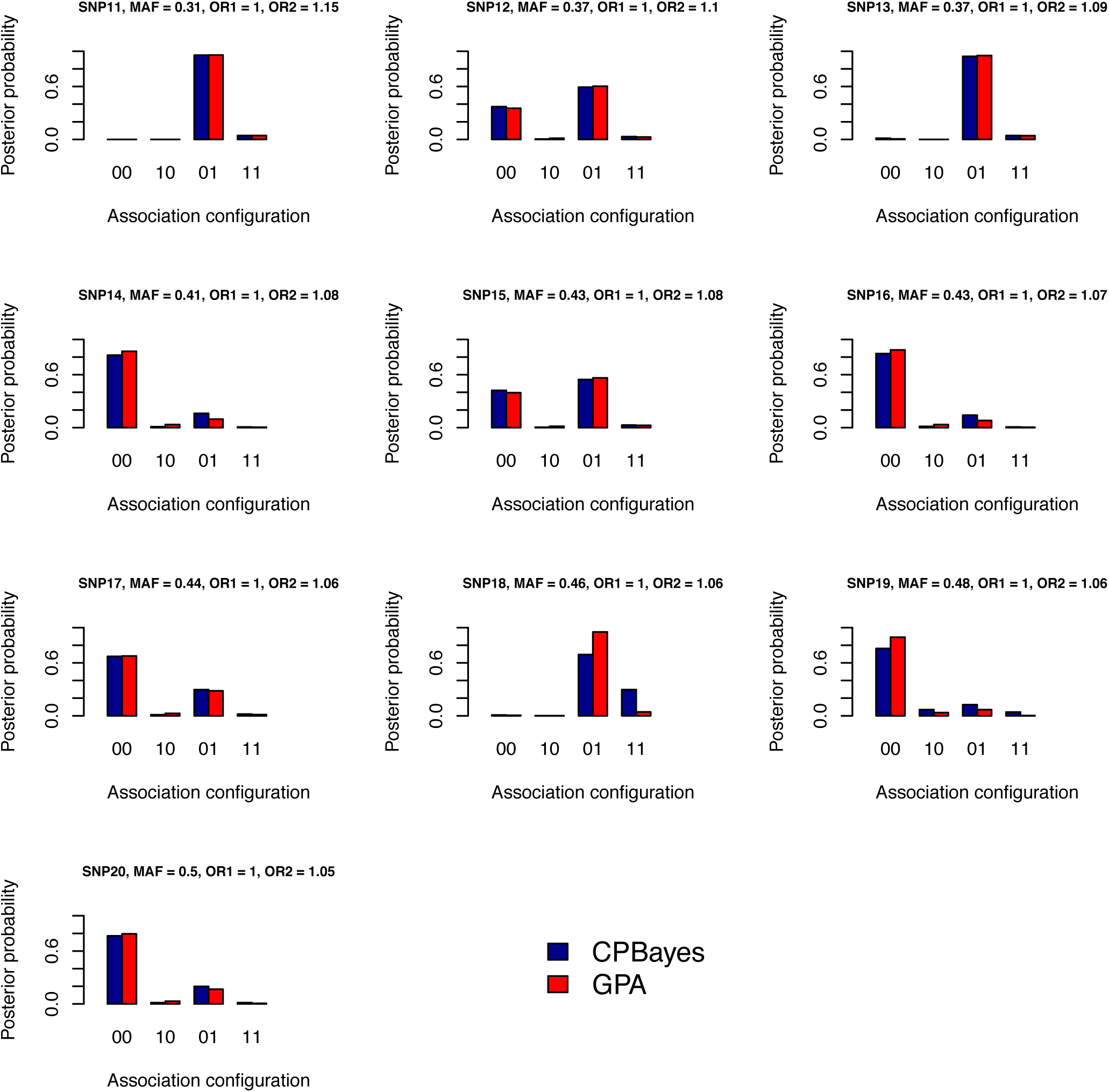
Estimated joint posterior probabilities of the trait configurations obtained by CPBayes and GPA for the last 10 risk SNPs. Here 2% of 1000 SNPs are risk SNPs and associated only with the second trait and 98% SNPs are null.

**Figure S11:**
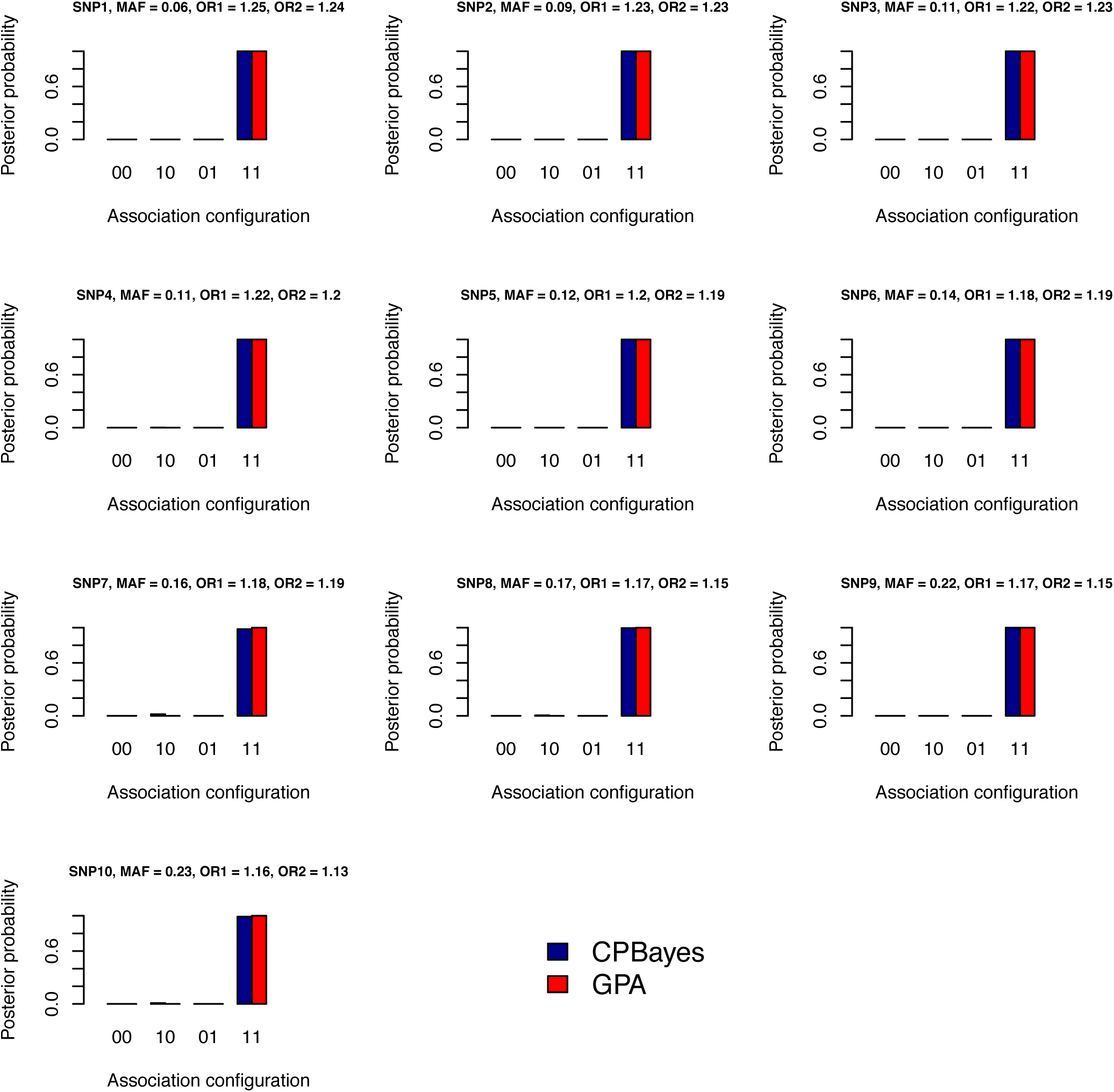
Estimated joint posterior probabilities of the trait configurations obtained by CPBayes and GPA for the first 10 risk SNPs. Here 2% of 1000 SNPs are risk SNPs and associated with both the traits and 98% SNPs are null.

**Figure S12:**
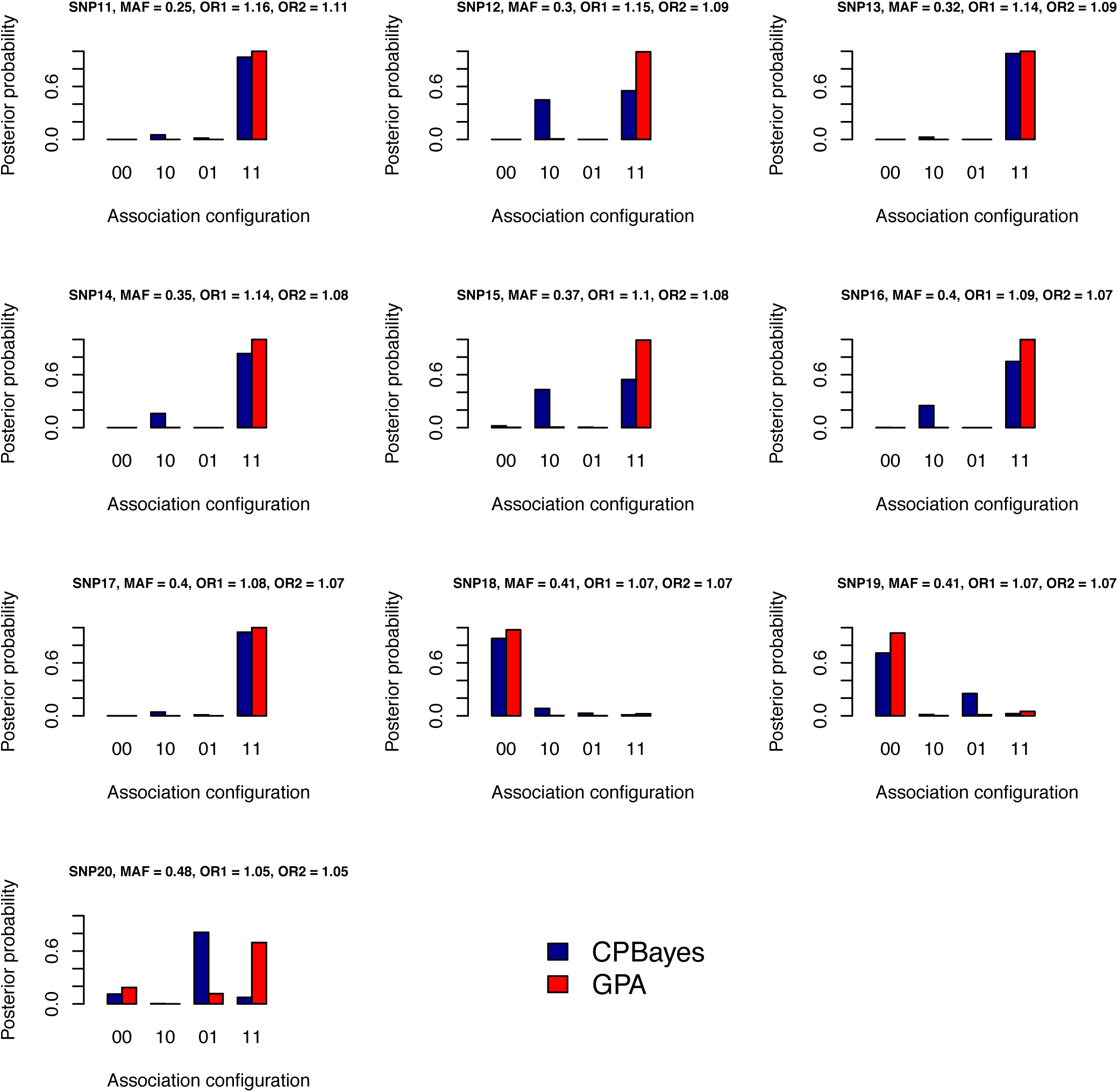
Estimated joint posterior probabilities of the trait configurations obtained by CPBayes and GPA for the last 10 risk SNPs. Here 2% of 1000 SNPs are risk SNPs and associated with both the traits and 98% SNPs are null.

## 7 Tables presenting the summaries of the measures of overall pleiotropic association under the global null hypothesis of no association

**Table S2:** Summary of measures for the evidence of the overall pleiotropic association under the global null hypothesis of no association when a cohort study with 15000 individuals is considered.

| K | m |  | mean | sd | min | quantile |  |  |  |  |  | max |
| --- | --- | --- | --- | --- | --- | --- | --- | --- | --- | --- | --- | --- |
|  |  |  |  |  |  | 5% | 25% | 50% | 75% | 95% | 99% |  |
| 5 | 0.3 | log<sub>10</sub>BF | -2.55 | 0.28 | -2.83 | -2.80 | -2.72 | -2.62 | -2.47 | -2.10 | -1.55 | -0.003 |
|  |  | -log<sub>10</sub>PPNA | 0.05 | 0.09 | 0.02 | 0.02 | 0.02 | 0.03 | 0.04 | 0.09 | 0.27 | 1.50 |
|  |  | -log<sub>10</sub>ASTpv | 0.36 | 0.53 | 0.00 | 0.004 | 0.06 | 0.20 | 0.45 | 1.21 | 2.32 | 6.20 |
|  | 0.1 | log<sub>10</sub>BF | -2.30 | 0.34 | -2.68 | -2.61 | -2.52 | -2.40 | -2.21 | -1.78 | -0.66 | -0.35 |
|  |  | -log<sub>10</sub>PPNA | 0.08 | 0.13 | 0.03 | 0.03 | 0.04 | 0.05 | 0.08 | 0.18 | 0.89 | 1.17 |
|  |  | -log<sub>10</sub>ASTpv | 0.36 | 0.45 | 0.00 | 0.002 | 0.06 | 0.20 | 0.50 | 1.28 | 2.13 | 3.48 |
| 10 | 0.3 | log<sub>10</sub>BF | -4.07 | 0.24 | -4.37 | -4.31 | -4.23 | -4.14 | -3.99 | -3.67 | -3.14 | -2.58 |
|  |  | -log<sub>10</sub>PPNA | 0.04 | 0.05 | 0.02 | 0.02 | 0.03 | 0.03 | 0.04 | 0.09 | 0.24 | 0.57 |
|  |  | -log<sub>10</sub>ASTpv | 0.18 | 0.50 | 0.00 | 0.00 | 0.00 | 0.02 | 0.14 | 0.80 | 2.34 | 4.72 |
|  | 0.1 | log<sub>10</sub>BF | -3.87 | 0.26 | -4.21 | -4.13 | -4.04 | -3.94 | -3.78 | -3.32 | -2.89 | -2.57 |
|  |  | -log<sub>10</sub>PPNA | 0.07 | 0.06 | 0.03 | 0.03 | 0.04 | 0.05 | 0.07 | 0.17 | 0.36 | 0.57 |
|  |  | -log<sub>10</sub>ASTpv | 0.15 | 0.31 | 0.00 | 0.00 | 0.00 | 0.02 | 0.15 | 0.72 | 1.69 | 2.12 |
| 15 | 0.3 | log<sub>10</sub>BF | -5.54 | 0.31 | -5.82 | -5.78 | -5.69 | -5.60 | -5.48 | -5.13 | -4.88 | -0.14 |
|  |  | -log<sub>10</sub>PPNA | 0.05 | 0.20 | 0.02 | 0.02 | 0.03 | 0.03 | 0.05 | 0.09 | 0.16 | 4.37 |
|  |  | -log<sub>10</sub>ASTpv | 0.07 | 0.50 | 0.00 | 0.00 | 0.00 | 0.00 | 0.00 | 0.19 | 1.38 | 8.27 |
|  | 0.1 | log<sub>10</sub>BF | -5.30 | 0.27 | -5.66 | -5.58 | -5.49 | -5.37 | -5.19 | -4.77 | -4.27 | -4.10 |
|  |  | -log<sub>10</sub>PPNA | 0.08 | 0.07 | 0.03 | 0.04 | 0.04 | 0.06 | 0.08 | 0.19 | 0.44 | 0.56 |
|  |  | -log<sub>10</sub>ASTpv | 0.06 | 0.25 | 0.00 | 0.00 | 0.00 | 0.00 | 0.01 | 0.27 | 1.11 | 3.28 |

Legend: $K$ - total number of phenotypes, $m$ - allele frequency at the marker SNP; $K = 5, 10, 15$ , and $m = 0.3, 0.1$ . The following abbreviations denote – log<sub>10</sub>BF: log<sub>10</sub>(Bayes factor), -log<sub>10</sub>PPNA: -log<sub>10</sub>(posterior probability of null association), -log<sub>10</sub>ASTpv: -log<sub>10</sub>(ASSET p-value). For a cohort study, since the summary statistics are correlated, the combined strategy of CPBayes is implemented. Different summary measures obtained across 500 replications are provided: mean, standard deviation (sd), minimum (min), 5% quantile, 25% quantile, 50% quantile, 75% quantile, 95% quantile, 99% quantile, and maximum (max).

## 8 Tables presenting the summaries of the measures of overall pleiotropic association under the global alternative hypothesis of association with one or more traits

**Table S3:** Summary of measures for the evidence of the overall pleiotropic association when a subset of traits are associated for **5 non-overlapping** case-control studies. Here **1** and **2** among **5** traits are associated.

| $K_1^+, K_1^-$ | $m$ | | mean | sd | min | quantile | | | | | | |
| --- | --- | --- | --- | --- | --- | --- | --- | --- | --- | --- | --- | --- |
|  |  |  |  |  |  | 5% | 25% | 50% | 75% | 95% | 99% | max |
| 1,0 | 0.3 | log<sub>10</sub>BF | 11.05 | 15.21 | -2.98 | -2.54 | -0.77 | 5.12 | 19.30 | 42.71 | 61.24 | 70.26 |
|  |  | -log<sub>10</sub>PPNA | 12.71 | 15.06 | 0.01 | 0.04 | 0.80 | 6.61 | 20.80 | 44.20 | 62.73 | 71.76 |
|  |  | -log<sub>10</sub>ASTpv | 7.73 | 5.74 | 0.06 | 0.82 | 2.97 | 6.44 | 11.68 | 18.36 | 23.98 | 26.34 |
|  | 0.1 | log<sub>10</sub>BF | 4.22 | 10.17 | -2.81 | -2.65 | -2.00 | -0.51 | 6.95 | 24.89 | 45.04 | 56.89 |
|  |  | -log<sub>10</sub>PPNA | 6.02 | 9.97 | 0.02 | 0.03 | 0.12 | 1.03 | 8.44 | 26.38 | 46.54 | 58.38 |
|  |  | -log<sub>10</sub>ASTpv | 3.48 | 2.83 | 0.01 | 0.25 | 1.13 | 2.75 | 5.16 | 8.73 | 11.92 | 14.65 |
| 2,0 | 0.3 | log<sub>10</sub>BF | 31.70 | 26.39 | -2.78 | -1.03 | 9.11 | 27.99 | 48.92 | 80.29 | 102.88 | 126.71 |
|  |  | -log<sub>10</sub>PPNA | 33.22 | 26.36 | 0.02 | 0.59 | 10.60 | 29.48 | 50.41 | 81.79 | 104.37 | 128.21 |
|  |  | -log<sub>10</sub>ASTpv | 15.27 | 7.92 | 0.23 | 3.44 | 9.27 | 14.67 | 20.05 | 29.14 | 34.48 | 43.27 |
|  | 0.1 | log<sub>10</sub>BF | 15.00 | 19.73 | -2.74 | -1.99 | -0.03 | 6.88 | 24.90 | 57.87 | 78.48 | 90.53 |
|  |  | -log<sub>10</sub>PPNA | 16.58 | 19.65 | 0.02 | 0.12 | 1.48 | 8.37 | 26.39 | 59.36 | 79.97 | 92.02 |
|  |  | -log<sub>10</sub>ASTpv | 6.54 | 3.84 | 0.07 | 1.28 | 3.24 | 6.08 | 9.06 | 13.53 | 16.44 | 19.03 |
| 1,1 | 0.3 | log<sub>10</sub>BF | 28.15 | 23.99 | -2.75 | -0.70 | 7.78 | 24.71 | 42.89 | 72.80 | 98.50 | 117.92 |
|  |  | -log<sub>10</sub>PPNA | 29.67 | 23.96 | 0.02 | 0.86 | 9.28 | 26.20 | 44.38 | 74.29 | 100.00 | 119.41 |
|  |  | -log<sub>10</sub>ASTpv | 15.41 | 7.92 | 0.57 | 3.82 | 9.20 | 14.96 | 20.57 | 29.46 | 36.45 | 41.72 |
|  | 0.1 | log<sub>10</sub>BF | 11.18 | 17.06 | -2.61 | -2.09 | -0.16 | 3.28 | 17.60 | 44.54 | 70.04 | 141.15 |
|  |  | -log<sub>10</sub>PPNA | 12.76 | 16.99 | 0.03 | 0.10 | 1.35 | 4.77 | 19.09 | 46.03 | 71.53 | 142.64 |
|  |  | -log<sub>10</sub>ASTpv | 6.32 | 3.63 | 0.19 | 1.31 | 3.76 | 5.61 | 8.63 | 12.94 | 16.11 | 24.97 |

Legend: $m$ - allele frequency at the risk SNP; $m = 0.3, 0.1$ . The number of positively and negatively associated traits are denoted by $K_1^+$ and $K_1^-$ , respectively. Hence the total number of associated traits is $K_1 = K_1^+ + K_1^-$ . The following abbreviations denote – log<sub>10</sub>BF: log<sub>10</sub>(Bayes factor), -log<sub>10</sub>PPNA: -log<sub>10</sub>(posterior probability of null association), -log<sub>10</sub>(ASTpv): -log<sub>10</sub>(ASSET p-value). For multiple studies with no overlapping subjects, the uncorrelated version of CPBayes is implemented. Different summary measures obtained across 500 replications are provided: mean, standard deviation (sd), minimum (min), 5% quantile, 25% quantile, 50% quantile, 75% quantile, 95% quantile, 99% quantile, and maximum (max).

**Table S4:** Summary of measures for the evidence of the overall pleiotropic association when a subset of traits are associated for **5 non-overlapping** case-control studies. Here **3** and **4** among **5** traits are associated.

| $K_1^+, K_1^-$ | $m$ | | mean | sd | min | quantile | | | | | | max |
| --- | --- | --- | --- | --- | --- | --- | --- | --- | --- | --- | --- | --- |
|  |  |  |  |  |  | 5% | 25% | 50% | 75% | 95% | 99% |  |
| 3,0 | 0.3 | log<sub>10</sub>BF | 61.61 | 40.01 | -2.44 | 4.61 | 30.29 | 58.08 | 85.48 | 134.62 | 175.19 | 197.55 |
|  |  | -log<sub>10</sub>PPNA | 63.11 | 40.00 | 0.05 | 6.11 | 31.78 | 59.57 | 86.97 | 136.11 | 176.68 | 199.04 |
|  |  | -log<sub>10</sub>ASTpv | 22.88 | 10.03 | 1.13 | 8.78 | 15.36 | 21.96 | 28.93 | 41.63 | 50.08 | 53.57 |
|  | 0.1 | log<sub>10</sub>BF | 33.38 | 34.50 | -2.37 | -0.84 | 3.62 | 24.34 | 52.62 | 99.66 | 142.34 | 171.82 |
|  |  | -log<sub>10</sub>PPNA | 34.89 | 34.48 | 0.05 | 0.74 | 5.11 | 25.83 | 54.11 | 101.15 | 143.83 | 173.31 |
|  |  | -log<sub>10</sub>ASTpv | 9.94 | 4.87 | 0.38 | 3.28 | 6.09 | 9.52 | 12.93 | 18.91 | 23.68 | 25.43 |
| 2,1 | 0.3 | log<sub>10</sub>BF | 60.31 | 39.41 | -2.07 | 4.80 | 29.29 | 56.15 | 84.89 | 131.91 | 170.04 | 201.46 |
|  |  | -log<sub>10</sub>PPNA | 61.80 | 39.41 | 0.10 | 6.29 | 30.79 | 57.64 | 86.38 | 133.40 | 171.53 | 202.95 |
|  |  | -log<sub>10</sub>ASTpv | 23.01 | 9.65 | 1.95 | 8.48 | 16.03 | 22.08 | 29.42 | 40.06 | 47.69 | 56.17 |
|  | 0.1 | log<sub>10</sub>BF | 30.24 | 31.41 | -2.60 | -1.22 | 3.58 | 21.99 | 47.77 | 90.33 | 129.23 | 188.29 |
|  |  | -log<sub>10</sub>PPNA | 31.76 | 31.39 | 0.03 | 0.46 | 5.07 | 23.48 | 49.26 | 91.82 | 130.72 | 189.78 |
|  |  | -log<sub>10</sub>ASTpv | 9.67 | 4.47 | 0.51 | 3.36 | 6.25 | 9.22 | 12.39 | 17.44 | 21.33 | 27.02 |
| 4,0 | 0.3 | log<sub>10</sub>BF | 95.17 | 48.22 | -0.33 | 24.89 | 59.37 | 91.99 | 128.33 | 179.22 | 214.40 | 245.43 |
|  |  | -log<sub>10</sub>PPNA | 96.66 | 48.22 | 1.19 | 26.38 | 60.86 | 93.48 | 129.82 | 180.71 | 215.89 | 246.92 |
|  |  | -log<sub>10</sub>ASTpv | 31.10 | 11.45 | 3.93 | 13.73 | 22.87 | 30.59 | 39.11 | 50.95 | 58.48 | 67.47 |
|  | 0.1 | log<sub>10</sub>BF | 56.83 | 46.68 | -2.16 | 0.80 | 19.87 | 48.05 | 80.55 | 146.62 | 213.33 | 234.68 |
|  |  | -log<sub>10</sub>PPNA | 58.32 | 46.68 | 0.08 | 2.30 | 21.36 | 49.54 | 82.04 | 148.12 | 214.82 | 236.17 |
|  |  | -log<sub>10</sub>ASTpv | 13.50 | 5.62 | 1.01 | 5.64 | 9.36 | 12.71 | 16.70 | 24.19 | 28.88 | 33.48 |
| 2,2 | 0.3 | log<sub>10</sub>BF | 88.32 | 44.33 | 0.31 | 23.17 | 55.83 | 86.42 | 114.73 | 168.46 | 192.95 | 224.88 |
|  |  | -log<sub>10</sub>PPNA | 89.81 | 44.33 | 1.81 | 24.66 | 57.32 | 87.91 | 116.22 | 169.95 | 194.44 | 226.37 |
|  |  | -log<sub>10</sub>ASTpv | 30.90 | 11.07 | 5.90 | 14.24 | 22.88 | 30.45 | 37.58 | 51.09 | 55.65 | 65.78 |
|  | 0.1 | log<sub>10</sub>BF | 51.67 | 41.90 | -1.73 | 0.39 | 17.85 | 43.79 | 77.10 | 127.42 | 176.04 | 257.33 |
|  |  | -log<sub>10</sub>PPNA | 53.16 | 41.89 | 0.20 | 1.88 | 19.34 | 45.28 | 78.59 | 128.91 | 177.53 | 258.83 |
|  |  | -log<sub>10</sub>ASTpv | 12.96 | 5.06 | 1.70 | 5.38 | 9.37 | 12.67 | 16.02 | 21.63 | 25.04 | 37.98 |

Legend: $m$ - allele frequency at the risk SNP; $m = 0.3, 0.1$ . The number of positively and negatively associated traits are denoted by $K_1^+$ and $K_1^-$ , respectively. Hence the total number of associated traits is $K_1 = K_1^+ + K_1^-$ . The following abbreviations denote - log<sub>10</sub>BF: log<sub>10</sub>(Bayes factor), -log<sub>10</sub>PPNA: -log<sub>10</sub>(posterior probability of null association), -log<sub>10</sub>(ASTpv): -log<sub>10</sub>(ASSET p-value). For multiple studies with no overlapping subjects, the uncorrelated version of CPBayes is implemented. Different summary measures obtained across 500 replications are provided: mean, standard deviation (sd), minimum (min), 5% quantile, 25% quantile, 50% quantile, 75% quantile, 95% quantile, 99% quantile, and maximum (max).

**Table S5:** Summary of measures for the evidence of the overall pleiotropic association when a subset of traits are associated for **10 non-overlapping** case-control studies. Here **2** and **4** among **10** traits are associated.

| $K_1^+, K_1^-$ | $m$ | | mean | sd | min | quantile | | | | | | max |
| --- | --- | --- | --- | --- | --- | --- | --- | --- | --- | --- | --- | --- |
|  |  |  |  |  |  | 5% | 25% | 50% | 75% | 95% | 99% |  |
| 2,0 | 0.3 | $\log_{10}\text{BF}$ | 27.15 | 25.74 | -4.30 | -2.69 | 5.25 | 21.92 | 43.93 | 75.80 | 101.52 | 129.36 |
| | | $-\log_{10}\text{PPNA}$ | 30.19 | 25.70 | 0.02 | 0.49 | 8.26 | 24.93 | 46.94 | 78.81 | 104.53 | 132.37 |
| | | $-\log_{10}\text{ASTpv}$ | 13.69 | 7.69 | 0.30 | 2.76 | 7.79 | 12.76 | 18.96 | 27.45 | 33.79 | 41.97 |
| | 0.1 | $\log_{10}\text{BF}$ | 11.17 | 21.28 | -4.34 | -3.79 | -2.10 | 1.72 | 15.69 | 63.79 | 86.82 | 112.94 |
| | | $-\log_{10}\text{PPNA}$ | 14.30 | 21.20 | 0.02 | 0.07 | 0.96 | 4.73 | 18.70 | 66.80 | 89.83 | 115.95 |
| | | $-\log_{10}\text{ASTpv}$ | 5.54 | 3.90 | 0.00 | 0.79 | 2.57 | 4.72 | 7.63 | 13.72 | 16.25 | 19.99 |
| 1,1 | 0.3 | $\log_{10}\text{BF}$ | 29.01 | 27.54 | -4.18 | -2.55 | 6.54 | 24.33 | 46.85 | 81.66 | 117.15 | 138.98 |
| | | $-\log_{10}\text{PPNA}$ | 32.06 | 27.50 | 0.03 | 0.59 | 9.55 | 27.34 | 49.86 | 84.67 | 120.16 | 141.99 |
| | | $-\log_{10}\text{ASTpv}$ | 13.48 | 7.47 | 0.56 | 2.75 | 7.71 | 13.09 | 18.61 | 26.26 | 34.03 | 40.35 |
| | 0.1 | $\log_{10}\text{BF}$ | 11.73 | 21.45 | -4.25 | -3.85 | -2.22 | 2.19 | 18.41 | 63.34 | 88.81 | 122.15 |
| | | $-\log_{10}\text{PPNA}$ | 14.88 | 21.35 | 0.02 | 0.06 | 0.85 | 5.20 | 21.42 | 66.35 | 91.82 | 125.16 |
| | | $-\log_{10}\text{ASTpv}$ | 5.25 | 3.54 | 0.05 | 0.61 | 2.67 | 4.77 | 6.98 | 12.29 | 15.35 | 19.13 |
| 4,0 | 0.3 | $\log_{10}\text{BF}$ | 89.89 | 47.02 | -3.04 | 17.36 | 56.24 | 88.18 | 117.85 | 174.97 | 214.41 | 257.88 |
| | | $-\log_{10}\text{PPNA}$ | 92.90 | 47.02 | 0.29 | 20.37 | 59.25 | 91.19 | 120.86 | 177.98 | 217.41 | 260.89 |
| | | $-\log_{10}\text{ASTpv}$ | 28.90 | 10.95 | 2.45 | 12.23 | 21.61 | 28.34 | 35.62 | 48.58 | 59.27 | 64.09 |
| | 0.1 | $\log_{10}\text{BF}$ | 48.12 | 44.63 | -3.91 | -2.02 | 9.87 | 38.46 | 72.24 | 133.95 | 162.01 | 208.74 |
| | | $-\log_{10}\text{PPNA}$ | 51.15 | 44.62 | 0.05 | 1.03 | 12.88 | 41.47 | 75.25 | 136.96 | 165.02 | 211.75 |
| | | $-\log_{10}\text{ASTpv}$ | 11.54 | 5.42 | 0.99 | 3.44 | 7.58 | 11.01 | 15.03 | 21.23 | 24.74 | 28.48 |
| 2,2 | 0.3 | $\log_{10}\text{BF}$ | 85.65 | 45.24 | -3.28 | 14.94 | 52.11 | 84.36 | 117.50 | 161.07 | 188.45 | 227.38 |
| | | $-\log_{10}\text{PPNA}$ | 88.66 | 45.24 | 0.19 | 17.95 | 55.12 | 87.37 | 120.51 | 164.08 | 191.46 | 230.39 |
| | | $-\log_{10}\text{ASTpv}$ | 28.33 | 10.51 | 2.58 | 12.07 | 20.62 | 27.62 | 34.88 | 46.53 | 52.30 | 58.24 |
| | 0.1 | $\log_{10}\text{BF}$ | 43.43 | 44.04 | -4.13 | -2.32 | 7.47 | 32.08 | 67.89 | 136.26 | 172.62 | 215.32 |
| | | $-\log_{10}\text{PPNA}$ | 46.45 | 44.02 | 0.03 | 0.77 | 10.48 | 35.09 | 70.90 | 139.27 | 175.63 | 218.33 |
| | | $-\log_{10}\text{ASTpv}$ | 10.80 | 5.12 | 0.09 | 3.64 | 7.04 | 10.20 | 14.12 | 20.77 | 24.02 | 27.71 |

Legend: $m$ - allele frequency at the risk SNP; $m = 0.3, 0.1$ . The number of positively and negatively associated traits are denoted by $K_1^+$ and $K_1^-$ , respectively. Hence the total number of associated traits is $K_1 = K_1^+ + K_1^-$ . The following abbreviations denote $-\log_{10}\text{BF}$ : $\log_{10}(\text{Bayes factor})$ , $-\log_{10}\text{PPNA}$ : $-\log_{10}(\text{posterior probability of null association})$ , $-\log_{10}(\text{ASTpv})$ : $-\log_{10}(\text{ASSET p-value})$ . For multiple studies with no overlapping subjects, the uncorrelated version of CPBayes is implemented. Different summary measures obtained across 500 replications are provided: mean, standard deviation (sd), minimum (min), 5% quantile, 25% quantile, 50% quantile, 75% quantile, 95% quantile, 99% quantile, and maximum (max).

**Table S6:** Summary of measures for the evidence of the overall pleiotropic association when a subset of traits are associated for **10 non-overlapping** case-control studies. Here **6** and **8** among **10** traits are associated.

| $K_1^+, K_1^-$ | $m$ | | mean | sd | min | quantile | | | | | | |
| --- | --- | --- | --- | --- | --- | --- | --- | --- | --- | --- | --- | --- |
|  |  |  |  |  |  | 5% | 25% | 50% | 75% | 95% | 99% | max |
| 6,0 | 0.3 | log<sub>10</sub>BF | 147.42 | 56.38 | -2.28 | 59.44 | 108.43 | 147.16 | 181.30 | 247.10 | 284.08 | 300.00 |
|  |  | -log<sub>10</sub>PPNA | 150.42 | 56.36 | 0.81 | 62.45 | 111.44 | 150.17 | 184.31 | 250.11 | 287.09 | 302.01 |
|  |  | -log<sub>10</sub>ASTpv | 43.75 | 13.70 | 4.92 | 23.89 | 33.58 | 42.92 | 51.90 | 66.37 | 78.13 | 100.85 |
|  | 0.1 | log<sub>10</sub>BF | 101.47 | 63.28 | -2.37 | 4.40 | 54.63 | 99.43 | 139.34 | 212.69 | 266.31 | 300.00 |
|  |  | -log<sub>10</sub>PPNA | 104.48 | 63.26 | 0.73 | 7.41 | 57.64 | 102.44 | 142.35 | 215.70 | 269.32 | 300.00 |
|  |  | -log<sub>10</sub>ASTpv | 18.21 | 6.92 | 2.45 | 7.46 | 13.29 | 17.83 | 22.36 | 30.50 | 35.63 | 39.72 |
| 3,3 | 0.3 | log<sub>10</sub>BF | 155.54 | 61.70 | 6.09 | 61.82 | 109.56 | 153.34 | 199.97 | 259.31 | 300.00 | 306.79 |
|  |  | -log<sub>10</sub>PPNA | 158.49 | 61.57 | 9.10 | 64.83 | 112.57 | 156.35 | 202.98 | 262.32 | 300.00 | 300.00 |
|  |  | -log<sub>10</sub>ASTpv | 43.29 | 13.66 | 10.44 | 22.95 | 33.37 | 42.07 | 52.78 | 64.85 | 75.52 | 107.79 |
|  | 0.1 | log<sub>10</sub>BF | 97.35 | 67.35 | -3.44 | 5.99 | 44.30 | 85.84 | 141.52 | 226.59 | 288.29 | 300.00 |
|  |  | -log<sub>10</sub>PPNA | 100.33 | 67.26 | 0.14 | 9.00 | 47.31 | 88.85 | 144.53 | 229.60 | 291.27 | 300.00 |
|  |  | -log<sub>10</sub>ASTpv | 17.17 | 6.66 | 2.14 | 7.99 | 12.28 | 16.12 | 21.33 | 28.82 | 35.29 | 44.07 |
| 8,0 | 0.3 | log<sub>10</sub>BF | 220.79 | 62.53 | -2.96 | 111.95 | 174.89 | 220.83 | 282.06 | 300.00 | 306.07 | 308.16 |
|  |  | -log<sub>10</sub>PPNA | 223.15 | 61.70 | 0.33 | 114.95 | 177.90 | 223.84 | 285.07 | 300.00 | 303.24 | 305.75 |
|  |  | -log<sub>10</sub>ASTpv | 59.89 | 16.60 | 6.50 | 34.85 | 48.36 | 57.74 | 71.32 | 90.44 | 99.24 | 116.55 |
|  | 0.1 | log<sub>10</sub>BF | 166.21 | 77.32 | -2.73 | 40.09 | 108.27 | 161.28 | 223.77 | 300.00 | 301.56 | 308.02 |
|  |  | -log<sub>10</sub>PPNA | 169.02 | 76.98 | 0.46 | 43.10 | 111.28 | 164.29 | 226.78 | 300.00 | 302.10 | 304.80 |
|  |  | -log<sub>10</sub>ASTpv | 25.39 | 8.28 | 2.97 | 13.27 | 19.34 | 24.44 | 30.72 | 39.65 | 46.85 | 66.42 |
| 4,4 | 0.3 | log<sub>10</sub>BF | 223.58 | 62.88 | 5.44 | 109.97 | 177.21 | 226.38 | 284.23 | 300.00 | 303.46 | 307.82 |
|  |  | -log<sub>10</sub>PPNA | 225.99 | 62.17 | 8.45 | 112.98 | 180.22 | 229.39 | 287.24 | 300.00 | 305.37 | 308.10 |
|  |  | -log<sub>10</sub>ASTpv | 58.72 | 16.15 | 12.15 | 34.11 | 47.51 | 56.49 | 68.94 | 87.91 | 97.65 | 107.58 |
|  | 0.1 | log<sub>10</sub>BF | 167.13 | 78.46 | -2.65 | 37.36 | 108.28 | 161.39 | 228.15 | 300.00 | 305.07 | 307.48 |
|  |  | -log<sub>10</sub>PPNA | 169.86 | 77.98 | 0.52 | 40.37 | 111.29 | 164.40 | 231.16 | 300.00 | 300.00 | 308.08 |
|  |  | -log<sub>10</sub>ASTpv | 24.25 | 7.62 | 4.21 | 13.03 | 18.68 | 23.23 | 29.24 | 37.93 | 44.33 | 50.27 |

Legend: $m$ - allele frequency at the risk SNP; $m = 0.3, 0.1$ . The number of positively and negatively associated traits are denoted by $K_1^+$ and $K_1^-$ , respectively. Hence the total number of associated traits is $K_1 = K_1^+ + K_1^-$ . The following abbreviations denote - log<sub>10</sub>BF: log<sub>10</sub>(Bayes factor), -log<sub>10</sub>PPNA: -log<sub>10</sub>(posterior probability of null association), -log<sub>10</sub>(ASTpv): -log<sub>10</sub>(ASSET p-value). For multiple studies with no overlapping subjects, the uncorrelated version of CPBayes is implemented. Different summary measures obtained across 500 replications are provided: mean, standard deviation (sd), minimum (min), 5% quantile, 25% quantile, 50% quantile, 75% quantile, 95% quantile, 99% quantile, and maximum (max).

**Table S7:** Summary of measures for the evidence of the overall pleiotropic association when a subset of traits are associated for **15 non-overlapping** case-control studies. Here 3, 6, and 9 among 15 traits are associated.

| $K_1^+, K_1^-$ | $m$ | | mean | sd | min | quantile | | | | | | |
| --- | --- | --- | --- | --- | --- | --- | --- | --- | --- | --- | --- | --- |
|  |  |  |  |  |  | 5% | 25% | 50% | 75% | 95% | 99% | max |
| 3,0 | 0.3 | $\log_{10}\text{BF}$ | 55.17 | 41.68 | -5.35 | -0.51 | 23.33 | 51.13 | 81.53 | 132.41 | 168.25 | 256.16 |
| | | $-\log_{10}\text{PPNA}$ | 59.69 | 41.68 | 0.06 | 4.01 | 27.85 | 55.64 | 86.05 | 136.92 | 172.77 | 260.68 |
| | | $-\log_{10}\text{ASTpv}$ | 20.09 | 10.16 | 1.57 | 5.88 | 12.41 | 19.06 | 26.12 | 38.63 | 48.88 | 66.07 |
| | 0.1 | $\log_{10}\text{BF}$ | 27.28 | 34.09 | -5.48 | -4.48 | 0.77 | 16.30 | 42.46 | 103.17 | 136.60 | 146.95 |
| | | $-\log_{10}\text{PPNA}$ | 31.83 | 34.05 | 0.05 | 0.32 | 5.29 | 20.82 | 46.98 | 107.69 | 141.12 | 151.46 |
| | | $-\log_{10}\text{ASTpv}$ | 7.84 | 4.49 | 0.22 | 1.69 | 4.49 | 7.31 | 10.18 | 16.73 | 20.07 | 22.00 |
| 2,1 | 0.3 | $\log_{10}\text{BF}$ | 51.39 | 37.55 | -5.32 | 0.31 | 21.42 | 48.76 | 73.17 | 120.28 | 154.40 | 185.80 |
| | | $-\log_{10}\text{PPNA}$ | 55.91 | 37.54 | 0.06 | 4.82 | 25.94 | 53.27 | 77.68 | 124.79 | 158.91 | 190.31 |
| | | $-\log_{10}\text{ASTpv}$ | 19.12 | 9.31 | 1.60 | 5.86 | 12.04 | 18.38 | 25.13 | 35.86 | 44.07 | 51.10 |
| | 0.1 | $\log_{10}\text{BF}$ | 25.10 | 31.59 | -5.58 | -4.74 | -0.81 | 12.28 | 44.58 | 84.75 | 119.30 | 167.94 |
| | | $-\log_{10}\text{PPNA}$ | 29.67 | 31.54 | 0.04 | 0.20 | 3.71 | 16.79 | 49.09 | 89.27 | 123.82 | 172.45 |
| | | $-\log_{10}\text{ASTpv}$ | 7.07 | 4.19 | 0.14 | 1.31 | 3.80 | 6.46 | 10.06 | 14.46 | 19.37 | 24.86 |
| 6,0 | 0.3 | $\log_{10}\text{BF}$ | 150.85 | 64.62 | 2.07 | 50.34 | 100.90 | 149.05 | 196.66 | 265.13 | 300.00 | 301.07 |
| | | $-\log_{10}\text{PPNA}$ | 155.29 | 64.46 | 6.58 | 54.86 | 105.41 | 153.57 | 201.17 | 269.64 | 300.00 | 305.59 |
| | | $-\log_{10}\text{ASTpv}$ | 42.15 | 14.59 | 11.11 | 20.04 | 31.75 | 40.78 | 51.27 | 67.53 | 80.81 | 96.92 |
| | 0.1 | $\log_{10}\text{BF}$ | 93.93 | 64.16 | -3.85 | 2.17 | 42.78 | 84.86 | 135.96 | 216.35 | 261.75 | 294.71 |
| | | $-\log_{10}\text{PPNA}$ | 98.44 | 64.16 | 0.75 | 6.69 | 47.29 | 89.38 | 140.48 | 220.87 | 266.26 | 299.22 |
| | | $-\log_{10}\text{ASTpv}$ | 16.90 | 6.52 | 4.34 | 7.10 | 12.14 | 16.19 | 20.84 | 28.50 | 33.52 | 35.63 |
| 3,3 | 0.3 | $\log_{10}\text{BF}$ | 148.37 | 62.97 | 7.85 | 53.78 | 102.07 | 146.29 | 186.63 | 261.24 | 298.32 | 306.15 |
| | | $-\log_{10}\text{PPNA}$ | 152.84 | 62.85 | 12.37 | 58.29 | 106.58 | 150.81 | 191.15 | 265.75 | 300.00 | 303.71 |
| | | $-\log_{10}\text{ASTpv}$ | 40.80 | 13.42 | 11.36 | 21.27 | 30.75 | 40.37 | 49.04 | 64.17 | 71.31 | 98.25 |
| | 0.1 | $\log_{10}\text{BF}$ | 87.73 | 64.91 | -4.57 | 0.54 | 36.33 | 78.09 | 124.58 | 215.44 | 266.81 | 302.95 |
| | | $-\log_{10}\text{PPNA}$ | 92.24 | 64.88 | 0.28 | 5.05 | 40.84 | 82.61 | 129.09 | 219.95 | 271.33 | 307.46 |
| | | $-\log_{10}\text{ASTpv}$ | 15.30 | 6.04 | 3.00 | 6.59 | 10.96 | 14.70 | 18.84 | 26.57 | 32.37 | 35.26 |
| 9,0 | 0.3 | $\log_{10}\text{BF}$ | 246.57 | 56.94 | 49.23 | 139.33 | 202.60 | 261.84 | 300.00 | 300.00 | 305.87 | 307.81 |
| | | $-\log_{10}\text{PPNA}$ | 249.49 | 55.45 | 53.74 | 143.85 | 207.11 | 266.36 | 300.00 | 300.00 | 303.63 | 308.15 |
| | | $-\log_{10}\text{ASTpv}$ | 65.43 | 17.91 | 21.04 | 39.79 | 52.46 | 64.45 | 76.44 | 97.32 | 111.35 | 132.49 |
| | 0.1 | $\log_{10}\text{BF}$ | 181.55 | 81.36 | -1.88 | 48.93 | 114.80 | 185.53 | 247.14 | 300.00 | 300.62 | 305.66 |
| | | $-\log_{10}\text{PPNA}$ | 185.47 | 80.49 | 2.63 | 53.45 | 119.31 | 190.04 | 251.66 | 300.00 | 303.71 | 307.81 |
| | | $-\log_{10}\text{ASTpv}$ | 26.31 | 8.63 | 6.98 | 13.68 | 20.04 | 25.99 | 31.70 | 41.36 | 49.88 | 57.39 |
| 5,4 | 0.3 | $\log_{10}\text{BF}$ | 244.60 | 56.81 | 41.08 | 138.93 | 206.04 | 260.15 | 300.00 | 300.00 | 302.91 | 307.72 |
| | | $-\log_{10}\text{PPNA}$ | 247.74 | 55.48 | 45.60 | 143.44 | 210.56 | 264.66 | 300.00 | 300.00 | 305.56 | 308.16 |
| | | $-\log_{10}\text{ASTpv}$ | 63.44 | 16.19 | 20.03 | 38.69 | 50.97 | 61.76 | 74.80 | 90.39 | 104.75 | 127.61 |
| | 0.1 | $\log_{10}\text{BF}$ | 176.98 | 81.15 | 1.87 | 40.71 | 116.37 | 173.77 | 243.61 | 300.00 | 303.21 | 308.20 |
| | | $-\log_{10}\text{PPNA}$ | 180.93 | 80.29 | 6.39 | 45.22 | 120.88 | 178.29 | 248.13 | 300.00 | 301.91 | 307.71 |
| | | $-\log_{10}\text{ASTpv}$ | 24.89 | 7.76 | 4.96 | 12.84 | 19.11 | 24.29 | 30.06 | 38.86 | 42.55 | 52.72 |

Legend: $m$ - allele frequency at the risk SNP; $m = 0.3, 0.1$ . The number of positively and negatively associated traits are denoted by $K_1^+$ and $K_1^-$ , respectively. Hence the total number of associated traits is $K_1 = K_1^+ + K_1^-$ . The following abbreviations denote $-\log_{10}\text{BF}$ : $\log_{10}(\text{Bayes factor})$ , $-\log_{10}\text{PPNA}$ : $-\log_{10}(\text{posterior probability of null association})$ , $-\log_{10}(\text{ASTpv})$ : $-\log_{10}(\text{ASSET p-value})$ . For multiple studies with no overlapping subjects, the uncorrelated version of CPBayes is implemented. Different summary measures obtained across 500 replications are provided: mean, standard deviation (sd), minimum (min), 5% quantile, 25% quantile, 50% quantile, 75% quantile, 95% quantile, 99% quantile, and maximum (max).

**Table S8:**
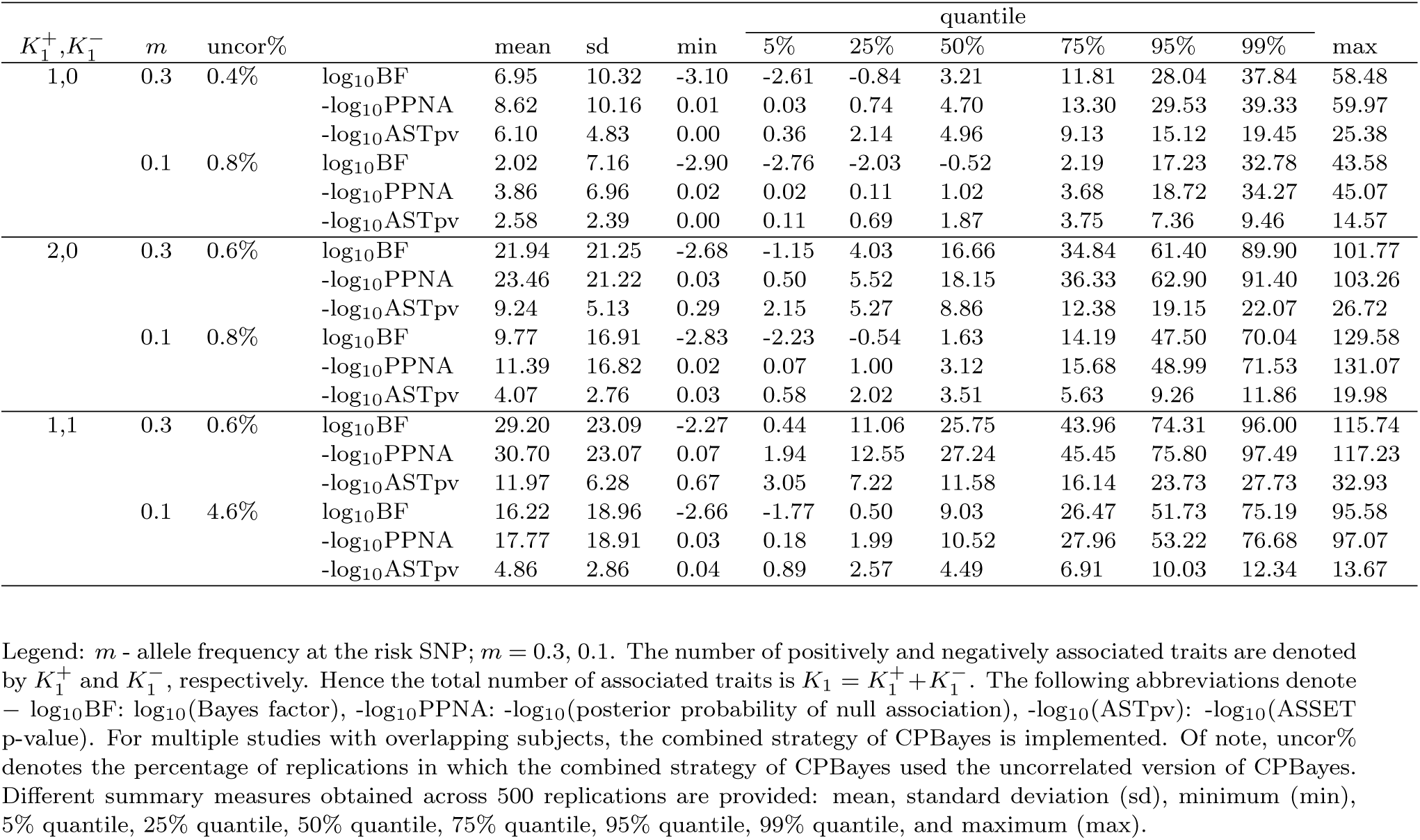
Summary of measures for the evidence of the overall pleiotropic association when a subset of traits are associated for **5 overlapping** case-control studies. Here **1** and **2** among **5** traits are associated.

**Table S9:** Summary of measures for the evidence of the overall pleiotropic association when a subset of traits are associated for **5 overlapping** case-control studies. Here **3** and **4** among **5** traits are associated.

| $K_1^+, K_1^-$ | $m$ | uncor% | | mean | sd | min | quantile | | | | | | | max |
| --- | --- | --- | --- | --- | --- | --- | --- | --- | --- | --- | --- | --- | --- | --- |
|  |  |  |  |  |  |  | 5% | 25% | 50% | 75% | 95% | 99% |  |  |
| 3,0 | 0.3 | 0.4% | log<sub>10</sub>BF | 34.94 | 28.81 | -2.55 | 0.14 | 12.46 | 28.52 | 51.75 | 87.75 | 119.47 | 161.65 |  |
|  |  |  | -log<sub>10</sub>PPNA | 36.45 | 28.80 | 0.04 | 1.64 | 13.95 | 30.01 | 53.24 | 89.24 | 120.96 | 163.15 |  |
|  |  |  | -log<sub>10</sub>ASTpv | 11.64 | 5.40 | 0.78 | 3.71 | 7.81 | 10.96 | 15.17 | 21.38 | 25.35 | 31.84 |  |
|  | 0.1 | 0.8% | log<sub>10</sub>BF | 15.32 | 22.10 | -2.73 | -2.04 | -0.10 | 5.92 | 23.74 | 60.50 | 88.52 | 139.69 |  |
|  |  |  | -log<sub>10</sub>PPNA | 16.90 | 22.03 | 0.02 | 0.11 | 1.41 | 7.41 | 25.23 | 61.99 | 90.01 | 141.18 |  |
|  |  |  | -log<sub>10</sub>ASTpv | 5.08 | 2.98 | 0.03 | 1.05 | 2.94 | 4.59 | 6.77 | 10.77 | 13.30 | 16.52 |  |
| 2,1 | 0.3 | 2.2% | log<sub>10</sub>BF | 52.74 | 31.83 | -2.01 | 7.77 | 28.42 | 49.47 | 72.46 | 112.17 | 141.18 | 156.62 |  |
|  |  |  | -log<sub>10</sub>PPNA | 54.24 | 31.83 | 0.12 | 9.27 | 29.91 | 50.96 | 73.95 | 113.66 | 142.67 | 158.11 |  |
|  |  |  | -log<sub>10</sub>ASTpv | 15.68 | 6.63 | 1.21 | 5.67 | 10.92 | 15.00 | 20.00 | 27.83 | 33.05 | 35.93 |  |
|  | 0.1 | 5.8% | log<sub>10</sub>BF | 32.79 | 29.50 | -2.78 | -0.93 | 6.50 | 26.35 | 52.58 | 88.01 | 113.65 | 122.83 |  |
|  |  |  | -log<sub>10</sub>PPNA | 34.30 | 29.48 | 0.02 | 0.66 | 8.00 | 27.84 | 54.08 | 89.50 | 115.14 | 124.33 |  |
|  |  |  | -log<sub>10</sub>ASTpv | 6.45 | 3.21 | 0.02 | 2.00 | 3.93 | 6.16 | 8.59 | 11.99 | 14.84 | 16.27 |  |
| 4,0 | 0.3 | 1 | log<sub>10</sub>BF | 40.37 | 32.14 | -2.32 | 0.74 | 14.01 | 35.31 | 59.80 | 98.97 | 127.94 | 169.53 |  |
|  |  |  | -log<sub>10</sub>PPNA | 41.87 | 32.13 | 0.06 | 2.24 | 15.50 | 36.80 | 61.29 | 100.46 | 129.43 | 171.03 |  |
|  |  |  | -log<sub>10</sub>ASTpv | 13.67 | 5.53 | 0.76 | 5.38 | 9.76 | 12.96 | 17.49 | 23.38 | 28.23 | 31.19 |  |
|  | 0.1 | 3.6 | log<sub>10</sub>BF | 14.98 | 24.90 | -2.65 | -1.94 | -0.29 | 3.55 | 21.02 | 69.20 | 105.87 | 151.60 |  |
|  |  |  | -log<sub>10</sub>PPNA | 16.56 | 24.84 | 0.03 | 0.13 | 1.23 | 5.05 | 22.51 | 70.69 | 107.36 | 153.09 |  |
|  |  |  | -log<sub>10</sub>ASTpv | 5.94 | 3.19 | 0.35 | 1.54 | 3.61 | 5.56 | 7.85 | 12.20 | 14.84 | 17.87 |  |
| 2,2 | 0.3 | 1 | log<sub>10</sub>BF | 82.12 | 37.92 | -0.81 | 24.63 | 53.20 | 78.45 | 109.34 | 143.83 | 178.97 | 195.74 |  |
|  |  |  | -log<sub>10</sub>PPNA | 83.61 | 37.92 | 0.76 | 26.12 | 54.70 | 79.95 | 110.83 | 145.32 | 180.46 | 197.24 |  |
|  |  |  | -log<sub>10</sub>ASTpv | 19.07 | 6.56 | 2.14 | 8.70 | 14.05 | 18.80 | 23.69 | 30.08 | 33.76 | 37.17 |  |
|  | 0.1 | 6.8 | log<sub>10</sub>BF | 53.77 | 35.96 | -1.82 | 0.89 | 25.71 | 49.63 | 78.00 | 123.01 | 135.74 | 151.22 |  |
|  |  |  | -log<sub>10</sub>PPNA | 55.26 | 35.95 | 0.17 | 2.39 | 27.20 | 51.12 | 79.49 | 124.50 | 137.23 | 152.71 |  |
|  |  |  | -log<sub>10</sub>ASTpv | 7.74 | 3.03 | 0.80 | 3.17 | 5.54 | 7.29 | 9.91 | 12.92 | 15.16 | 16.92 |  |

Legend: $m$ - allele frequency at the risk SNP; $m = 0.3, 0.1$ . The number of positively and negatively associated traits are denoted by $K_1^+$ and $K_1^-$ , respectively. Hence the total number of associated traits is $K_1 = K_1^+ + K_1^-$ . The following abbreviations denote - log<sub>10</sub>BF: log<sub>10</sub>(Bayes factor), -log<sub>10</sub>PPNA: -log<sub>10</sub>(posterior probability of null association), -log<sub>10</sub>(ASTpv): -log<sub>10</sub>(ASSET p-value). For multiple studies with overlapping subjects, the combined strategy of CPBayes is implemented. Of note, uncor% denotes the percentage of replications in which the combined strategy of CPBayes used the uncorrelated version of CPBayes. Different summary measures obtained across 500 replications are provided: mean, standard deviation (sd), minimum (min), 5% quantile, 25% quantile, 50% quantile, 75% quantile, 95% quantile, 99% quantile, and maximum (max).

**Table S10:** Summary of measures for the evidence of the overall pleiotropic association when a subset of traits are associated for **10 overlapping** case-control studies. Here **2** and **4** among **10** traits are associated.

| $K_1^+, K_1^-$ | $m$ | uncor% | | mean | sd | min | quantile | | | | | | | max |
| --- | --- | --- | --- | --- | --- | --- | --- | --- | --- | --- | --- | --- | --- | --- |
|  |  |  |  |  |  |  | 5% | 25% | 50% | 75% | 95% | 99% |  |  |
| 2,0 | 0.3 | 0.2% | log<sub>10</sub>BF | 22.76 | 22.21 | -4.25 | -2.54 | 4.48 | 17.93 | 37.13 | 65.19 | 87.68 | 113.37 |  |
|  |  |  | -log<sub>10</sub>PPNA | 25.80 | 22.17 | 0.02 | 0.60 | 7.49 | 20.94 | 40.14 | 68.20 | 90.69 | 116.38 |  |
|  |  |  | -log<sub>10</sub>ASTpv | 7.93 | 4.91 | 0.00 | 0.99 | 4.12 | 7.38 | 11.04 | 16.83 | 21.79 | 26.42 |  |
|  | 0.1 | 2.4% | log<sub>10</sub>BF | 9.70 | 17.03 | -4.32 | -3.86 | -2.07 | 3.30 | 15.21 | 47.44 | 72.03 | 93.28 |  |
|  |  |  | -log<sub>10</sub>PPNA | 12.84 | 16.92 | 0.02 | 0.06 | 0.99 | 6.31 | 18.22 | 50.45 | 75.04 | 96.29 |  |
|  |  |  | -log<sub>10</sub>ASTpv | 2.92 | 2.57 | 0.00 | 0.03 | 0.91 | 2.38 | 4.24 | 7.53 | 12.19 | 14.01 |  |
| 1,1 | 0.3 | 1% | log<sub>10</sub>BF | 27.39 | 22.87 | -4.08 | -2.22 | 8.91 | 23.86 | 41.70 | 71.44 | 87.53 | 117.80 |  |
|  |  |  | -log<sub>10</sub>PPNA | 30.43 | 22.84 | 0.04 | 0.85 | 11.92 | 26.87 | 44.71 | 74.45 | 90.54 | 120.81 |  |
|  |  |  | -log<sub>10</sub>ASTpv | 10.07 | 6.08 | 0.02 | 1.18 | 5.12 | 9.55 | 13.97 | 21.60 | 24.71 | 29.81 |  |
|  | 0.1 | 3.8% | log<sub>10</sub>BF | 13.86 | 19.58 | -4.39 | -3.75 | -1.16 | 6.53 | 22.99 | 52.39 | 82.73 | 105.61 |  |
|  |  |  | -log<sub>10</sub>PPNA | 16.97 | 19.49 | 0.02 | 0.07 | 1.86 | 9.54 | 26.00 | 55.40 | 85.74 | 108.62 |  |
|  |  |  | -log<sub>10</sub>ASTpv | 3.28 | 2.58 | 0.00 | 0.07 | 1.22 | 2.78 | 4.92 | 8.47 | 10.73 | 12.49 |  |
| 4,0 | 0.3 | 0.6% | log<sub>10</sub>BF | 57.66 | 38.31 | -3.66 | 3.74 | 28.53 | 53.68 | 81.07 | 132.47 | 157.92 | 183.85 |  |
|  |  |  | -log<sub>10</sub>PPNA | 60.67 | 38.31 | 0.09 | 6.75 | 31.54 | 56.69 | 84.08 | 135.48 | 160.93 | 186.85 |  |
|  |  |  | -log<sub>10</sub>ASTpv | 11.73 | 5.22 | 0.34 | 4.46 | 8.02 | 11.03 | 14.89 | 21.50 | 25.96 | 30.74 |  |
|  | 0.1 | 3.6% | log<sub>10</sub>BF | 25.97 | 31.75 | -3.81 | -2.76 | 0.36 | 14.82 | 40.68 | 93.18 | 122.71 | 161.92 |  |
|  |  |  | -log<sub>10</sub>PPNA | 29.01 | 31.73 | 0.06 | 0.44 | 3.37 | 17.83 | 43.69 | 96.19 | 125.72 | 164.93 |  |
|  |  |  | -log<sub>10</sub>ASTpv | 4.41 | 2.84 | 0.02 | 0.72 | 2.28 | 4.03 | 5.88 | 9.77 | 13.47 | 15.66 |  |
| 2,2 | 0.3 | 4.8% | log<sub>10</sub>BF | 77.81 | 40.39 | -1.34 | 18.29 | 50.34 | 72.52 | 100.73 | 152.82 | 172.32 | 199.91 |  |
|  |  |  | -log<sub>10</sub>PPNA | 80.82 | 40.39 | 1.68 | 21.30 | 53.35 | 75.53 | 103.74 | 155.83 | 175.33 | 202.92 |  |
|  |  |  | -log<sub>10</sub>ASTpv | 16.80 | 6.35 | 1.63 | 6.71 | 12.27 | 16.17 | 21.39 | 28.00 | 30.75 | 32.92 |  |
|  | 0.1 | 12.4% | log<sub>10</sub>BF | 46.82 | 36.71 | -3.32 | -1.54 | 18.64 | 39.66 | 70.42 | 114.56 | 141.42 | 179.66 |  |
|  |  |  | -log<sub>10</sub>PPNA | 49.84 | 36.69 | 0.17 | 1.48 | 21.65 | 42.67 | 73.43 | 117.57 | 144.43 | 182.67 |  |
|  |  |  | -log<sub>10</sub>ASTpv | 5.88 | 2.76 | 0.08 | 1.56 | 3.76 | 5.71 | 7.80 | 10.59 | 12.33 | 14.16 |  |

Legend: $m$ - allele frequency at the risk SNP; $m = 0.3, 0.1$ . The number of positively and negatively associated traits are denoted by $K_1^+$ and $K_1^-$ , respectively. Hence the total number of associated traits is $K_1 = K_1^+ + K_1^-$ . The following abbreviations denote - log<sub>10</sub>BF: log<sub>10</sub>(Bayes factor), -log<sub>10</sub>PPNA: -log<sub>10</sub>(posterior probability of null association), -log<sub>10</sub>(ASTpv): -log<sub>10</sub>(ASSET p-value). For multiple studies with overlapping subjects, the combined strategy of CPBayes is implemented. Of note, uncor% denotes the percentage of replications in which the combined strategy of CPBayes used the uncorrelated version of CPBayes. Different summary measures obtained across 500 replications are provided: mean, standard deviation (sd), minimum (min), 5% quantile, 25% quantile, 50% quantile, 75% quantile, 95% quantile, 99% quantile, and maximum (max).

**Table S11:** Summary of measures for the evidence of the overall pleiotropic association when a subset of traits are associated for **10 overlapping** case-control studies. Here **6** and **8** among **10** traits are associated.

| $K_1^+, K_1^-$ | $m$ | uncor% | | mean | sd | min | quantile | | | | | | | max |
| --- | --- | --- | --- | --- | --- | --- | --- | --- | --- | --- | --- | --- | --- | --- |
|  |  |  |  |  |  |  | 5% | 25% | 50% | 75% | 95% | 99% |  |  |
| 6,0 | 0.3 | 3% | log<sub>10</sub>BF | 83.92 | 49.08 | -3.71 | 12.94 | 48.13 | 79.33 | 116.43 | 170.26 | 224.05 | 265.78 |  |
|  |  |  | -log<sub>10</sub>PPNA | 86.93 | 49.08 | 0.08 | 15.95 | 51.14 | 82.34 | 119.44 | 173.27 | 227.06 | 268.79 |  |
|  |  |  | -log<sub>10</sub>ASTpv | 14.06 | 5.37 | 0.94 | 6.58 | 10.02 | 13.72 | 17.25 | 24.07 | 27.59 | 34.60 |  |
|  | 0.1 | 8.6% | log<sub>10</sub>BF | 35.46 | 42.51 | -3.52 | -2.38 | 3.21 | 21.12 | 53.48 | 122.70 | 162.39 | 261.30 |  |
|  |  |  | -log<sub>10</sub>PPNA | 38.48 | 42.50 | 0.12 | 0.72 | 6.22 | 24.13 | 56.49 | 125.71 | 165.40 | 264.31 |  |
|  |  |  | -log<sub>10</sub>ASTpv | 5.41 | 3.03 | 0.03 | 1.52 | 3.07 | 4.91 | 7.07 | 11.41 | 14.02 | 17.92 |  |
| 3,3 | 0.3 | 6.8% | log<sub>10</sub>BF | 139.07 | 53.94 | 8.81 | 55.16 | 101.18 | 135.69 | 173.47 | 230.87 | 289.54 | 300.51 |  |
|  |  |  | -log<sub>10</sub>PPNA | 142.07 | 53.90 | 11.82 | 58.17 | 104.19 | 138.70 | 176.48 | 233.88 | 292.55 | 303.52 |  |
|  |  |  | -log<sub>10</sub>ASTpv | 21.21 | 6.49 | 5.19 | 11.23 | 16.48 | 20.85 | 25.65 | 32.16 | 36.14 | 39.79 |  |
|  | 0.1 | 14.8% | log<sub>10</sub>BF | 97.04 | 57.63 | -2.57 | 4.98 | 53.47 | 92.07 | 136.05 | 200.53 | 234.69 | 280.41 |  |
|  |  |  | -log<sub>10</sub>PPNA | 100.05 | 57.63 | 0.58 | 7.99 | 56.48 | 95.08 | 139.06 | 203.54 | 237.70 | 283.42 |  |
|  |  |  | -log<sub>10</sub>ASTpv | 7.84 | 3.02 | 0.92 | 3.21 | 5.69 | 7.69 | 9.84 | 12.86 | 15.05 | 18.50 |  |
| 8,0 | 0.3 | 6.4% | log<sub>10</sub>BF | 90.62 | 59.51 | -3.79 | 11.84 | 45.35 | 79.68 | 123.46 | 205.05 | 266.41 | 300.00 |  |
|  |  |  | -log<sub>10</sub>PPNA | 93.63 | 59.49 | 0.07 | 14.85 | 48.36 | 82.69 | 126.47 | 208.06 | 269.42 | 300.00 |  |
|  |  |  | -log<sub>10</sub>ASTpv | 16.28 | 5.76 | 0.77 | 8.38 | 12.13 | 15.56 | 19.73 | 26.55 | 32.47 | 34.89 |  |
|  | 0.1 | 21.8% | log<sub>10</sub>BF | 39.87 | 52.80 | -3.99 | -2.85 | -0.56 | 19.04 | 65.18 | 143.11 | 222.08 | 301.12 |  |
|  |  |  | -log<sub>10</sub>PPNA | 42.91 | 52.78 | 0.04 | 0.39 | 2.45 | 22.05 | 68.19 | 146.12 | 225.09 | 304.13 |  |
|  |  |  | -log<sub>10</sub>ASTpv | 6.41 | 3.40 | 0.05 | 1.88 | 3.81 | 5.84 | 8.43 | 12.73 | 15.66 | 18.32 |  |
| 4,4 | 0.3 | 7% | log<sub>10</sub>BF | 201.68 | 59.53 | 30.89 | 103.30 | 159.75 | 200.35 | 245.62 | 300.00 | 303.97 | 308.11 |  |
|  |  |  | -log<sub>10</sub>PPNA | 204.45 | 59.13 | 33.90 | 106.31 | 162.76 | 203.36 | 248.63 | 300.00 | 303.90 | 307.17 |  |
|  |  |  | -log<sub>10</sub>ASTpv | 24.99 | 6.41 | 5.69 | 14.63 | 20.24 | 25.42 | 29.21 | 35.53 | 39.73 | 43.02 |  |
|  | 0.1 | 23.4% | log<sub>10</sub>BF | 153.58 | 75.61 | -2.99 | 25.13 | 98.67 | 157.58 | 208.83 | 279.24 | 300.00 | 304.04 |  |
|  |  |  | -log<sub>10</sub>PPNA | 156.52 | 75.48 | 0.31 | 28.14 | 101.68 | 160.59 | 211.84 | 282.25 | 300.00 | 307.05 |  |
|  |  |  | -log<sub>10</sub>ASTpv | 9.49 | 3.05 | 1.02 | 4.55 | 7.33 | 9.46 | 11.69 | 14.54 | 16.51 | 17.63 |  |

Legend: $m$ - allele frequency at the risk SNP; $m = 0.3, 0.1$ . The number of positively and negatively associated traits are denoted by $K_1^+$ and $K_1^-$ , respectively. Hence the total number of associated traits is $K_1 = K_1^+ + K_1^-$ . The following abbreviations denote - log<sub>10</sub>BF: log<sub>10</sub>(Bayes factor), -log<sub>10</sub>PPNA: -log<sub>10</sub>(posterior probability of null association), -log<sub>10</sub>(ASTpv): -log<sub>10</sub>(ASSET p-value). For multiple studies with overlapping subjects, the combined strategy of CPBayes is implemented. Of note, uncor% denotes the percentage of replications in which the combined strategy of CPBayes used the uncorrelated version of CPBayes. Different summary measures obtained across 500 replications are provided: mean, standard deviation (sd), minimum (min), 5% quantile, 25% quantile, 50% quantile, 75% quantile, 95% quantile, 99% quantile, and maximum (max).

**Table S12:** Summary of measures for the evidence of the overall pleiotropic association when a subset of traits are associated for **15 overlapping** case-control studies. Here 3,6, and 9 among 15 traits are associated.

| $K_1^+, K_1^-$ | $m$ | uncor% | | mean | sd | min | quantile | | | | | | | max |
| --- | --- | --- | --- | --- | --- | --- | --- | --- | --- | --- | --- | --- | --- | --- |
|  |  |  |  |  |  |  | 5% | 25% | 50% | 75% | 95% | 99% |  |  |
| 3,0 | 0.3 | 2% | log<sub>10</sub>BF | 40.22 | 30.92 | -5.56 | -1.62 | 15.82 | 35.76 | 58.47 | 99.67 | 123.13 | 144.36 |  |
|  |  |  | -log<sub>10</sub>PPNA | 44.75 | 30.91 | 0.04 | 2.89 | 20.34 | 40.27 | 62.99 | 104.19 | 127.65 | 148.88 |  |
|  |  |  | -log<sub>10</sub>ASTpv | 9.06 | 4.79 | 0.00 | 1.84 | 5.87 | 8.65 | 11.75 | 17.76 | 21.93 | 27.17 |  |
|  | 0.1 | 3.6% | log<sub>10</sub>BF | 19.65 | 26.51 | -5.70 | -4.71 | -2.00 | 10.42 | 31.76 | 72.91 | 106.83 | 151.34 |  |
|  |  |  | -log<sub>10</sub>PPNA | 24.22 | 26.46 | 0.03 | 0.21 | 2.51 | 14.94 | 36.27 | 77.42 | 111.34 | 155.86 |  |
|  |  |  | -log<sub>10</sub>ASTpv | 3.00 | 2.58 | 0.00 | 0.01 | 1.08 | 2.50 | 4.08 | 7.83 | 11.60 | 16.89 |  |
| 2,1 | 0.3 | 5% | log<sub>10</sub>BF | 47.03 | 32.44 | -5.39 | 2.96 | 23.44 | 41.37 | 66.60 | 110.51 | 132.43 | 153.69 |  |
|  |  |  | -log<sub>10</sub>PPNA | 51.55 | 32.44 | 0.05 | 7.47 | 27.96 | 45.89 | 71.11 | 115.02 | 136.94 | 158.20 |  |
|  |  |  | -log<sub>10</sub>ASTpv | 11.83 | 6.32 | 0.00 | 2.74 | 6.86 | 11.11 | 16.08 | 23.60 | 27.04 | 30.20 |  |
|  | 0.1 | 11% | log<sub>10</sub>BF | 24.17 | 28.34 | -5.64 | -4.58 | 0.24 | 16.58 | 41.62 | 76.20 | 112.88 | 147.56 |  |
|  |  |  | -log<sub>10</sub>PPNA | 28.73 | 28.30 | 0.03 | 0.27 | 4.76 | 21.10 | 46.14 | 80.72 | 117.39 | 152.07 |  |
|  |  |  | -log<sub>10</sub>ASTpv | 3.26 | 2.56 | 0.00 | 0.04 | 1.11 | 2.90 | 5.05 | 7.78 | 10.58 | 12.66 |  |
| 6,0 | 0.3 | 3.2% | log<sub>10</sub>BF | 98.03 | 51.34 | -4.30 | 22.72 | 61.20 | 92.73 | 132.10 | 186.17 | 226.55 | 249.79 |  |
|  |  |  | -log<sub>10</sub>PPNA | 102.55 | 51.34 | 0.42 | 27.23 | 65.71 | 97.25 | 136.62 | 190.69 | 231.07 | 254.31 |  |
|  |  |  | -log<sub>10</sub>ASTpv | 13.00 | 5.03 | 1.41 | 5.77 | 9.33 | 12.38 | 16.18 | 22.16 | 27.71 | 29.63 |  |
|  | 0.1 | 7.4% | log<sub>10</sub>BF | 50.67 | 48.04 | -5.20 | -3.52 | 11.23 | 38.68 | 79.61 | 143.06 | 190.66 | 228.90 |  |
|  |  |  | -log<sub>10</sub>PPNA | 55.20 | 48.03 | 0.08 | 1.04 | 15.74 | 43.20 | 84.12 | 147.57 | 195.17 | 233.42 |  |
|  |  |  | -log<sub>10</sub>ASTpv | 4.40 | 2.87 | 0.00 | 0.85 | 2.40 | 3.88 | 5.78 | 10.21 | 13.85 | 18.72 |  |
| 3,3 | 0.3 | 9.6% | log<sub>10</sub>BF | 131.30 | 55.63 | 2.77 | 40.71 | 94.71 | 124.78 | 167.57 | 228.71 | 254.75 | 300.00 |  |
|  |  |  | -log<sub>10</sub>PPNA | 135.81 | 55.60 | 7.28 | 45.23 | 99.22 | 129.29 | 172.09 | 233.23 | 259.26 | 300.00 |  |
|  |  |  | -log<sub>10</sub>ASTpv | 19.19 | 6.29 | 2.98 | 8.81 | 14.82 | 18.85 | 23.33 | 30.05 | 32.90 | 39.04 |  |
|  | 0.1 | 18% | log<sub>10</sub>BF | 88.17 | 56.96 | -4.76 | 2.81 | 43.27 | 81.75 | 128.31 | 192.61 | 238.06 | 280.95 |  |
|  |  |  | -log<sub>10</sub>PPNA | 92.69 | 56.95 | 0.20 | 7.32 | 47.78 | 86.26 | 132.83 | 197.13 | 242.58 | 285.47 |  |
|  |  |  | -log<sub>10</sub>ASTpv | 6.20 | 2.80 | 0.42 | 1.88 | 4.13 | 6.06 | 8.13 | 10.84 | 12.86 | 14.86 |  |
| 9,0 | 0.3 | 3% | log<sub>10</sub>BF | 138.27 | 67.50 | 5.60 | 38.98 | 88.00 | 129.53 | 176.81 | 269.42 | 300.00 | 304.92 |  |
|  |  |  | -log<sub>10</sub>PPNA | 142.66 | 67.19 | 10.11 | 43.50 | 92.52 | 134.04 | 181.33 | 273.93 | 300.00 | 306.17 |  |
|  |  |  | -log<sub>10</sub>ASTpv | 15.23 | 5.37 | 3.73 | 8.09 | 11.41 | 14.08 | 18.47 | 25.55 | 30.46 | 34.76 |  |
|  | 0.1 | 13% | log<sub>10</sub>BF | 63.42 | 64.23 | -5.03 | -3.23 | 13.52 | 48.72 | 96.29 | 201.05 | 284.78 | 303.37 |  |
|  |  |  | -log<sub>10</sub>PPNA | 67.91 | 64.09 | 0.11 | 1.31 | 18.04 | 53.23 | 100.80 | 205.57 | 289.25 | 307.89 |  |
|  |  |  | -log<sub>10</sub>ASTpv | 5.36 | 3.19 | 0.07 | 1.31 | 3.09 | 4.81 | 7.03 | 11.85 | 15.46 | 17.46 |  |
| 5,4 | 0.3 | 12.4% | log<sub>10</sub>BF | 215.86 | 60.21 | 34.66 | 111.91 | 172.70 | 217.38 | 266.32 | 300.00 | 301.70 | 307.35 |  |
|  |  |  | -log<sub>10</sub>PPNA | 219.77 | 59.38 | 39.17 | 116.43 | 177.22 | 221.90 | 270.83 | 300.00 | 303.80 | 307.01 |  |
|  |  |  | -log<sub>10</sub>ASTpv | 23.88 | 5.96 | 8.05 | 14.39 | 19.81 | 23.71 | 28.19 | 33.76 | 38.67 | 40.10 |  |
|  | 0.1 | 26.2% | log<sub>10</sub>BF | 162.58 | 75.70 | -3.20 | 34.04 | 106.51 | 161.99 | 219.54 | 298.60 | 300.00 | 305.56 |  |
|  |  |  | -log<sub>10</sub>PPNA | 166.90 | 75.34 | 1.34 | 38.56 | 111.02 | 166.50 | 224.06 | 300.00 | 302.77 | 306.51 |  |
|  |  |  | -log<sub>10</sub>ASTpv | 8.18 | 2.85 | 1.57 | 3.71 | 6.25 | 8.13 | 10.16 | 12.84 | 15.23 | 16.42 |  |

Legend: $m$ - allele frequency at the risk SNP; $m = 0.3, 0.1$ . The number of positively and negatively associated traits are denoted by $K_1^+$ and $K_1^-$ , respectively. Hence the total number of associated traits is $K_1 = K_1^+ + K_1^-$ . The following abbreviations denote - log<sub>10</sub>BF: log<sub>10</sub>(Bayes factor), -log<sub>10</sub>PPNA: -log<sub>10</sub>(posterior probability of null association), -log<sub>10</sub>(ASTpv): -log<sub>10</sub>(ASSET p-value). For multiple studies with overlapping subjects, the combined strategy of CPBayes is implemented. Of note, uncor% denotes the percentage of replications in which the combined strategy of CPBayes used the uncorrelated version of CPBayes. Different summary measures obtained across 500 replications are provided: mean, standard deviation (sd), minimum (min), 5% quantile, 25% quantile, 50% quantile, 75% quantile, 95% quantile, 99% quantile, and maximum (max).

**Table S13:** Summary of measures for the evidence of the overall pleiotropic association for a **cohort** study with **5** binary phenotypes. Here 1, 2, and 3 among 5 traits are associated.

| $K_1^+, K_1^-$ | $m$ | uncor% | | mean | sd | min | quantile | | | | | | | max |
| --- | --- | --- | --- | --- | --- | --- | --- | --- | --- | --- | --- | --- | --- | --- |
|  |  |  |  |  |  |  | 5% | 25% | 50% | 75% | 95% | 99% |  |  |
| 1,0 | 0.3 | 0.4% | log<sub>10</sub>BF | 2.35 | 5.71 | -2.60 | -1.95 | -0.68 | 0.30 | 3.18 | 13.14 | 25.07 | 36.20 |  |
|  |  |  | -log<sub>10</sub>PPNA | 3.94 | 5.64 | 0.03 | 0.13 | 0.87 | 1.80 | 4.67 | 14.63 | 26.56 | 37.69 |  |
|  |  |  | -log<sub>10</sub>ASTpv | 3.28 | 1.80 | 0.14 | 0.90 | 1.94 | 3.03 | 4.41 | 6.41 | 8.54 | 9.60 |  |
|  | 0.1 | 0.8% | log<sub>10</sub>BF | 5.58 | 13.62 | -2.56 | -1.84 | -0.60 | 0.31 | 5.62 | 31.54 | 61.89 | 134.17 |  |
|  |  |  | -log<sub>10</sub>PPNA | 7.15 | 13.58 | 0.04 | 0.16 | 0.95 | 1.81 | 7.11 | 33.03 | 63.38 | 135.67 |  |
|  |  |  | -log<sub>10</sub>ASTpv | 2.94 | 1.59 | 0.22 | 0.79 | 1.82 | 2.68 | 3.86 | 5.97 | 7.59 | 9.02 |  |
| 2,0 | 0.3 | 0.4% | log<sub>10</sub>BF | 16.93 | 20.10 | -2.49 | -1.30 | 0.49 | 8.88 | 29.37 | 56.92 | 70.49 | 144.78 |  |
|  |  |  | -log<sub>10</sub>PPNA | 18.46 | 20.06 | 0.04 | 0.40 | 1.99 | 10.37 | 30.86 | 58.41 | 71.98 | 146.27 |  |
|  |  |  | -log<sub>10</sub>ASTpv | 5.92 | 2.65 | 0.42 | 2.15 | 4.05 | 5.74 | 7.39 | 10.80 | 12.73 | 15.19 |  |
|  | 0.1 | 0.6% | log<sub>10</sub>BF | 37.24 | 44.95 | -2.39 | -0.98 | 0.70 | 20.95 | 57.70 | 134.10 | 172.37 | 260.25 |  |
|  |  |  | -log<sub>10</sub>PPNA | 38.76 | 44.93 | 0.05 | 0.63 | 2.19 | 22.44 | 59.19 | 135.59 | 173.86 | 261.74 |  |
|  |  |  | -log<sub>10</sub>ASTpv | 5.24 | 2.25 | 0.18 | 2.04 | 3.57 | 5.12 | 6.59 | 9.31 | 11.24 | 13.47 |  |
| 1,1 | 0.3 | 1% | log<sub>10</sub>BF | 28.69 | 25.32 | -2.30 | -0.05 | 8.70 | 23.34 | 43.06 | 75.65 | 104.41 | 131.11 |  |
|  |  |  | -log<sub>10</sub>PPNA | 30.19 | 25.32 | 0.06 | 1.46 | 10.19 | 24.83 | 44.55 | 77.15 | 105.90 | 132.60 |  |
|  |  |  | -log<sub>10</sub>ASTpv | 6.87 | 2.63 | 0.46 | 3.05 | 5.04 | 6.77 | 8.49 | 11.44 | 13.71 | 17.00 |  |
|  | 0.1 | 1.4% | log<sub>10</sub>BF | 75.35 | 57.50 | -1.77 | 0.88 | 29.54 | 66.54 | 110.21 | 193.99 | 242.63 | 270.47 |  |
|  |  |  | -log<sub>10</sub>PPNA | 76.84 | 57.50 | 0.18 | 2.37 | 31.04 | 68.03 | 111.70 | 195.49 | 244.12 | 271.96 |  |
|  |  |  | -log<sub>10</sub>ASTpv | 6.82 | 2.54 | 1.20 | 3.03 | 5.02 | 6.59 | 8.28 | 11.60 | 13.07 | 15.80 |  |
| 3,0 | 0.3 | 0.6% | log<sub>10</sub>BF | 29.04 | 32.40 | -2.15 | -1.04 | 1.24 | 17.90 | 47.42 | 96.83 | 130.47 | 157.60 |  |
|  |  |  | -log<sub>10</sub>PPNA | 30.56 | 32.38 | 0.09 | 0.58 | 2.73 | 19.39 | 48.91 | 98.32 | 131.96 | 159.09 |  |
|  |  |  | -log<sub>10</sub>ASTpv | 8.24 | 3.29 | 1.32 | 3.59 | 5.76 | 7.84 | 10.32 | 14.16 | 16.41 | 19.55 |  |
|  | 0.1 | 0.6% | log<sub>10</sub>BF | 66.73 | 70.53 | -2.29 | -0.76 | 5.17 | 43.91 | 110.55 | 204.88 | 285.55 | 305.56 |  |
|  |  |  | -log<sub>10</sub>PPNA | 68.23 | 70.48 | 0.06 | 0.80 | 6.66 | 45.40 | 112.04 | 206.37 | 287.04 | 307.05 |  |
|  |  |  | -log<sub>10</sub>ASTpv | 7.44 | 2.92 | 0.89 | 3.37 | 5.22 | 7.14 | 9.15 | 13.09 | 15.32 | 18.67 |  |
| 2,1 | 0.3 | 0.4% | log<sub>10</sub>BF | 60.72 | 36.96 | -0.05 | 8.79 | 31.84 | 58.14 | 83.58 | 122.86 | 149.41 | 225.07 |  |
|  |  |  | -log<sub>10</sub>PPNA | 62.21 | 36.96 | 1.46 | 10.28 | 33.33 | 59.63 | 85.08 | 124.35 | 150.91 | 226.56 |  |
|  |  |  | -log<sub>10</sub>ASTpv | 9.64 | 2.96 | 3.58 | 5.21 | 7.51 | 9.36 | 11.55 | 14.86 | 16.86 | 20.40 |  |
|  | 0.1 | 0.2% | log<sub>10</sub>BF | 144.11 | 78.84 | 0.47 | 23.29 | 84.36 | 139.72 | 203.80 | 300.00 | 300.00 | 305.28 |  |
|  |  |  | -log<sub>10</sub>PPNA | 145.54 | 78.71 | 1.96 | 24.78 | 85.85 | 141.21 | 205.29 | 300.00 | 300.00 | 306.77 |  |
|  |  |  | -log<sub>10</sub>ASTpv | 9.37 | 2.69 | 3.52 | 5.27 | 7.42 | 9.27 | 11.18 | 13.93 | 16.43 | 20.06 |  |

Legend: $m$ - allele frequency at the risk SNP; $m = 0.3, 0.1$ . The number of positively and negatively associated traits are denoted by $K_1^+$ and $K_1^-$ , respectively. Hence the total number of associated traits is $K_1 = K_1^+ + K_1^-$ . The following abbreviations denote - log<sub>10</sub>BF: log<sub>10</sub>(Bayes factor), -log<sub>10</sub>PPNA: -log<sub>10</sub>(posterior probability of null association), -log<sub>10</sub>(ASTpv): -log<sub>10</sub>(ASSET p-value). For a cohort study, since the summary statistics are correlated, the combined strategy of CPBayes is implemented. Of note, uncor% denotes the percentage of replications in which the combined strategy of CPBayes used the uncorrelated version of CPBayes. Different summary measures obtained across 500 replications are provided: mean, standard deviation (sd), minimum (min), 5% quantile, 25% quantile, 50% quantile, 75% quantile, 95% quantile, 99% quantile, and maximum (max).

**Table S14:** Summary of measures for the evidence of the overall pleiotropic association for a **cohort** study with **10** binary phenotypes. Here 2, 4, and 6 among 10 traits are associated.

| $K_1^+, K_1^-$ | $m$ | uncor% | | mean | sd | min | quantile | | | | | | |
| --- | --- | --- | --- | --- | --- | --- | --- | --- | --- | --- | --- | --- | --- |
|  |  |  |  |  |  |  | 5% | 25% | 50% | 75% | 95% | 99% | max |
| 2,0 | 0.3 | 1.4% | log<sub>10</sub>BF | 15.43 | 22.86 | -3.84 | -2.83 | -0.96 | 6.94 | 23.85 | 64.26 | 94.62 | 141.25 |
|  |  |  | -log<sub>10</sub>PPNA | 18.47 | 22.83 | 0.06 | 0.40 | 2.05 | 9.95 | 26.86 | 67.27 | 97.63 | 144.26 |
|  |  |  | -log<sub>10</sub>ASTpv | 4.42 | 2.22 | 0.21 | 1.32 | 2.82 | 4.09 | 5.69 | 8.65 | 9.98 | 14.28 |
|  | 0.1 | 2% | log<sub>10</sub>BF | 39.56 | 52.79 | -3.65 | -2.65 | -0.73 | 17.77 | 61.14 | 150.17 | 208.91 | 300.00 |
|  |  |  | -log<sub>10</sub>PPNA | 42.59 | 52.74 | 0.09 | 0.52 | 2.28 | 20.78 | 64.15 | 153.18 | 211.92 | 300.00 |
|  |  |  | -log<sub>10</sub>ASTpv | 3.83 | 1.94 | 0.32 | 0.90 | 2.47 | 3.54 | 5.04 | 7.19 | 9.34 | 10.46 |
| 1,1 | 0.3 | 2.4% | log<sub>10</sub>BF | 25.82 | 24.81 | -3.56 | -1.85 | 5.20 | 19.64 | 39.79 | 74.28 | 98.89 | 120.00 |
|  |  |  | -log<sub>10</sub>PPNA | 28.84 | 24.80 | 0.11 | 1.18 | 8.21 | 22.65 | 42.80 | 77.29 | 101.90 | 123.01 |
|  |  |  | -log<sub>10</sub>ASTpv | 4.88 | 2.42 | 0.18 | 1.58 | 3.06 | 4.63 | 6.24 | 9.40 | 11.67 | 15.97 |
|  | 0.1 | 2.2% | log<sub>10</sub>BF | 53.73 | 49.08 | -3.88 | -0.92 | 17.83 | 40.63 | 76.23 | 145.13 | 224.11 | 306.52 |
|  |  |  | -log<sub>10</sub>PPNA | 56.72 | 48.95 | 0.06 | 2.10 | 20.84 | 43.64 | 79.24 | 148.14 | 227.12 | 305.48 |
|  |  |  | -log<sub>10</sub>ASTpv | 4.93 | 2.41 | 0.06 | 1.74 | 3.25 | 4.56 | 6.36 | 9.42 | 11.40 | 17.14 |
| 4,0 | 0.3 | 2.2% | log<sub>10</sub>BF | 56.33 | 54.33 | -3.51 | -1.89 | 6.75 | 46.66 | 86.96 | 169.80 | 215.64 | 244.32 |
|  |  |  | -log<sub>10</sub>PPNA | 59.35 | 54.32 | 0.12 | 1.15 | 9.76 | 49.67 | 89.97 | 172.81 | 218.65 | 247.33 |
|  |  |  | -log<sub>10</sub>ASTpv | 8.47 | 3.36 | 0.93 | 3.89 | 5.94 | 7.89 | 10.38 | 14.76 | 18.33 | 20.98 |
|  | 0.1 | 3.8% | log<sub>10</sub>BF | 119.80 | 100.88 | -3.34 | -1.95 | 23.09 | 104.41 | 193.24 | 300.00 | 304.52 | 308.12 |
|  |  |  | -log<sub>10</sub>PPNA | 122.52 | 100.33 | 0.17 | 1.09 | 26.10 | 107.42 | 196.25 | 300.00 | 300.01 | 308.03 |
|  |  |  | -log<sub>10</sub>ASTpv | 7.23 | 2.92 | 1.29 | 3.07 | 5.24 | 6.92 | 8.79 | 12.70 | 15.48 | 17.78 |
| 2,2 | 0.3 | 3.8% | log<sub>10</sub>BF | 99.93 | 46.20 | -2.36 | 26.66 | 66.67 | 99.40 | 130.90 | 176.83 | 216.07 | 254.43 |
|  |  |  | -log<sub>10</sub>PPNA | 102.94 | 46.20 | 0.74 | 29.67 | 69.68 | 102.41 | 133.91 | 179.84 | 219.08 | 257.44 |
|  |  |  | -log<sub>10</sub>ASTpv | 10.42 | 2.98 | 3.39 | 5.87 | 8.26 | 10.36 | 12.27 | 15.42 | 18.26 | 21.29 |
|  | 0.1 | 4.4% | log<sub>10</sub>BF | 227.05 | 78.66 | -2.35 | 71.00 | 171.94 | 249.85 | 300.00 | 300.00 | 303.33 | 307.17 |
|  |  |  | -log<sub>10</sub>PPNA | 229.08 | 77.75 | 0.75 | 74.01 | 174.95 | 252.86 | 300.00 | 300.00 | 302.47 | 306.32 |
|  |  |  | -log<sub>10</sub>ASTpv | 10.63 | 3.30 | 1.40 | 5.53 | 8.25 | 10.43 | 12.64 | 16.28 | 18.50 | 26.30 |
| 6,0 | 0.3 | 5.6% | log<sub>10</sub>BF | 88.10 | 77.83 | -3.57 | -1.81 | 17.54 | 73.16 | 144.29 | 232.21 | 300.00 | 302.61 |
|  |  |  | -log<sub>10</sub>PPNA | 91.09 | 77.73 | 0.11 | 1.23 | 20.55 | 76.17 | 147.30 | 235.22 | 300.00 | 305.62 |
|  |  |  | -log<sub>10</sub>ASTpv | 11.59 | 4.12 | 1.83 | 5.67 | 8.64 | 11.23 | 13.91 | 19.29 | 22.88 | 29.19 |
|  | 0.1 | 4.8% | log<sub>10</sub>BF | 174.39 | 115.88 | -3.43 | -1.72 | 66.20 | 181.02 | 300.00 | 300.00 | 301.73 | 306.12 |
|  |  |  | -log<sub>10</sub>PPNA | 176.49 | 114.88 | 0.14 | 1.31 | 69.20 | 184.03 | 300.00 | 300.00 | 303.51 | 307.87 |
|  |  |  | -log<sub>10</sub>ASTpv | 10.22 | 3.52 | 1.83 | 5.29 | 7.76 | 9.89 | 12.26 | 16.74 | 20.79 | 23.54 |
| 3,3 | 0.3 | 5.2% | log<sub>10</sub>BF | 202.10 | 57.94 | 32.55 | 100.05 | 165.63 | 204.42 | 245.35 | 300.00 | 302.67 | 305.65 |
|  |  |  | -log<sub>10</sub>PPNA | 204.94 | 57.66 | 35.56 | 103.06 | 168.64 | 207.43 | 248.36 | 300.00 | 302.39 | 307.74 |
|  |  |  | -log<sub>10</sub>ASTpv | 15.67 | 3.58 | 6.08 | 10.05 | 13.19 | 15.49 | 17.99 | 21.41 | 25.39 | 27.98 |
|  | 0.1 | 4% | log<sub>10</sub>BF | 295.46 | 22.55 | 93.92 | 277.38 | 300.00 | 300.00 | 300.00 | 300.00 | 300.00 | 305.60 |
|  |  |  | -log<sub>10</sub>PPNA | 295.64 | 21.98 | 96.93 | 280.39 | 300.00 | 300.00 | 300.00 | 300.00 | 300.00 | 304.24 |
|  |  |  | -log<sub>10</sub>ASTpv | 15.89 | 3.57 | 7.36 | 10.46 | 13.71 | 15.67 | 18.22 | 21.72 | 24.68 | 32.26 |

Legend: $m$ - allele frequency at the risk SNP; $m = 0.3, 0.1$ . The number of positively and negatively associated traits are denoted by $K_1^+$ and $K_1^-$ , respectively. Hence the total number of associated traits is $K_1 = K_1^+ + K_1^-$ . The following abbreviations denote - log<sub>10</sub>BF: log<sub>10</sub>(Bayes factor), -log<sub>10</sub>PPNA: -log<sub>10</sub>(posterior probability of null association), -log<sub>10</sub>(ASTpv): -log<sub>10</sub>(ASSET p-value). For a cohort study, since the summary statistics are correlated, the combined strategy of CPBayes is implemented. Of note, uncor% denotes the percentage of replications in which the combined strategy of CPBayes used the uncorrelated version of CPBayes. Different summary measures obtained across 500 replications are provided: mean, standard deviation (sd), minimum (min), 5% quantile, 25% quantile, 50% quantile, 75% quantile, 95% quantile, 99% quantile, and maximum (max).

**Table S15:**
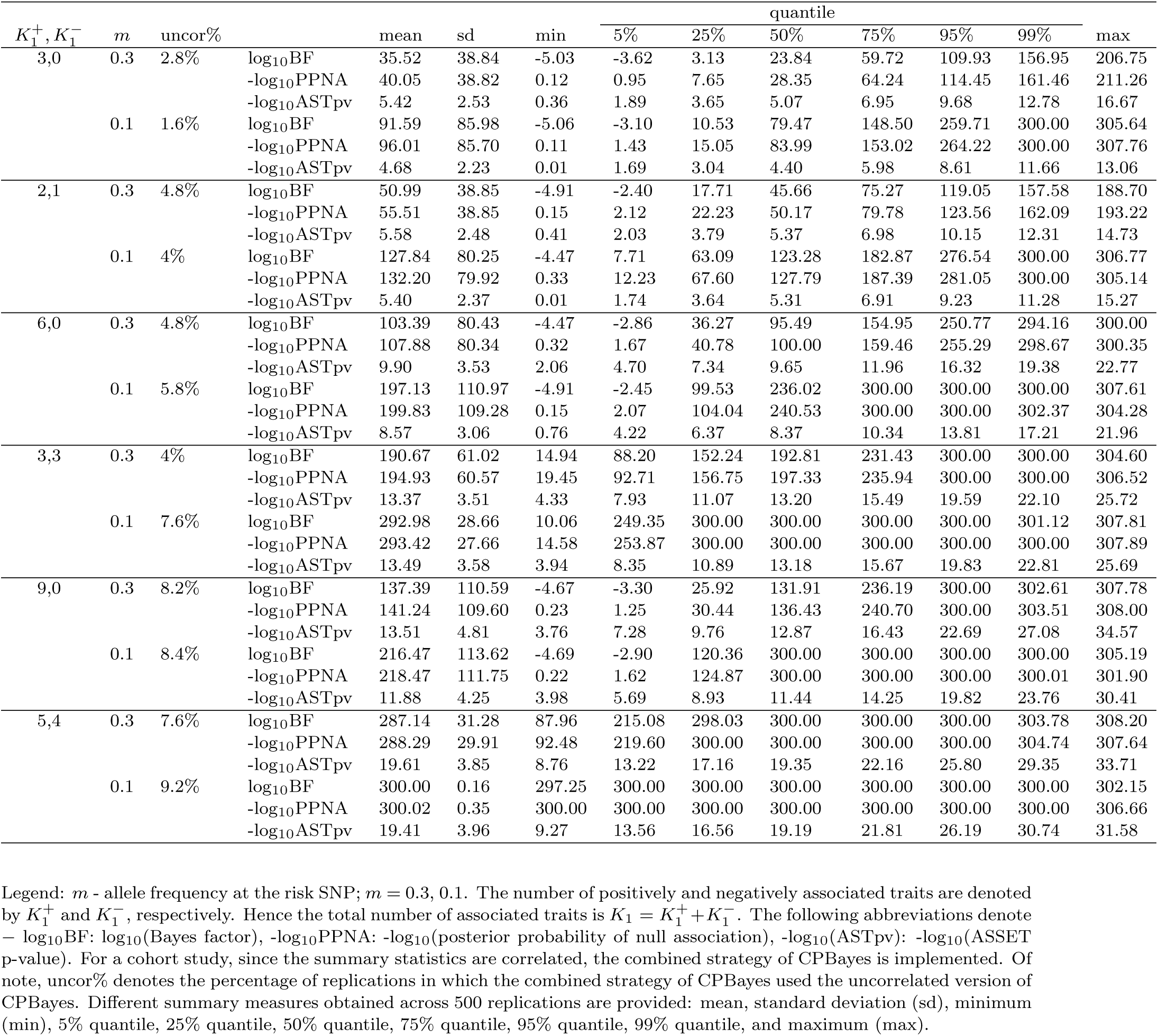
Summary of measures for the evidence of the overall pleiotropic association for a **cohort** study with **15** binary phenotypes. Here 3, 6, and 9 among 15 traits are associated.

## 9 Comparison of selection accuracy in cohort study

In the next diagram, we present the results of selection accuracy obtained by various methods for cohort study.

**Figure S13:**
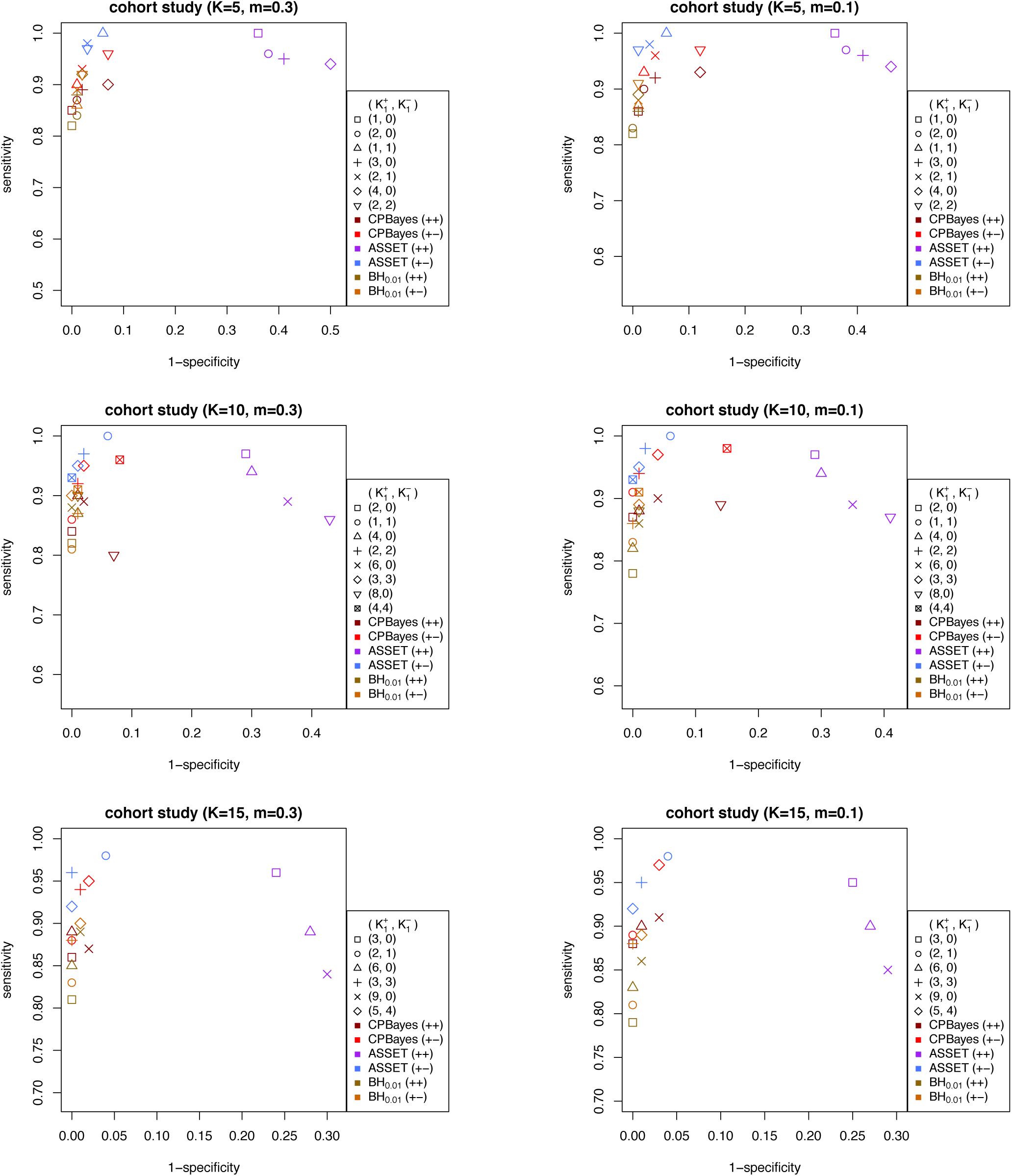
Comparison of the accuracy of selection of associated traits by CPBayes and ASSET for cohort study. The total number of phenotypes/studies is denoted by *K* and *m* denotes the minor allele frequency at the risk SNP.

## 10 Simulation results for 50 traits

**Table S16:**
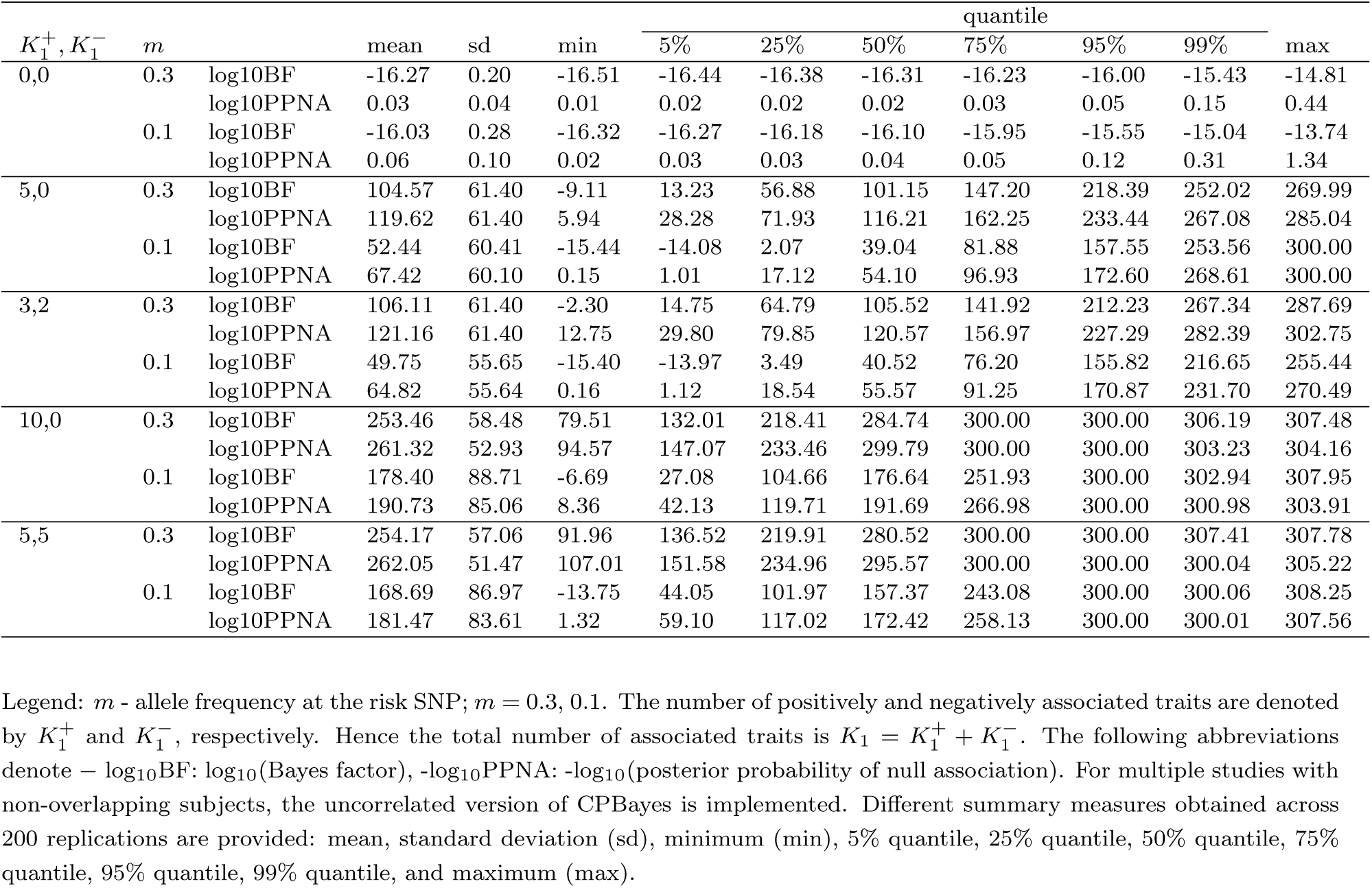
Simulation study for **50** traits. Summary of measures for the evidence of the overall pleiotropic association for **50** non-overlapping case-control studies. Here 0, 5, and 10 among 50 traits are associated.

**Table S17:** Simulation study for **50** traits. Summary of measures for the evidence of the overall pleiotropic association for **50** overlapping case-control studies. Here 0, 5, and 10 among 50 traits are associated.

| $K_1^+, K_1^-$ | $m$ | | mean | sd | min | quantile | | | | | | |
| --- | --- | --- | --- | --- | --- | --- | --- | --- | --- | --- | --- | --- |
|  |  |  |  |  |  | 5% | 25% | 50% | 75% | 95% | 99% | max |
| 0,0 | 0.3 | log<sub>10</sub>BF | -16.43 | 0.11 | -16.61 | -16.56 | -16.50 | -16.44 | -16.38 | -16.25 | -16.08 | -15.83 |
|  |  | -log<sub>10</sub>PPNA | 0.02 | 0.006 | 0.01 | 0.01 | 0.02 | 0.02 | 0.02 | 0.03 | 0.04 | 0.07 |
|  | 0.1 | log<sub>10</sub>BF | -16.18 | 0.25 | -16.42 | -16.38 | -16.30 | -16.24 | -16.15 | -15.83 | -15.51 | -13.96 |
|  |  | -log<sub>10</sub>PPNA | 0.04 | 0.08 | 0.02 | 0.02 | 0.02 | 0.03 | 0.03 | 0.07 | 0.13 | 1.12 |
| 5,0 | 0.3 | log<sub>10</sub>BF | 80.75 | 49.38 | -10.03 | 9.21 | 45.43 | 74.39 | 107.94 | 170.54 | 216.63 | 240.42 |
|  |  | -log<sub>10</sub>PPNA | 95.80 | 49.38 | 5.02 | 24.26 | 60.48 | 89.44 | 123.00 | 185.60 | 231.68 | 255.47 |
|  | 0.1 | log<sub>10</sub>BF | 44.74 | 48.08 | -15.93 | -14.86 | 1.51 | 38.10 | 74.46 | 124.66 | 164.88 | 205.38 |
|  |  | -log<sub>10</sub>PPNA | 59.82 | 48.05 | 0.05 | 0.41 | 16.56 | 53.15 | 89.51 | 139.71 | 179.94 | 220.43 |
| 3,2 | 0.3 | log<sub>10</sub>BF | 90.92 | 52.29 | -8.18 | 19.74 | 51.37 | 85.36 | 120.45 | 192.00 | 224.04 | 235.20 |
|  |  | -log<sub>10</sub>PPNA | 105.97 | 52.29 | 6.87 | 34.80 | 66.42 | 100.41 | 135.50 | 207.05 | 239.10 | 250.25 |
|  | 0.1 | log<sub>10</sub>BF | 49.80 | 55.62 | -15.27 | -14.33 | 3.45 | 34.91 | 81.20 | 161.52 | 196.32 | 221.93 |
|  |  | -log<sub>10</sub>PPNA | 64.86 | 55.60 | 0.21 | 0.80 | 18.51 | 49.96 | 96.25 | 176.58 | 211.37 | 236.98 |
| 10,0 | 0.3 | log<sub>10</sub>BF | 199.00 | 67.19 | 25.03 | 96.56 | 146.74 | 201.41 | 249.66 | 300.00 | 300.01 | 307.05 |
|  |  | -log<sub>10</sub>PPNA | 212.35 | 64.74 | 40.08 | 111.61 | 161.79 | 216.46 | 264.71 | 300.00 | 305.53 | 306.50 |
|  | 0.1 | log<sub>10</sub>BF | 130.40 | 89.63 | -15.33 | -12.20 | 58.70 | 128.76 | 189.76 | 285.52 | 300.02 | 306.13 |
|  |  | -log<sub>10</sub>PPNA | 144.74 | 88.29 | 0.18 | 2.85 | 73.75 | 143.81 | 204.81 | 300.00 | 300.16 | 305.42 |
| 5,5 | 0.3 | log<sub>10</sub>BF | 226.35 | 61.43 | 86.31 | 122.24 | 179.19 | 232.38 | 288.33 | 300.00 | 303.86 | 307.56 |
|  |  | -log<sub>10</sub>PPNA | 237.90 | 57.39 | 101.36 | 137.29 | 194.24 | 247.43 | 300.00 | 300.00 | 305.03 | 306.94 |
|  | 0.1 | log<sub>10</sub>BF | 159.54 | 83.40 | -14.64 | 21.29 | 98.02 | 158.45 | 222.71 | 300.00 | 301.30 | 308.25 |
|  |  | -log<sub>10</sub>PPNA | 173.33 | 81.33 | 0.56 | 36.34 | 113.07 | 173.50 | 237.76 | 300.00 | 302.45 | 305.23 |

Legend: $m$ - allele frequency at the risk SNP; $m = 0.3, 0.1$ . The number of positively and negatively associated traits are denoted by $K_1^+$ and $K_1^-$ , respectively. Hence the total number of associated traits is $K_1 = K_1^+ + K_1^-$ . The following abbreviations denote $-\log_{10}$ BF: $\log_{10}$ (Bayes factor), $-\log_{10}$ PPNA: $-\log_{10}$ (posterior probability of null association). For multiple studies with overlapping subjects, the combined strategy of CPBayes is implemented. Different summary measures obtained across 200 replications are provided: mean, standard deviation (sd), minimum (min), 5% quantile, 25% quantile, 50% quantile, 75% quantile, 95% quantile, 99% quantile, and maximum (max).

**Table S18:** Simulation study for 50 traits. Accuracy in selection of associated traits by CPBayes and BH_0.01_ for 50 nonoverlapping case-control studies. The number of positively and negatively associated traits are denoted by 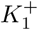 and 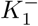, respectively. And *m* denotes the minor allele frequency at the risk SNP.

| $K_1^+, K_1^-$ | $m$ | CPBayes | | BH<sub>0.01</sub> | |
| --- | --- | --- | --- | --- | --- |
|  |  | specificity | sensitivity | specificity | sensitivity |
| 5,0 | 0.3 | 1.00 | 0.79 | 0.998 | 0.84 |
|  | 0.1 | 1.00 | 0.58 | 1.00 | 0.61 |
| 3,2 | 0.3 | 1.00 | 0.81 | 1.00 | 0.85 |
|  | 0.1 | 1.00 | 0.57 | 1.00 | 0.61 |
| 10,0 | 0.3 | 1.00 | 0.82 | 1.00 | 0.86 |
|  | 0.1 | 1.00 | 0.62 | 1.00 | 0.65 |
| 5,5 | 0.3 | 1.00 | 0.82 | 1.00 | 0.86 |
|  | 0.1 | 1.00 | 0.59 | 0.998 | 0.61 |

**Table S19:**
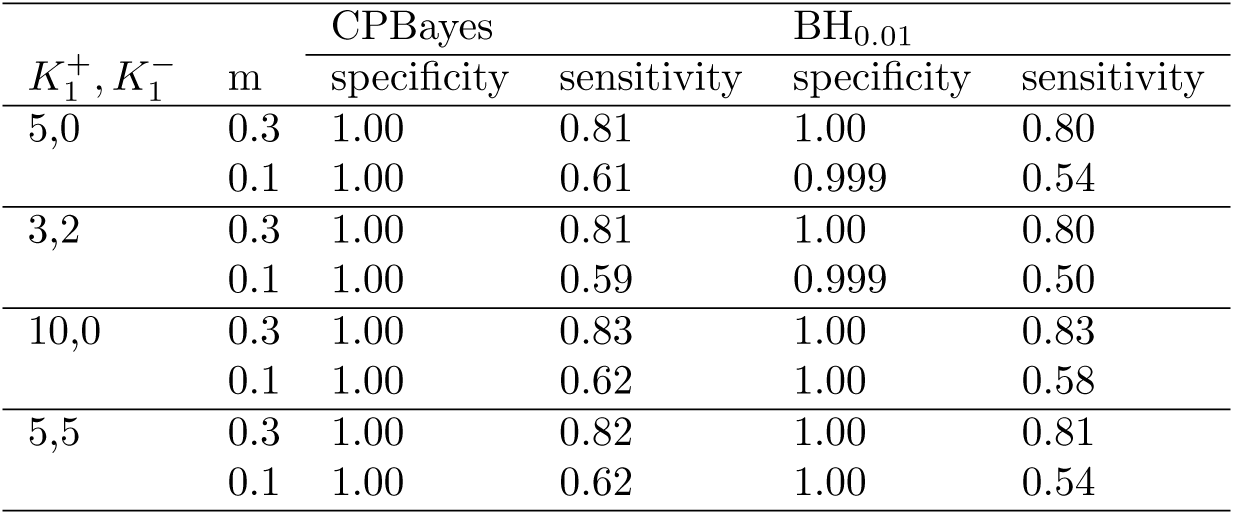
Simulation study for 50 traits. Accuracy in selection of associated traits by CPBayes and BH_0.01_ for 50 overlapping case-control studies. The number of positively and negatively associated traits are denoted by 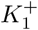 and 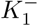, respectively. And *m* denotes the minor allele frequency at the risk SNP.

| $K_1^+, K_1^-$ | $m$ | CPBayes | | BH<sub>0.01</sub> | |
| --- | --- | --- | --- | --- | --- |
|  |  | specificity | sensitivity | specificity | sensitivity |
| 5,0 | 0.3 | 1.00 | 0.81 | 1.00 | 0.80 |
|  | 0.1 | 1.00 | 0.61 | 0.999 | 0.54 |
| 3,2 | 0.3 | 1.00 | 0.81 | 1.00 | 0.80 |
|  | 0.1 | 1.00 | 0.59 | 0.999 | 0.50 |
| 10,0 | 0.3 | 1.00 | 0.83 | 1.00 | 0.83 |
|  | 0.1 | 1.00 | 0.62 | 1.00 | 0.58 |
| 5,5 | 0.3 | 1.00 | 0.82 | 1.00 | 0.81 |
|  | 0.1 | 1.00 | 0.62 | 1.00 | 0.54 |

## 11 Comparison between the continuous spike and Dirac spike

We carried out simulation study to compare the selection accuracy of two different type of spikes. We chose the same set-up of multiple overlapping case-control studies considered while comparing CPBayes and ASSET in the main simulation study. We implemented the Gibbs sampling algorithm for the Dirac spike described in Algorithm S1. Since the summary statistics are correlated, we implemented the combined strategy for the Dirac spike as well as for the continuous spike. The slab variance for the Dirac spike (*b*^2^) is considered to vary in 0.8 − 1.2 (the same as that for the continuous spike). We compute the mean specificity and sensitivity across 200 replications. The results are provided in Figure S14 (see the next page).

We observe that the Dirac spike produces less specificity than the continuous spike. The Dirac spike suffers from reduced specificity more for larger number of associated traits (*K*_1_). The continuous spike consistently yields very good level of specificity across different scenarios. The Dirac spike offers higher sensitivity, but at the expense of lower specificity compared to the continuous spike. For example, for *K* = 10, 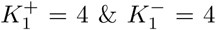 (*K*_1_ = 8), and *m* = 0.1, the Dirac spike produced a mean specificity of 64% and sensitivity of 87%, whereas the continuous spike gave a mean specificity of 97% and sensitivity of 72%. Similarly, for *K* = 15, 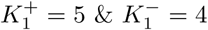 (*K*_1_ = 9), and *m* = 0.1, the Dirac spike gave 65% specificity and 85% sensitivity, whereas the continuous spike produced 99% specificity and 69% sensitivity.

**Figure S14:**
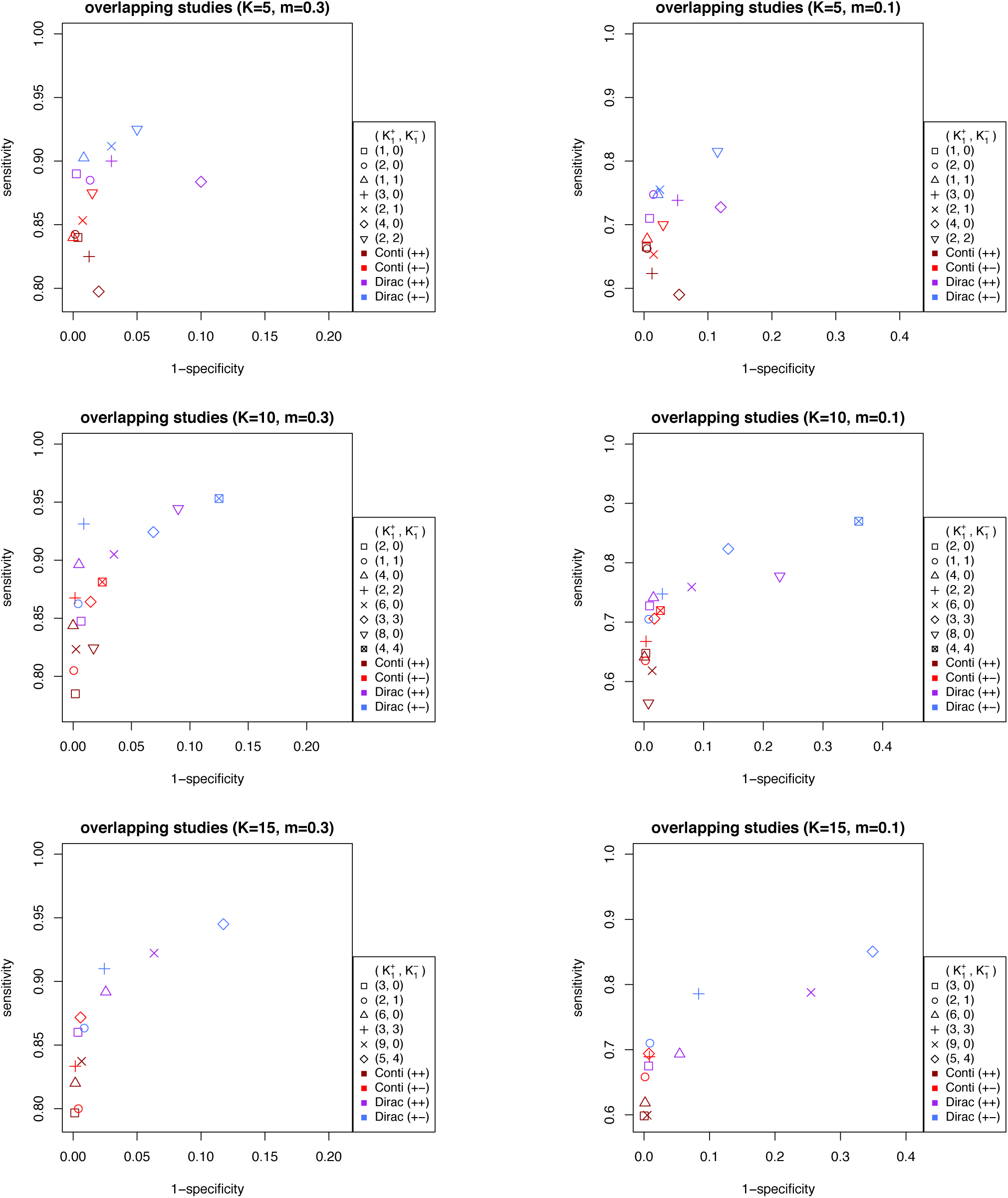
Comparison of the accuracy of selection of associated traits by the continuous and the Dirac spike for multiple overlapping case-control studies. The total number of phenotypes/studies is denoted by *K* and *m* denotes the minor allele frequency at the risk SNP.

## 12 List 22 traits in the GERA cohort analyzed by CPBayes and ASSET

**Table S20:** Name of 22 phenotypes in the GERA cohort analyzed by CPBayes and ASSET

| GERA cohort phenotypes |  |  |  |
| --- | --- | --- | --- |
| Asthma | Type 2 Diabetes | Iron Deficiency | Peptic Ulcer |
| Allergic Rhinitis | Dyslipidemia | Irritable Bowel Syndrome | Psychiatric disorders |
| Cardiovascular Disease | Hypertension | Macular Degeneration | Stress Disorders |
| Cancers | Hemorrhoids | Osteoarthritis | Varicose Veins |
| Depressive Disorder | Abdominal Hernia | Osteoporosis |  |
| Dermatophytosis | Insomnia | Peripheral Vascular Disease |  |

## 13 Manhattan plots for CPBayes and ASSET

Here we present the Manhattan plots for the two methods. The red line corresponds to the genome-wide cutoff for log_10_BF of CPBayes and –log_10_(p-value) of ASSET. Note that, log_10_BF of CPBayes can be negative, in particular, for null SNPs. The Manhattan plots clearly show that CPBayes detected substantially larger number of pleiotropic regions than ASSET.

**Figure S15:**
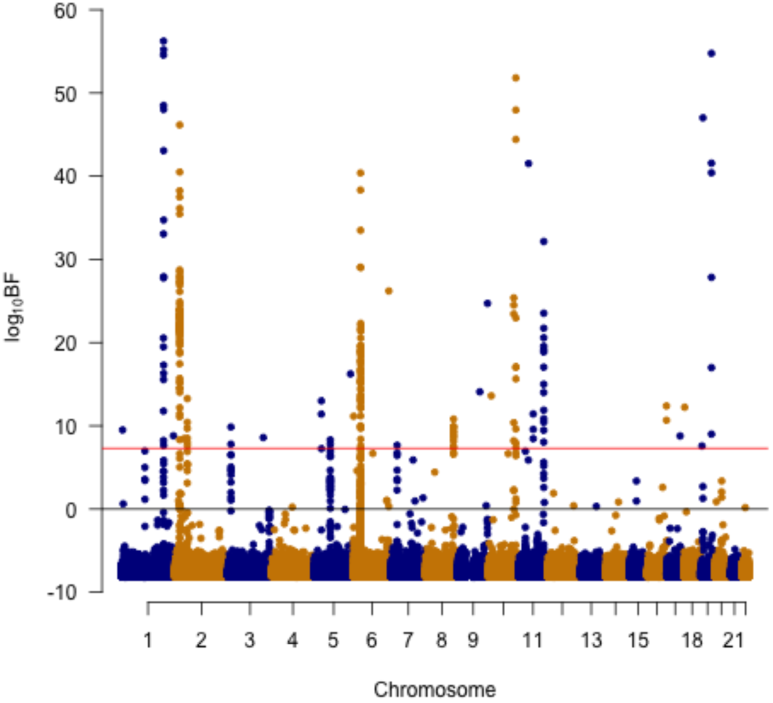
Manhattan plot for CPBayes.

**Figure S16:**
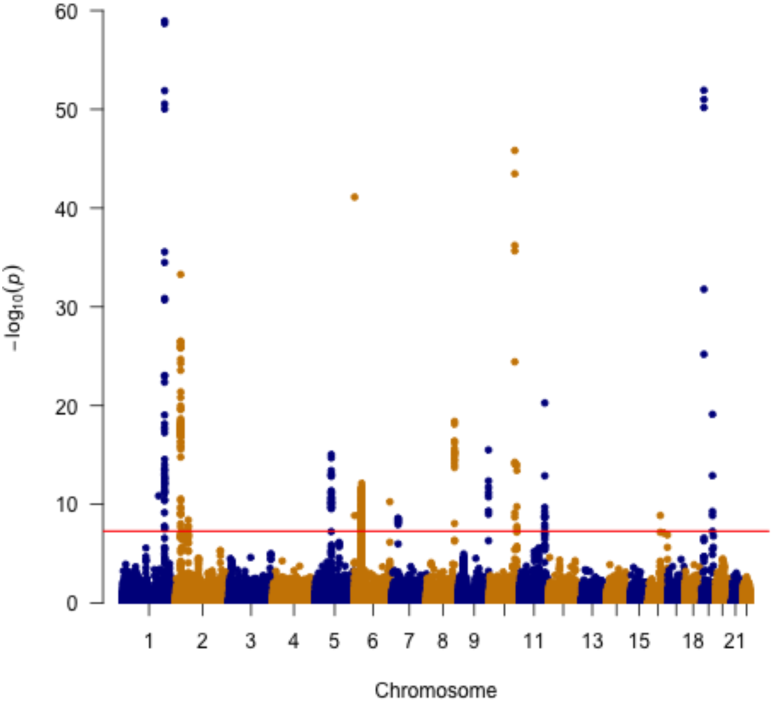
Manhattan plot for ASSET.

## 14 Forest plots for independent pleiotropic SNPs detected by CP-Bayes at which it selected at lease two traits in GERA cohort

**Figure S17:**
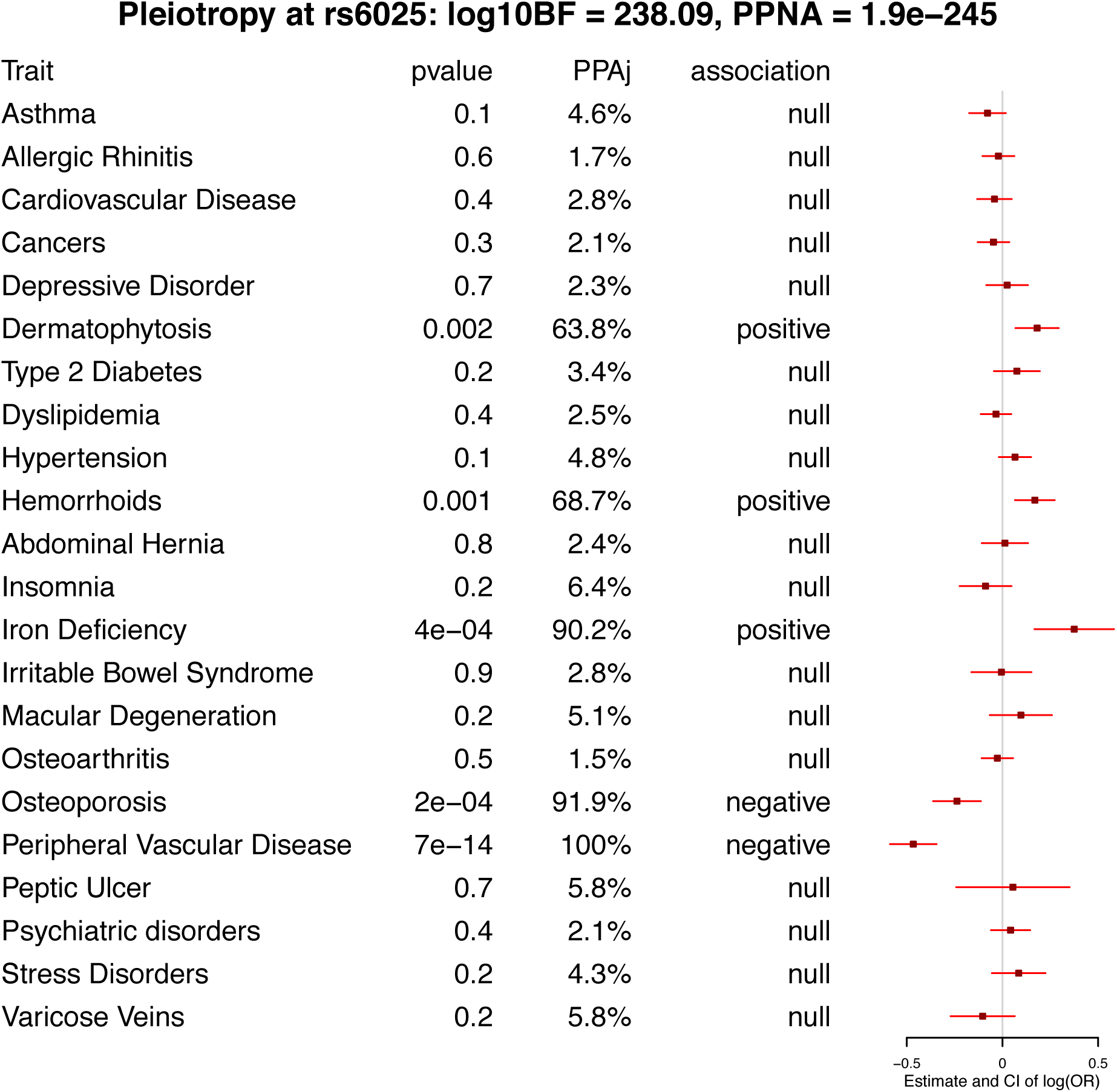
Forest plot for pleiotropic signal at rs6025 detected by CPBayes.

**Figure S18:**
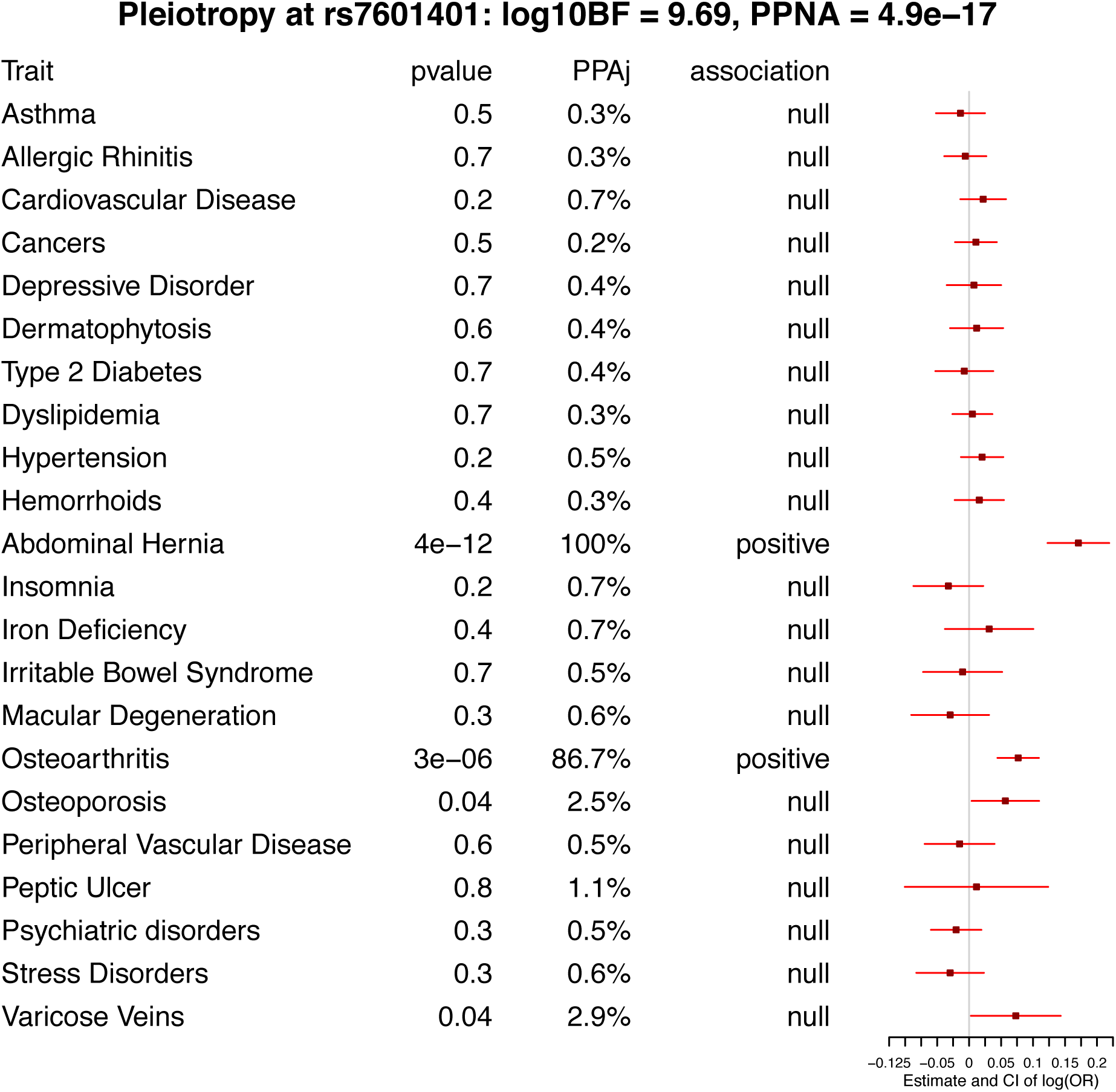
Forest plot for pleiotropic signal at rs7601401 detected by CPBayes.

**Figure S19:**
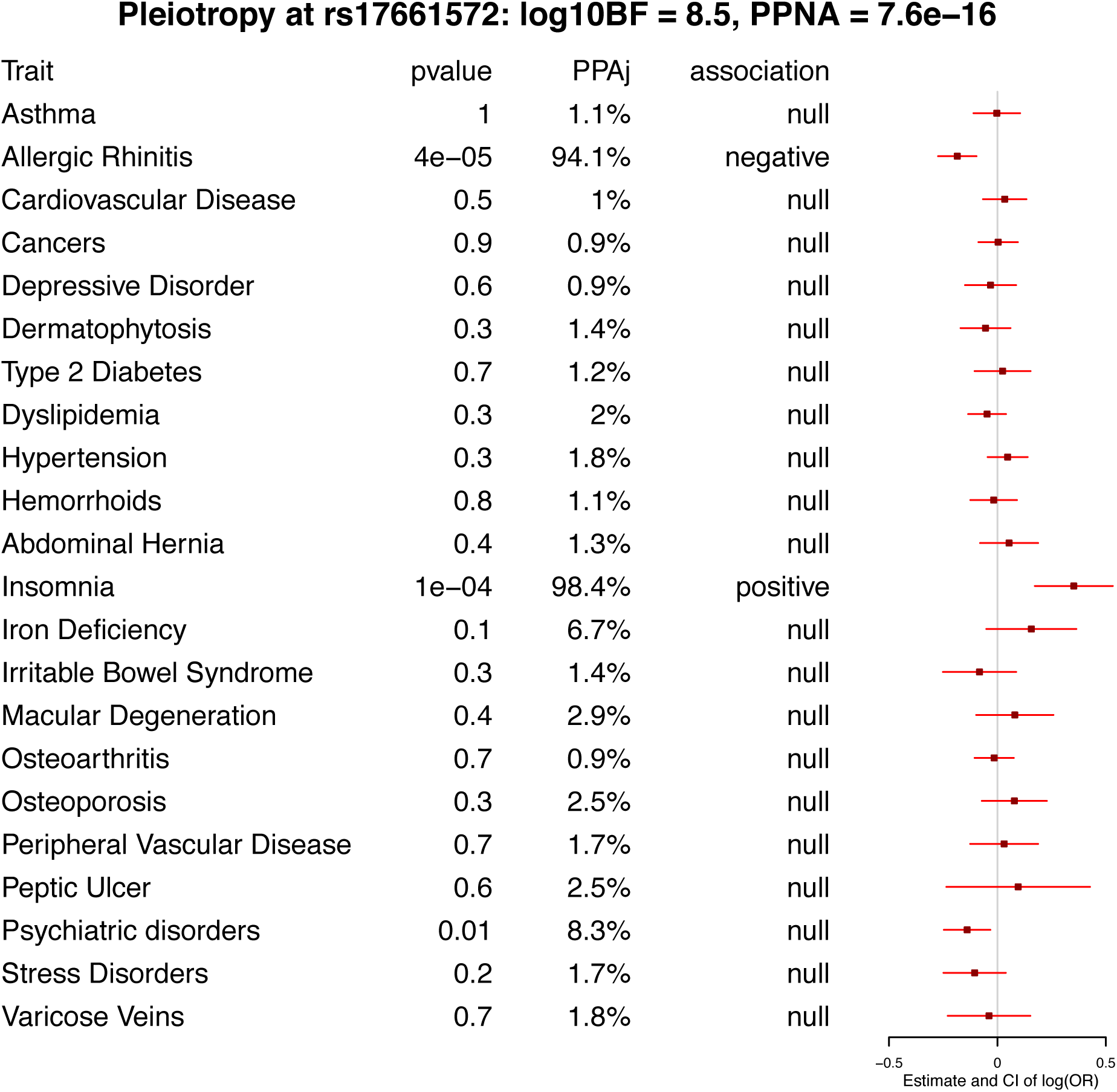
Forest plot for pleiotropic signal at rs17661572 detected by CPBayes.

**Figure S20:**
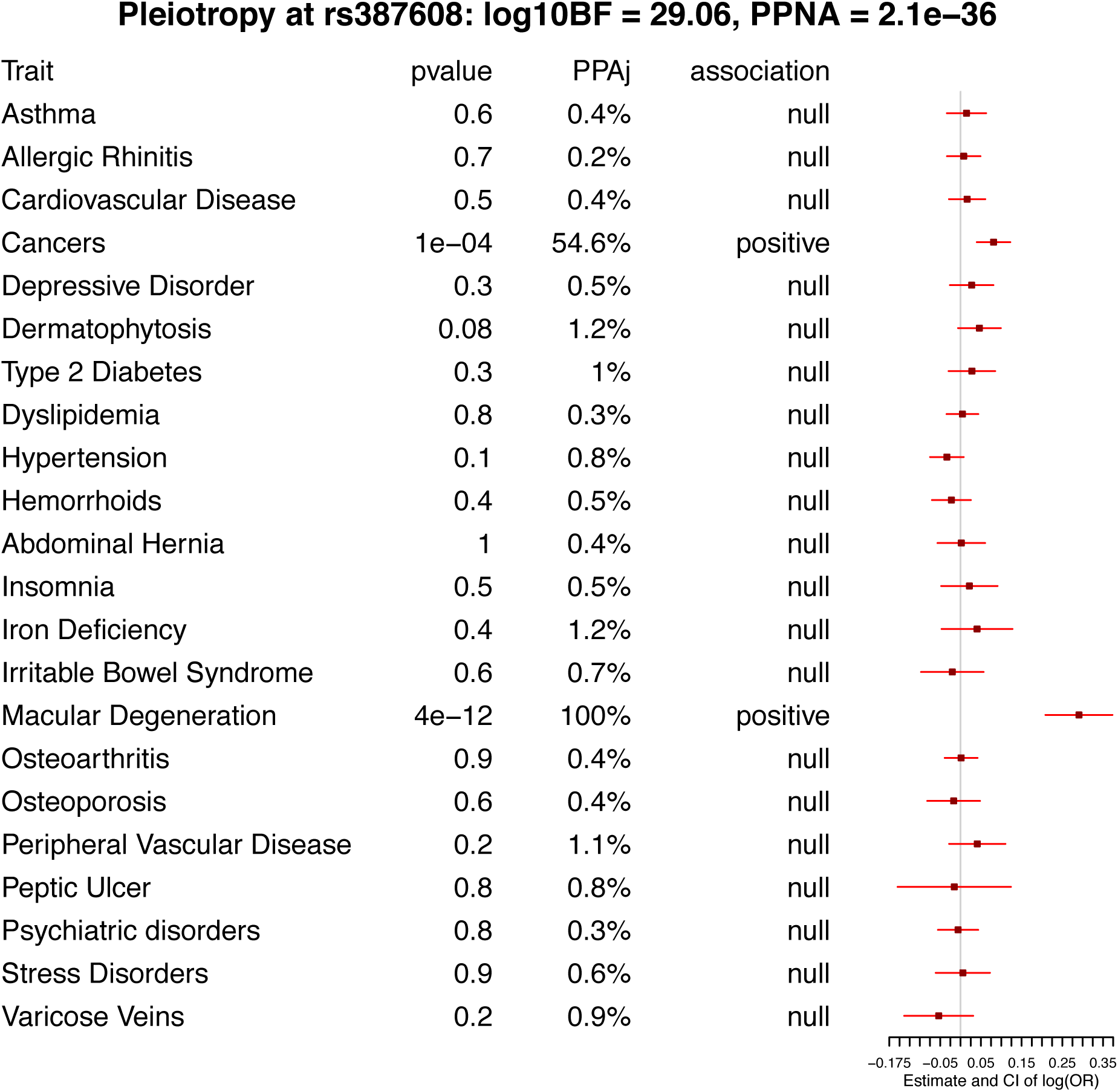
Forest plot for pleiotropic signal at rs387608 detected by CPBayes.

**Figure S21:**
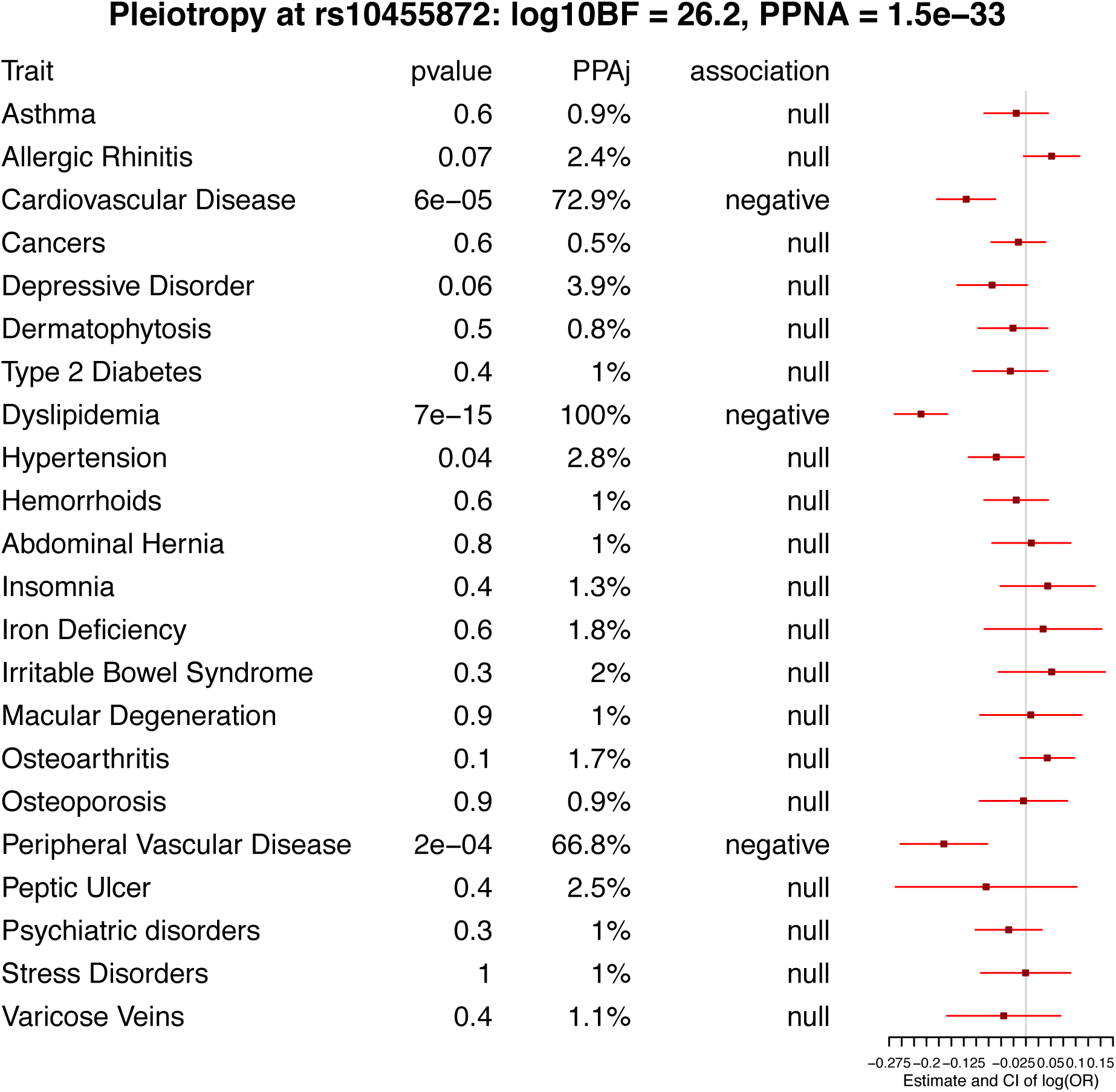
Forest plot for pleiotropic signal at rs10455872 detected by CPBayes.

**Figure S22:**
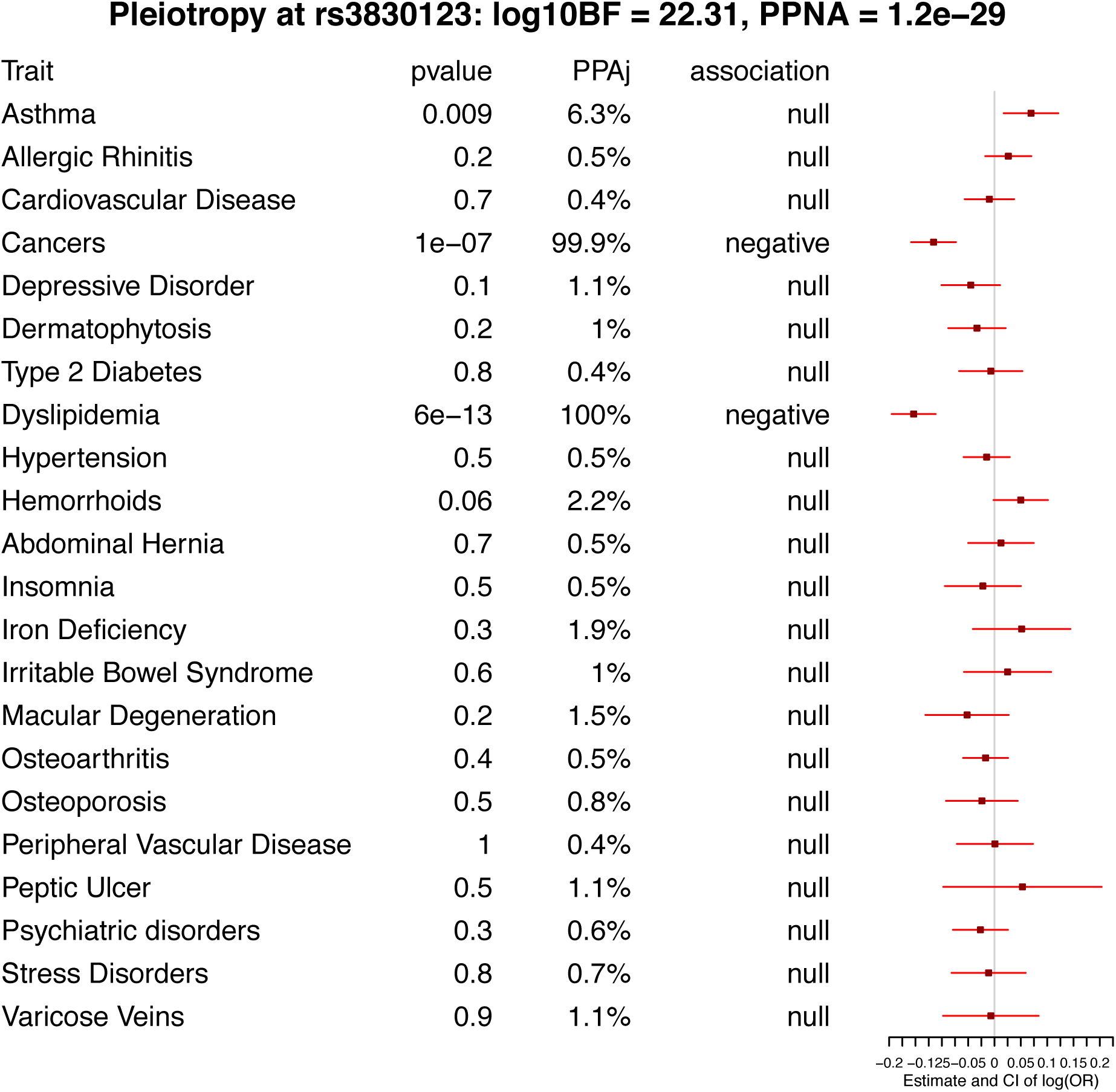
Forest plot for pleiotropic signal at rs3830123 detected by CPBayes.

**Figure S23:**
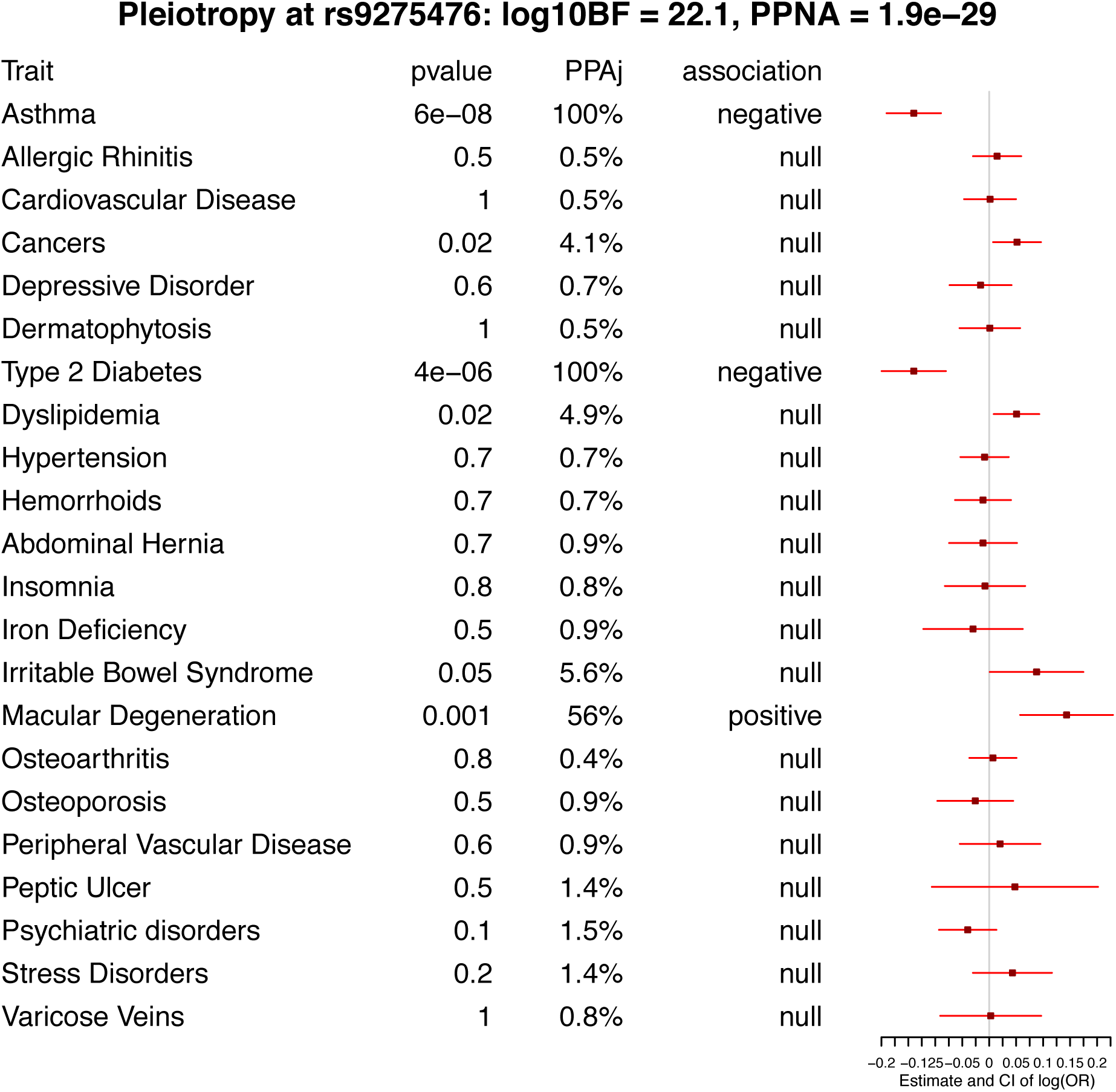
Forest plot for pleiotropic signal at rs9275476 detected by CPBayes.

**Figure S24:**
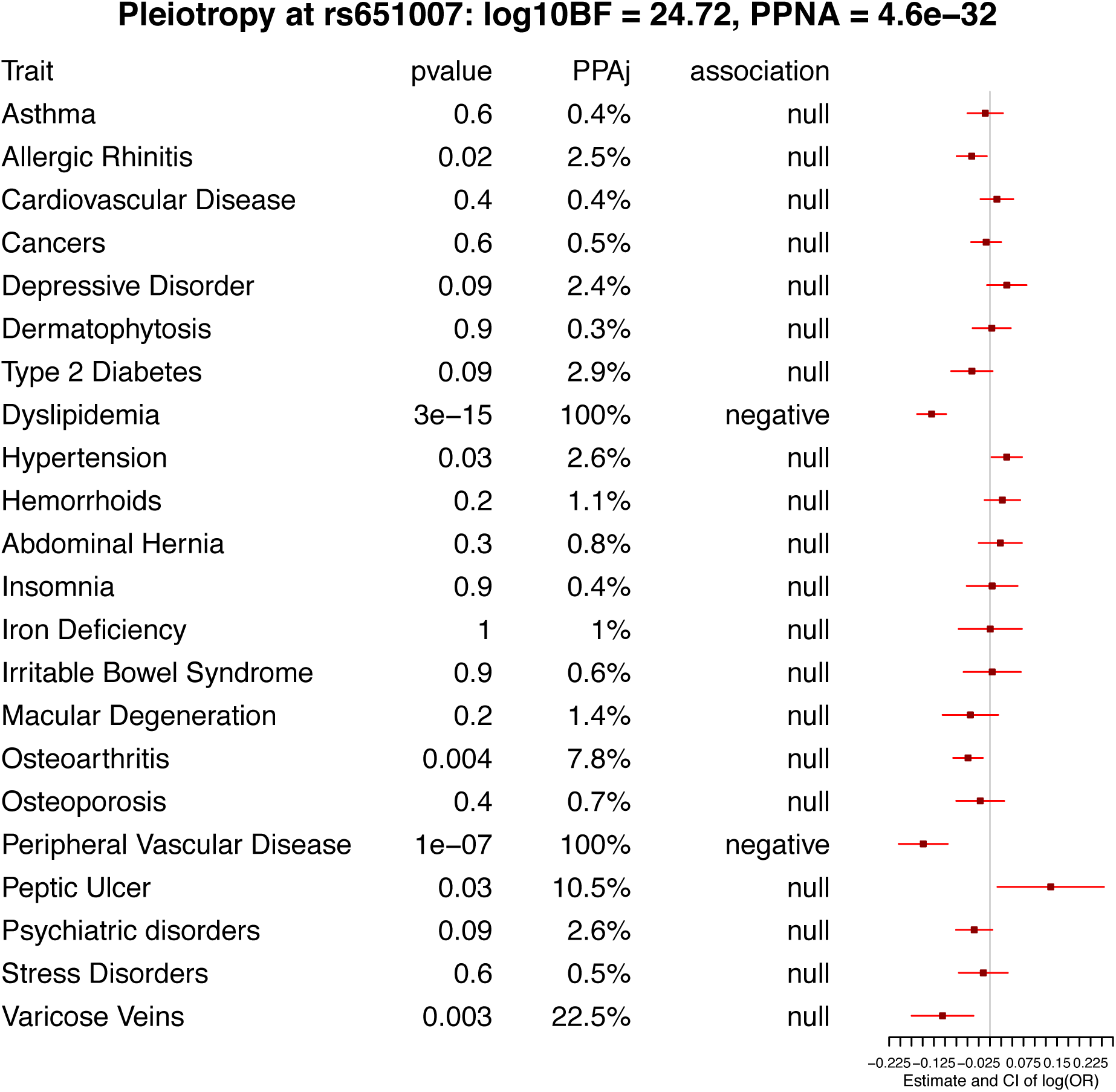
Forest plot for pleiotropic signal at rs651007 detected by CPBayes.

**Figure S25:**
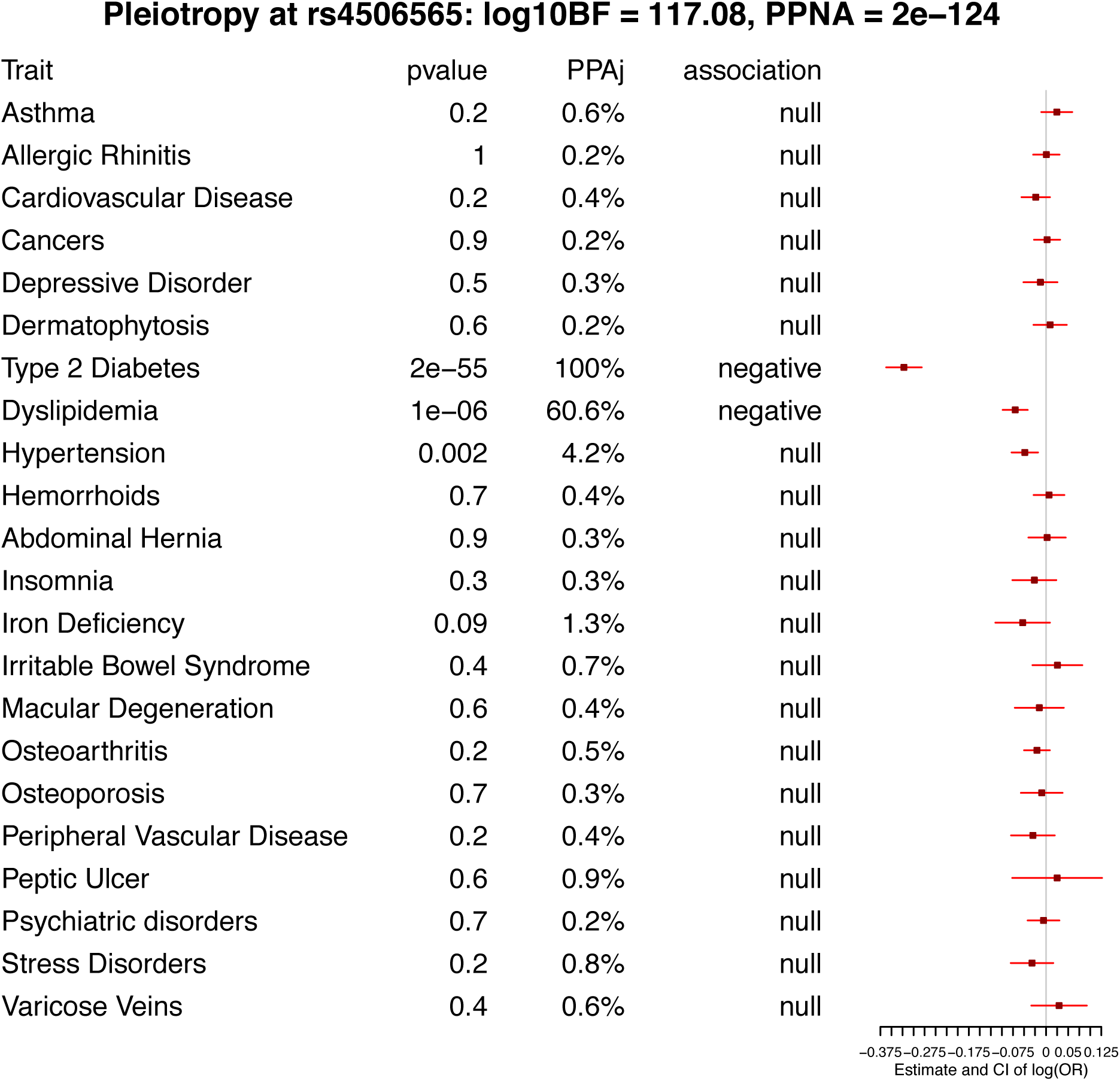
Forest plot for pleiotropic signal at rs4506565 detected by CPBayes.

## 15 Table presenting independent pleiotropic SNPs identified by CPBayes for which it selected one phenotype in GERA cohort

**Table S21:**
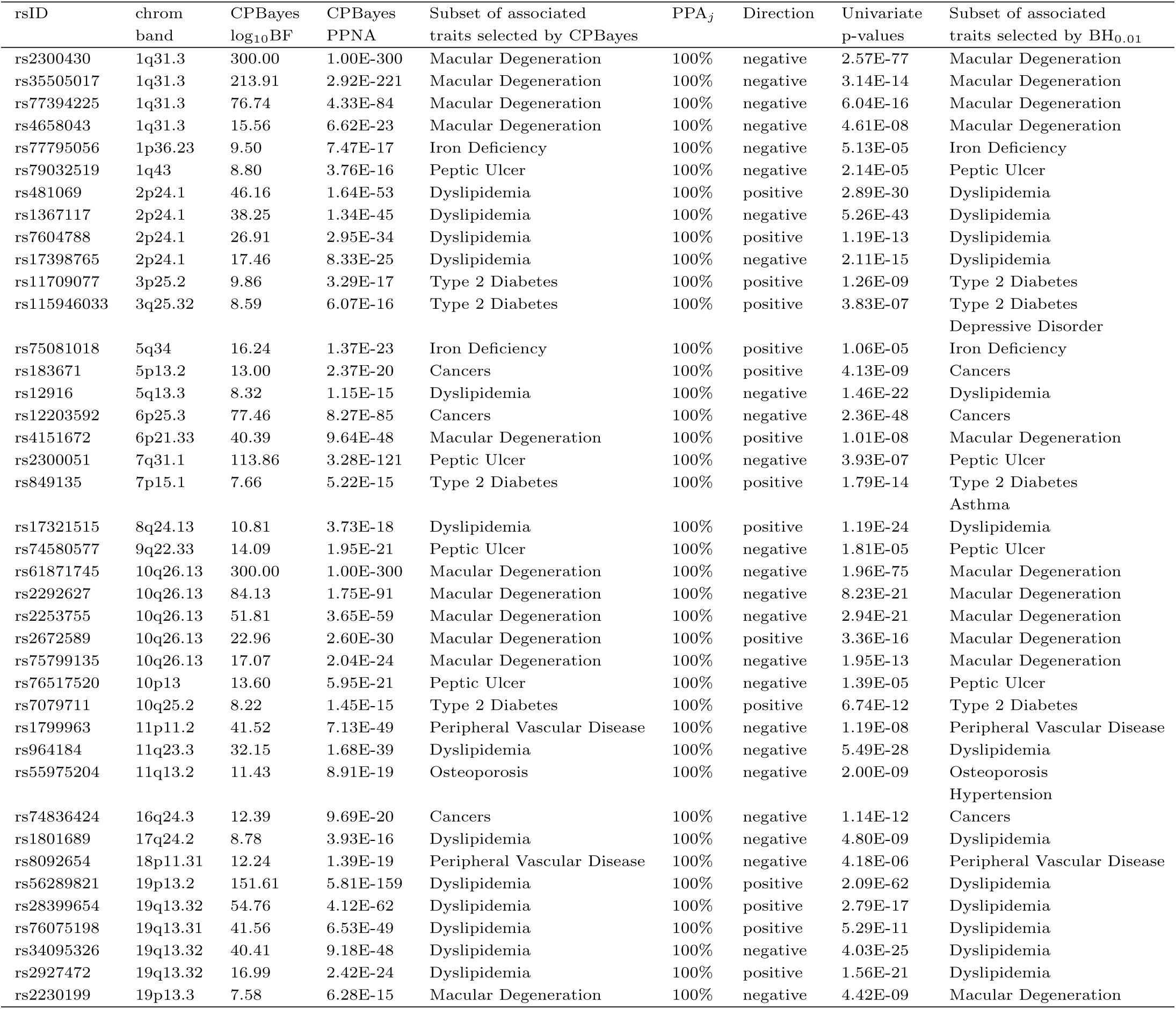
Independent pleiotropic SNPs identified by CPBayes for which one phenotype was selected.

| rsID | chrom<br>band | CPBayes<br>log<sub>10</sub>BF | CPBayes<br>PPNA | Subset of associated<br>traits selected by CPBayes | PPA<sub>j</sub> | Direction | Univariate<br>p-values | Subset of associated<br>traits selected by BH<sub>0.01</sub> |
| --- | --- | --- | --- | --- | --- | --- | --- | --- |
| rs2300430 | 1q31.3 | 300.00 | 1.00E-300 | Macular Degeneration | 100% | negative | 2.57E-77 | Macular Degeneration |
| rs35505017 | 1q31.3 | 213.91 | 2.92E-221 | Macular Degeneration | 100% | negative | 3.14E-14 | Macular Degeneration |
| rs77394225 | 1q31.3 | 76.74 | 4.33E-84 | Macular Degeneration | 100% | negative | 6.04E-16 | Macular Degeneration |
| rs4658043 | 1q31.3 | 15.56 | 6.62E-23 | Macular Degeneration | 100% | negative | 4.61E-08 | Macular Degeneration |
| rs77795056 | 1p36.23 | 9.50 | 7.47E-17 | Iron Deficiency | 100% | negative | 5.13E-05 | Iron Deficiency |
| rs79032519 | 1q43 | 8.80 | 3.76E-16 | Peptic Ulcer | 100% | negative | 2.14E-05 | Peptic Ulcer |
| rs481069 | 2p24.1 | 46.16 | 1.64E-53 | Dyslipidemia | 100% | positive | 2.89E-30 | Dyslipidemia |
| rs1367117 | 2p24.1 | 38.25 | 1.34E-45 | Dyslipidemia | 100% | negative | 5.26E-43 | Dyslipidemia |
| rs7604788 | 2p24.1 | 26.91 | 2.95E-34 | Dyslipidemia | 100% | positive | 1.19E-13 | Dyslipidemia |
| rs17398765 | 2p24.1 | 17.46 | 8.33E-25 | Dyslipidemia | 100% | negative | 2.11E-15 | Dyslipidemia |
| rs11709077 | 3p25.2 | 9.86 | 3.29E-17 | Type 2 Diabetes | 100% | positive | 1.26E-09 | Type 2 Diabetes |
| rs115946033 | 3q25.32 | 8.59 | 6.07E-16 | Type 2 Diabetes | 100% | positive | 3.83E-07 | Type 2 Diabetes<br>Depressive Disorder |
| rs75081018 | 5q34 | 16.24 | 1.37E-23 | Iron Deficiency | 100% | positive | 1.06E-05 | Iron Deficiency |
| rs183671 | 5p13.2 | 13.00 | 2.37E-20 | Cancers | 100% | positive | 4.13E-09 | Cancers |
| rs12916 | 5q13.3 | 8.32 | 1.15E-15 | Dyslipidemia | 100% | negative | 1.46E-22 | Dyslipidemia |
| rs12203592 | 6p25.3 | 77.46 | 8.27E-85 | Cancers | 100% | negative | 2.36E-48 | Cancers |
| rs4151672 | 6p21.33 | 40.39 | 9.64E-48 | Macular Degeneration | 100% | positive | 1.01E-08 | Macular Degeneration |
| rs2300051 | 7q31.1 | 113.86 | 3.28E-121 | Peptic Ulcer | 100% | negative | 3.93E-07 | Peptic Ulcer |
| rs849135 | 7p15.1 | 7.66 | 5.22E-15 | Type 2 Diabetes | 100% | positive | 1.79E-14 | Type 2 Diabetes<br>Asthma |
| rs17321515 | 8q24.13 | 10.81 | 3.73E-18 | Dyslipidemia | 100% | positive | 1.19E-24 | Dyslipidemia |
| rs74580577 | 9q22.33 | 14.09 | 1.95E-21 | Peptic Ulcer | 100% | negative | 1.81E-05 | Peptic Ulcer |
| rs61871745 | 10q26.13 | 300.00 | 1.00E-300 | Macular Degeneration | 100% | negative | 1.96E-75 | Macular Degeneration |
| rs2292627 | 10q26.13 | 84.13 | 1.75E-91 | Macular Degeneration | 100% | negative | 8.23E-21 | Macular Degeneration |
| rs2253755 | 10q26.13 | 51.81 | 3.65E-59 | Macular Degeneration | 100% | negative | 2.94E-21 | Macular Degeneration |
| rs2672589 | 10q26.13 | 22.96 | 2.60E-30 | Macular Degeneration | 100% | positive | 3.36E-16 | Macular Degeneration |
| rs75799135 | 10q26.13 | 17.07 | 2.04E-24 | Macular Degeneration | 100% | negative | 1.95E-13 | Macular Degeneration |
| rs76517520 | 10p13 | 13.60 | 5.95E-21 | Peptic Ulcer | 100% | negative | 1.39E-05 | Peptic Ulcer |
| rs7079711 | 10q25.2 | 8.22 | 1.45E-15 | Type 2 Diabetes | 100% | positive | 6.74E-12 | Type 2 Diabetes |
| rs1799963 | 11p11.2 | 41.52 | 7.13E-49 | Peripheral Vascular Disease | 100% | negative | 1.19E-08 | Peripheral Vascular Disease |
| rs964184 | 11q23.3 | 32.15 | 1.68E-39 | Dyslipidemia | 100% | negative | 5.49E-28 | Dyslipidemia |
| rs55975204 | 11q13.2 | 11.43 | 8.91E-19 | Osteoporosis | 100% | negative | 2.00E-09 | Osteoporosis<br>Hypertension |
| rs74836424 | 16q24.3 | 12.39 | 9.69E-20 | Cancers | 100% | negative | 1.14E-12 | Cancers |
| rs1801689 | 17q24.2 | 8.78 | 3.93E-16 | Dyslipidemia | 100% | negative | 4.80E-09 | Dyslipidemia |
| rs8092654 | 18p11.31 | 12.24 | 1.39E-19 | Peripheral Vascular Disease | 100% | negative | 4.18E-06 | Peripheral Vascular Disease |
| rs56289821 | 19p13.2 | 151.61 | 5.81E-159 | Dyslipidemia | 100% | positive | 2.09E-62 | Dyslipidemia |
| rs28399654 | 19q13.32 | 54.76 | 4.12E-62 | Dyslipidemia | 100% | positive | 2.79E-17 | Dyslipidemia |
| rs76075198 | 19q13.31 | 41.56 | 6.53E-49 | Dyslipidemia | 100% | positive | 5.29E-11 | Dyslipidemia |
| rs34095326 | 19q13.32 | 40.41 | 9.18E-48 | Dyslipidemia | 100% | negative | 4.03E-25 | Dyslipidemia |
| rs2927472 | 19q13.32 | 16.99 | 2.42E-24 | Dyslipidemia | 100% | positive | 1.56E-21 | Dyslipidemia |
| rs2230199 | 19p13.3 | 7.58 | 6.28E-15 | Macular Degeneration | 100% | negative | 4.42E-09 | Macular Degeneration |

## 16 Tables presenting independent pleiotropic signals detected by ASSET in GERA cohort

In the next four tables, we present the independent pleiotropic signals detected by ASSET in the GERA cohort.

**Table S22:** Independent pleiotropic signals detected by ASSET for chromosome 1-2

| rsID | chrom<br>band | ASSET<br>p-value | Subset of associated<br>traits selected by ASSET | Univariate<br>p-values | Subset of associated<br>traits selected by BH<sub>0.01</sub> |
| --- | --- | --- | --- | --- | --- |
| rs2300430 | 1q31.3 | 1.52E-72 | Allergic Rhinitis | 0.34 | Macular Degeneration |
|  |  |  | Depressive Disorder | 0.18 |  |
|  |  |  | Dermatophytosis | 0.09 |  |
|  |  |  | Hemorrhoids | 0.36 |  |
|  |  |  | Insomnia | 0.47 |  |
|  |  |  | Macular Degeneration | 2.57E-77 |  |
|  |  |  | Osteoporosis | 0.09 |  |
|  |  |  | Peptic Ulcer | 0.74 |  |
| rs10494745 | 1q31.3 | 1.05E-14 | Cancers | 0.04 | Macular Degeneration |
|  |  |  | Dermatophytosis | 0.27 |  |
|  |  |  | Type 2 Diabetes | 0.13 |  |
|  |  |  | Insomnia | 0.03 |  |
|  |  |  | Iron Deficiency | 0.28 |  |
|  |  |  | Macular Degeneration | 6.15E-21 |  |
|  |  |  | Peripheral Vascular Disease | 0.24 |  |
|  |  |  | Peptic Ulcer | 0.35 |  |
| rs6025 | 1q24.2 | 1.38E-11 | Dermatophytosis | 0.0018 | Dermatophytosis |
|  |  |  | Hemorrhoids | 0.0014 | Hemorrhoids |
|  |  |  | Iron Deficiency | 0.0004 | Iron Deficiency |
|  |  |  | Osteoporosis | 0.0002 | Osteoporosis |
|  |  |  | Peripheral Vascular Disease | 6.81E-14 | Peripheral Vascular Disease |
| rs77394225 | 1q31.3 | 6.84E-10 | Dermatophytosis | 0.62 | Macular Degeneration |
|  |  |  | Dyslipidemia | 0.47 |  |
|  |  |  | Abdominal Hernia | 0.65 |  |
|  |  |  | Macular Degeneration | 6.04E-16 |  |
|  |  |  | Osteoarthritis | 0.30 |  |
|  |  |  | Osteoporosis | 0.34 |  |
|  |  |  | Peptic Ulcer | 0.46 |  |
|  |  |  | Psychiatric disorders | 0.39 |  |
|  |  |  | Stress Disorders | 0.32 |  |
| rs35505017 | 1q31.3 | 3.18E-08 | Dermatophytosis | 0.56 | Macular Degeneration |
|  |  |  | Dyslipidemia | 0.40 |  |
|  |  |  | Macular Degeneration | 3.14E-14 |  |
|  |  |  | Peptic Ulcer | 0.63 |  |
|  |  |  | Stress Disorders | 0.21 |  |
| rs1367117 | 2p24.1 | 5.19E-34 | Depressive Disorder | 0.61 | Dyslipidemia |
|  |  |  | Dyslipidemia | 5.26E-43 |  |
|  |  |  | Hemorrhoids | 0.67 |  |
|  |  |  | Abdominal Hernia | 0.34 |  |
|  |  |  | Macular Degeneration | 0.50 |  |
|  |  |  | Peptic Ulcer | 0.57 |  |
|  |  |  | Varicose Veins | 0.70 |  |
| rs562338 | 2p24.1 | 3.15E-27 | Cardiovascular Disease | 0.26 | Dyslipidemia |
|  |  |  | Dyslipidemia | 3.43E-35 |  |
|  |  |  | Abdominal Hernia | 0.18 |  |
|  |  |  | Peptic Ulcer | 0.04 |  |
|  |  |  | Psychiatric disorders | 0.22 |  |
| rs79281791 | 2p24.1 | 9.75E-10 | Dyslipidemia | 4.91E-17 | Dyslipidemia |
|  |  |  | Insomnia | 0.007 |  |
|  |  |  | Iron Deficiency | 0.04 |  |
|  |  |  | Macular Degeneration | 0.17 |  |
|  |  |  | Peptic Ulcer | 0.31 |  |
|  |  |  | Varicose Veins | 0.13 |  |
| rs3791679 | 2p16.1 | 3.73E-09 | Type 2 Diabetes | 0.35 | Abdominal Hernia<br>Osteoarthritis |
|  |  |  | Abdominal Hernia | 2.66E-14 |  |
|  |  |  | Insomnia | 0.25 |  |
|  |  |  | Peripheral Vascular Disease | 0.25 |  |
|  |  |  | Psychiatric disorders | 0.11 |  |
|  |  |  | Stress Disorders | 0.24 |  |
|  |  |  | Varicose Veins | 0.002 |  |

**Table S23:**
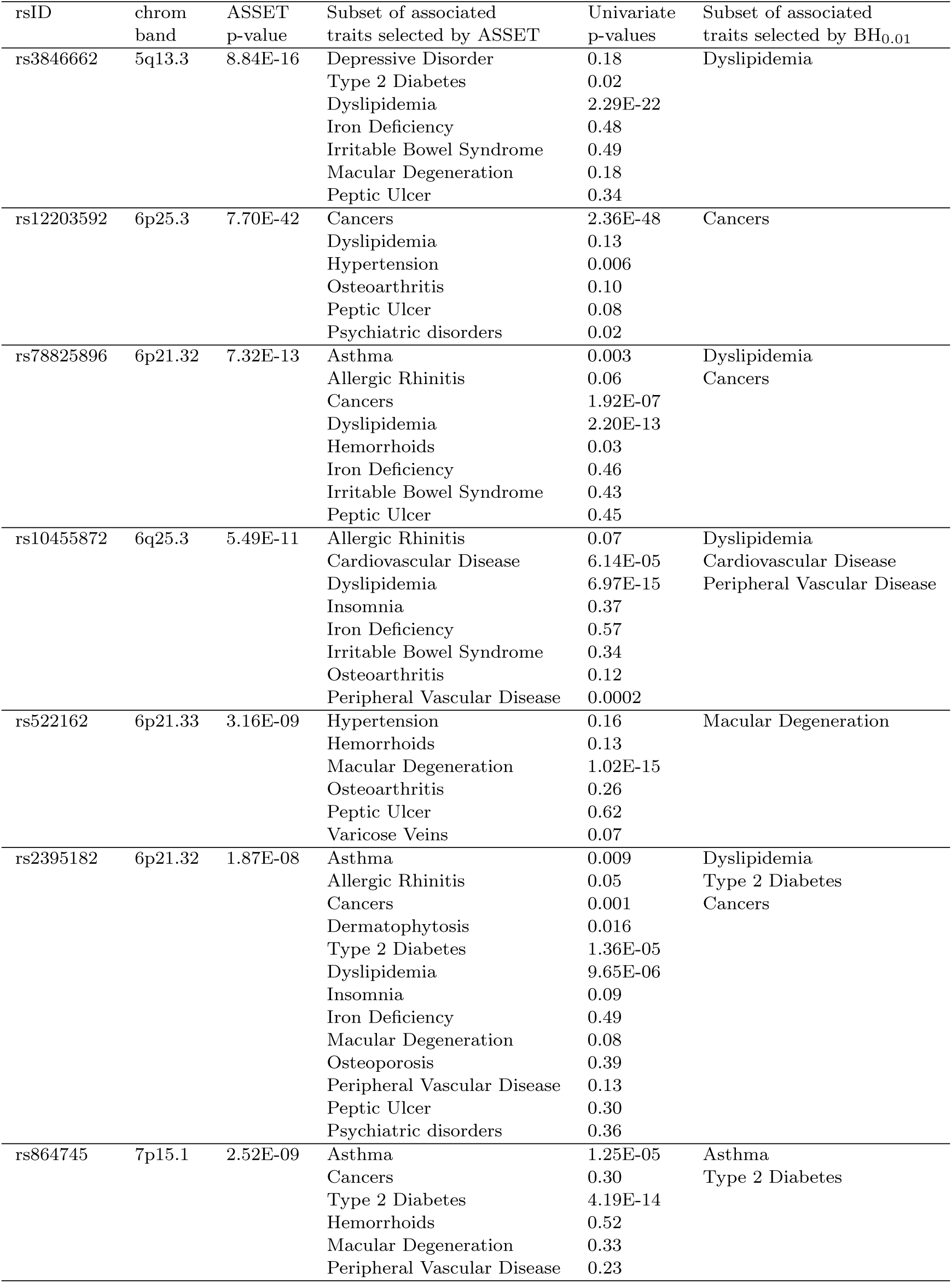
Independent pleiotropic signals detected by ASSET for chromosome 3-7

**Table S24:** Independent pleiotropic signals detected by ASSET for chromosome 8-10

| rsID | chrom<br>band | ASSET<br>p-value | Subset of associated<br>traits selected by ASSET | Univariate<br>p-values | Subset of associated<br>traits selected by BH<sub>0.01</sub> |
| --- | --- | --- | --- | --- | --- |
| rs2001945 | 8q24.13 | 4.01E-19 | Asthma | 0.43 | Dyslipidemia |
|  |  |  | Allergic Rhinitis | 0.26 |  |
|  |  |  | Depressive Disorder | 0.39 |  |
|  |  |  | Dyslipidemia | 1.86E-25 |  |
|  |  |  | Abdominal Hernia | 0.37 |  |
|  |  |  | Iron Deficiency | 0.61 |  |
|  |  |  | Macular Degeneration | 0.07 |  |
|  |  |  | Osteoporosis | 0.39 |  |
|  |  |  | Stress Disorders | 0.30 |  |
| rs651007 | 9q34.2 | 3.09E-16 | Depressive Disorder | 0.09 | Dyslipidemia<br>Peripheral Vascular Disease |
|  |  |  | Dyslipidemia | 2.89E-15 |  |
|  |  |  | Hypertension | 0.03 |  |
|  |  |  | Hemorrhoids | 0.17 |  |
|  |  |  | Abdominal Hernia | 0.34 |  |
|  |  |  | Peripheral Vascular Disease | 9.55E-08 |  |
|  |  |  | Peptic Ulcer | 0.03 |  |
|  |  |  | Varicose Veins | 0.0027 |  |
| rs61871745 | 10q26.13 | 6.99E-69 | Depressive Disorder | 0.04 | Macular Degeneration |
|  |  |  | Hemorrhoids | 0.05 |  |
|  |  |  | Insomnia | 0.16 |  |
|  |  |  | Macular Degeneration | 1.96E-75 |  |
|  |  |  | Stress Disorders | 0.01 |  |
| rs4506565 | 10q25.2 | 1.48E-46 | Asthma | 0.18 | Type 2 Diabetes<br>Dyslipidemia |
|  |  |  | Type 2 Diabetes | 2.02E-55 |  |
|  |  |  | Irritable Bowel Syndrome | 0.37 |  |
|  |  |  | Peptic Ulcer | 0.63 |  |
|  |  |  | Varicose Veins | 0.36 |  |
| rs2292627 | 10q26.13 | 9.71E-15 | Type 2 Diabetes | 0.06 | Macular Degeneration |
|  |  |  | Hemorrhoids | 0.34 |  |
|  |  |  | Insomnia | 0.29 |  |
|  |  |  | Irritable Bowel Syndrome | 0.54 |  |
|  |  |  | Macular Degeneration | 8.23E-21 |  |
|  |  |  | Peptic Ulcer | 0.54 |  |
|  |  |  | Stress Disorders | 0.36 |  |
|  |  |  | Varicose Veins | 0.65 |  |
| rs2253755 | 10q26.13 | 3.65E-14 | Allergic Rhinitis | 0.18 | Macular Degeneration |
|  |  |  | Cardiovascular Disease | 0.29 |  |
|  |  |  | Dermatophytosis | 0.37 |  |
|  |  |  | Type 2 Diabetes | 0.14 |  |
|  |  |  | Macular Degeneration | 2.94E-21 |  |
|  |  |  | Psychiatric disorders | 0.26 |  |
|  |  |  | Stress Disorders | 0.39 |  |
| rs2672589 | 10q26.13 | 1.70E-10 | Iron Deficiency | 0.17 | Macular Degeneration |
|  |  |  | Irritable Bowel Syndrome | 0.29 |  |
|  |  |  | Macular Degeneration | 3.36E-16 |  |
|  |  |  | Peptic Ulcer | 0.058 |  |
|  |  |  | Stress Disorders | 0.02 |  |
| rs75799135 | 10q26.13 | 2.45E-08 | Depressive Disorder | 0.04 | Macular Degeneration |
|  |  |  | Type 2 Diabetes | 0.029 |  |
|  |  |  | Hemorrhoids | 0.088 |  |
|  |  |  | Insomnia | 0.037 |  |
|  |  |  | Macular Degeneration | 1.95E-13 |  |
|  |  |  | Stress Disorders | 0.19 |  |

**Table S25:**
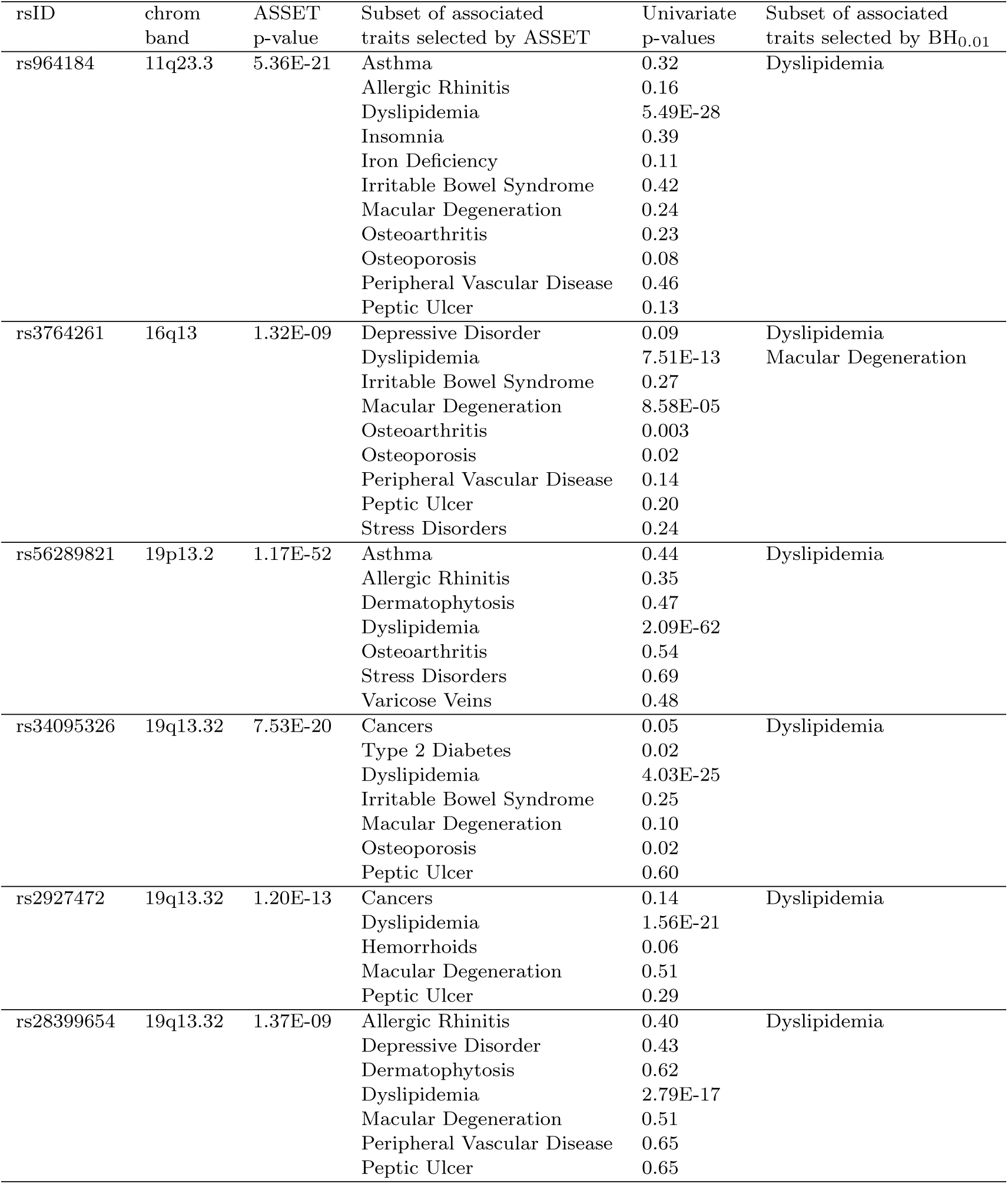
Independent pleiotropic signals detected by ASSET for chromosome 11-22

**Table S26:** Pair-wise trait-trait correlation matrix in GERA cohort.

|  | astm | alrg | card | canc | dprs | drmt | dia2 | dysl | hypr | hemr | hern | insm | iron | irrb | mcdg | osta | ostp | pvd | pept | psyc | strs | vrvn |
| --- | --- | --- | --- | --- | --- | --- | --- | --- | --- | --- | --- | --- | --- | --- | --- | --- | --- | --- | --- | --- | --- | --- |
| astm | 1.00 | 0.27 | 0.06 | 0.00 | 0.09 | 0.03 | 0.05 | 0.04 | 0.06 | 0.03 | 0.03 | 0.05 | 0.05 | 0.06 | 0.02 | 0.08 | 0.02 | 0.05 | 0.02 | 0.09 | 0.08 | 0.02 |
| alrg | 0.27 | 1.00 | 0.02 | 0.00 | 0.08 | 0.05 | 0.01 | 0.03 | 0.03 | 0.04 | 0.02 | 0.06 | 0.02 | 0.09 | 0.01 | 0.08 | 0.02 | 0.02 | 0.02 | 0.09 | 0.09 | 0.03 |
| card | 0.06 | 0.02 | 1.00 | 0.16 | 0.01 | 0.08 | 0.16 | 0.27 | 0.32 | 0.05 | 0.12 | 0.06 | 0.09 | 0.02 | 0.14 | 0.18 | 0.07 | 0.22 | 0.09 | 0.03 | 0.02 | 0.03 |
| canc | 0.00 | 0.00 | 0.16 | 1.00 | -0.02 | 0.07 | 0.05 | 0.10 | 0.14 | 0.06 | 0.10 | 0.03 | 0.05 | 0.01 | 0.10 | 0.11 | 0.07 | 0.14 | 0.04 | -0.01 | 0.00 | 0.03 |
| dprs | 0.09 | 0.08 | 0.01 | -0.02 | 1.00 | 0.03 | 0.03 | 0.02 | 0.01 | 0.03 | 0.00 | 0.13 | 0.04 | 0.09 | 0.00 | 0.06 | 0.01 | 0.03 | 0.02 | 0.35 | 0.17 | 0.01 |
| drmt | 0.03 | 0.05 | 0.08 | 0.07 | 0.03 | 1.00 | 0.08 | 0.06 | 0.07 | 0.04 | 0.06 | 0.04 | 0.03 | 0.02 | 0.04 | 0.10 | 0.01 | 0.07 | 0.02 | 0.03 | 0.04 | 0.05 |
| dia2 | 0.05 | 0.01 | 0.16 | 0.05 | 0.03 | 0.08 | 1.00 | 0.26 | 0.28 | 0.01 | 0.04 | 0.01 | 0.07 | -0.01 | 0.05 | 0.08 | -0.03 | 0.08 | 0.04 | 0.01 | 0.03 | 0.00 |
| dysl | 0.04 | 0.03 | 0.27 | 0.10 | 0.02 | 0.06 | 0.26 | 1.00 | 0.36 | 0.06 | 0.07 | 0.04 | 0.04 | 0.01 | 0.07 | 0.16 | 0.04 | 0.12 | 0.05 | 0.02 | 0.03 | 0.01 |
| hypr | 0.06 | 0.03 | 0.32 | 0.14 | 0.01 | 0.07 | 0.28 | 0.36 | 1.00 | 0.05 | 0.09 | 0.04 | 0.07 | 0.02 | 0.13 | 0.21 | 0.05 | 0.15 | 0.06 | 0.02 | 0.04 | 0.02 |
| hemr | 0.03 | 0.04 | 0.05 | 0.06 | 0.03 | 0.04 | 0.01 | 0.06 | 0.05 | 1.00 | 0.06 | 0.04 | 0.05 | 0.07 | 0.02 | 0.06 | 0.01 | 0.04 | 0.03 | 0.04 | 0.03 | 0.02 |
| hern | 0.03 | 0.02 | 0.12 | 0.10 | 0.00 | 0.06 | 0.04 | 0.07 | 0.09 | 0.06 | 1.00 | 0.02 | 0.08 | 0.01 | 0.05 | 0.07 | 0.00 | 0.09 | 0.06 | 0.00 | -0.01 | 0.02 |
| insm | 0.05 | 0.06 | 0.06 | 0.03 | 0.13 | 0.04 | 0.01 | 0.04 | 0.04 | 0.04 | 0.02 | 1.00 | 0.04 | 0.08 | 0.03 | 0.08 | 0.04 | 0.06 | 0.03 | 0.15 | 0.10 | 0.02 |
| iron | 0.05 | 0.02 | 0.09 | 0.05 | 0.04 | 0.03 | 0.07 | 0.04 | 0.07 | 0.05 | 0.08 | 0.04 | 1.00 | 0.03 | 0.04 | 0.06 | 0.03 | 0.09 | 0.10 | 0.04 | 0.03 | 0.02 |
| irrb | 0.06 | 0.09 | 0.02 | 0.01 | 0.09 | 0.02 | -0.01 | 0.01 | 0.02 | 0.07 | 0.01 | 0.08 | 0.03 | 1.00 | 0.02 | 0.07 | 0.05 | 0.01 | 0.03 | 0.14 | 0.11 | 0.02 |
| mcdg | 0.02 | 0.01 | 0.14 | 0.10 | 0.00 | 0.04 | 0.05 | 0.07 | 0.13 | 0.02 | 0.05 | 0.03 | 0.04 | 0.02 | 1.00 | 0.11 | 0.09 | 0.10 | 0.03 | 0.00 | 0.00 | 0.02 |
| osta | 0.08 | 0.08 | 0.18 | 0.11 | 0.06 | 0.10 | 0.08 | 0.16 | 0.21 | 0.06 | 0.07 | 0.08 | 0.06 | 0.07 | 0.11 | 1.00 | 0.11 | 0.13 | 0.06 | 0.04 | 0.07 | 0.07 |
| ostp | 0.02 | 0.02 | 0.07 | 0.07 | 0.01 | 0.01 | -0.03 | 0.04 | 0.05 | 0.01 | 0.00 | 0.04 | 0.03 | 0.05 | 0.09 | 0.11 | 1.00 | 0.08 | 0.02 | 0.03 | 0.03 | 0.04 |
| pvd | 0.05 | 0.02 | 0.22 | 0.14 | 0.03 | 0.07 | 0.08 | 0.12 | 0.15 | 0.04 | 0.09 | 0.06 | 0.09 | 0.01 | 0.10 | 0.13 | 0.08 | 1.00 | 0.06 | 0.03 | 0.02 | 0.04 |
| pept | 0.02 | 0.02 | 0.09 | 0.04 | 0.02 | 0.02 | 0.04 | 0.05 | 0.06 | 0.03 | 0.06 | 0.03 | 0.10 | 0.03 | 0.03 | 0.06 | 0.02 | 0.06 | 1.00 | 0.02 | 0.01 | 0.01 |
| psyc | 0.09 | 0.09 | 0.03 | -0.01 | 0.35 | 0.03 | 0.01 | 0.02 | 0.02 | 0.04 | 0.00 | 0.15 | 0.04 | 0.14 | 0.00 | 0.04 | 0.03 | 0.03 | 0.02 | 1.00 | 0.24 | 0.02 |
| strs | 0.08 | 0.09 | 0.02 | 0.00 | 0.17 | 0.04 | 0.03 | 0.03 | 0.04 | 0.03 | -0.01 | 0.10 | 0.03 | 0.11 | 0.00 | 0.07 | 0.03 | 0.02 | 0.01 | 0.24 | 1.00 | 0.03 |
| vrvn | 0.02 | 0.03 | 0.03 | 0.03 | 0.01 | 0.05 | 0.00 | 0.01 | 0.02 | 0.02 | 0.02 | 0.02 | 0.02 | 0.02 | 0.02 | 0.07 | 0.04 | 0.04 | 0.01 | 0.02 | 0.03 | 1.00 |

Legend: Phenotypes are abbreviated as: astm - Asthma, alrg - Allergic Rhinitis, card - Cardiovascular Disease, canc - Cancers, dprs - Depressive Disorder, drmt - Dermatophytosis, dia2 - Type 2 Diabetes, dysl - Dyslipidemia, hypr - Hypertension, hemr - Hemorrhoids, hern - Abdominal Hernia, insm - Insomnia, iron - Iron Deficiency, irrb - Irritable Bowel Syndrome, mcdg - Macular Degeneration, osta - Osteoarthritis, ostp - Osteoporosis, pvd - Peripheral Vascular Disease, pept - Peptic Ulcer, psyc - Psychiatric disorders, strs - Stress Disorders, vrvn - Varicose Veins.

## References

Andreassen, O. A., Djurovic, S., Thompson, W. K., Schork, A. J., Kendler, K. S., O?Donovan, M. C., Rujescu, D., Werge, T., van de Bunt, M., Morris, A. P., et al. (2013a). Improved detection of common variants associated with schizophrenia by leveraging pleiotropy with cardiovascular-disease risk factors. The American Journal of Human Genetics, 92(2): 197–209.

Andreassen, O. A., Thompson, W. K., Schork, A. J., Ripke, S., Mattingsdal, M., Kelsoe, J. R., Kendler, K. S., O’Donovan, M. C., Rujescu, D., Werge, T., et al. (2013b). Improved detection of common variants associated with schizophrenia and bipolar disorder using pleiotropy-informed conditional false discovery rate. PLoS Genet, 9(4): e1003455.

Benjamini, Y. and Hochberg, Y. (1995). Controlling the false discovery rate: a practical and powerful approach to multiple testing. Journal of the Royal Statistical Society. Series B (Methodological), pages 289–300.

Benjamini, Y. and Yekutieli, D. (2001). The control of the false discovery rate in multiple testing under dependency. Annals of statistics, pages 1165–1188.

Bhattacharjee, S., Rajaraman, P., Jacobs, K. B., Wheeler, W. A., Melin, B. S., Hartge, P., Yeager, M., Chung, C. C., Chanock, S. J., Chatterjee, N., et al. (2012). A subset-based approach improves power and interpretation for the combined analysis of genetic association studies of heterogeneous traits. The American Journal of Human Genetics, 90(5): 821–835.

Carty, C. L., Bhattacharjee, S., Haessler, J., Cheng, I., Hindorff, L. A., Aroda, V., Carlson, C. S., Hsu, C.-N., Wilkens, L., Liu, S., et al. (2014). Comparative analysis of metabolic syndrome components in over 15,000 african americans identifies pleiotropic variants: Results from the page study. Circulation: Cardiovascular Genetics, pages 505–513.

Chung, D., Yang, C., Li, C., Gelernter, J., and Zhao, H. (2014). Gpa: a statistical approach to prioritizing gwas results by integrating pleiotropy and annotation. PLoS Genet, 10(11): e1004787.

Efron, B. (2007). Size, power and false discovery rates. The Annals of Statistics, pages 1351–1377.

Efron, B. (2012). Large-scale inference: empirical Bayes methods for estimation, testing, and prediction, volume 1. Cambridge University Press.

Ellinghaus, D., Jostins, L., Spain, S. L., Cortes, A., Bethune, J., Han, B., Park, Y. R., Raychaudhuri, S., Pouget, J. G., Hübenthal, M., et al. (2016). Analysis of five chronic inflammatory diseases identifies 27 new associations and highlights disease-specific patterns at shared loci. Nature genetics.

Galesloot, T. E., van Steen, K., Kiemeney, L. A., Janss, L. L., and Vermeulen, S. H. (2014). A comparison of multivariate genome-wide association methods. PloS one, 9(4): e95923.

Gamazon, E. R., Wheeler, H. E., Shah, K. P., Mozaffari, S. V., Aquino-Michaels, K., Carroll, R. J., Eyler, A. E., Denny, J. C., Nicolae, D. L., Cox, N. J., et al. (2015). A gene-based association method for mapping traits using reference transcriptome data. Nature genetics, 47(9): 1091–1098.

George, E. I. and McCulloch, R. E. (1993). Variable selection via gibbs sampling. Journal of the American Statistical Association, 88(423): 881–889.

Giambartolomei, C., Vukcevic, D., Schadt, E. E., Franke, L., Hingorani, A. D., Wallace, C., and Plagnol, V. (2014). Bayesian test for colocalisation between pairs of genetic association studies using summary statistics. PLoS Genet, 10(5): e1004383.

Griffin, J. E., Brown, P. J., et al. (2010). Inference with normal-gamma prior distributions in regression problems. Bayesian Analysis, 5(1): 171–188.

Gu, F., Pfeiffer, R., Bhattacharjee, S., Han, S., Taylor, P., Berndt, S., Yang, H., Sigurdson, A., Toro, J., Mirabello, L., et al. (2013). Common genetic variants in the 9p21 region and their associations with multiple tumours. British journal of cancer, 108(6): 1378–1386.

Han, B. and Eskin, E. (2012). Interpreting meta-analyses of genome-wide association studies. PLoS Genet, 8(3): e1002555.

Holland, D., Wang, Y., Thompson, W. K., Schork, A., Chen, C.-H., Lo, M.-T., Witoelar, A., Werge, T., O’Donovan, M., Andreassen, O. A., et al. (2016). Estimating effect sizes and expected replication probabilities from gwas summary statistics. Frontiers in genetics, 7.

Ishwaran, H. and Rao, J. S. (2005). Spike and slab variable selection: frequentist and bayesian strategies. Annals of Statistics, pages 730–773.

Kar, S. P., Beesley, J., Al Olama, A. A., Michailidou, K., Tyrer, J., Kote-Jarai, Z., Lawrenson, K., Lindstrom, S., Ramus, S. J., Thompson, D. J., et al. (2016). Genome-wide meta-analyses of breast, ovarian, and prostate cancer association studies identify multiple new susceptibility loci shared by at least two cancer types. Cancer discovery, 6(9): 1052–1067.

Liley, J. and Wallace, C. (2015). A pleiotropy-informed bayesian false discovery rate adapted to a shared control design finds new disease associations from gwas summary statistics. PLoS Genet, 11(2): e1004926.

Lin, D.-Y. and Sullivan, P. F. (2009). Meta-analysis of genome-wide association studies with overlapping subjects. The American Journal of Human Genetics, 85(6): 862–872.

Majumdar, A., Haldar, T., and Witte, J. S. (2016). Determining which phenotypes underlie a pleiotropic signal. Genetic epidemiology.

Malsiner-Walli, G. and Wagner, H. (2011). Comparing spike and slab priors for bayesian variable selection. Austrian Journal of Statistics, 40(4): 241–264.

Mitchell, T. J. and Beauchamp, J. J. (1988). Bayesian variable selection in linear regression. Journal of the American Statistical Association, 83(404): 1023–1032.

Parkes, M., Cortes, A., van Heel, D. A., and Brown, M. A. (2013). Genetic insights into common pathways and complex relationships among immune-mediated diseases. Nature Reviews Genetics, 14(9): 661–673.

Pickrell, J. K., Berisa, T., Liu, J. Z., Ségurel, L., Tung, J. Y., and Hinds, D. A. (2016). Detection and interpretation of shared genetic influences on 42 human traits. Nature genetics.

Sakoda, L. C., Jorgenson, E., and Witte, J. S. (2013). Turning of cogs moves forward findings for hormonally mediated cancers. Nat Genet, 45(4): 345–8.

Sivakumaran, S., Agakov, F., Theodoratou, E., Prendergast, J. G., Zgaga, L., Manolio, T., Rudan, I., McKeigue, P., Wilson, J. F., and Campbell, H. (2011). Abundant pleiotropy in human complex diseases and traits. The American Journal of Human Genetics, 89(5): 607–618.

Stephens, M. and Balding, D. J. (2009). Bayesian statistical methods for genetic association studies. Nature Reviews Genetics, 10(10): 681–690.

Thompson, W. K., Wang, Y., Schork, A. J., Witoelar, A., Zuber, V., Xu, S., Werge, T., Holland, D., Andreassen, O. A., Dale, A. M., et al. (2015). An empirical bayes mixture model for effect size distributions in genome-wide association studies. PLoS Genet, 11(12): e1005717.

Vilhjálmsson, B. J., Yang, J., Finucane, H. K., Gusev, A., Lindström, S., Ripke, S., Genovese, G., Loh, P.-R., Bhatia, G., Do, R., et al. (2015). Modeling linkage disequilibrium increases accuracy of polygenic risk scores. The American Journal of Human Genetics, 97(4): 576–592.

Wang, Z., Zhu, B., Zhang, M., Parikh, H., Jia, J., Chung, C. C., Sampson, J. N., Hoskins, J. W., Hutchinson, A., Burdette, L., et al. (2014). Imputation and subset-based association analysis across different cancer types identifies multiple independent risk loci in the tert-clptm1l region on chromosome 5p15. 33. Human molecular genetics, page ddu363.

Wen, X. and Stephens, M. (2014). Bayesian methods for genetic association analysis with heterogeneous subgroups: from meta-analyses to gene-environment interactions. The annals of applied statistics, 8(1): 176.

Zaykin, D. V. and Kozbur, D. O. (2010). P-value based analysis for shared controls design in genome-wide association studies. Genetic epidemiology, 34(7): 725–738.

Zhou, X., Carbonetto, P., and Stephens, M. (2013). Polygenic modeling with bayesian sparse linear mixed models. PLoS Genet, 9(2): e1003264.

Zhu, X., Feng, T., Tayo, B. O., Liang, J., Young, J. H., Franceschini, N., Smith, J. A., Yanek, L. R., Sun, Y. V., Edwards, T. L., et al. (2015). Meta-analysis of correlated traits via summary statistics from gwass with an application in hypertension. The American Journal of Human Genetics, 96(1): 21–36.

Global Lipids Genetics Consortium (2013). Discovery and refinement of loci associated with lipid levels. Nature genetics, 45(11): 1274–1283.

